# A pro-inflammatory effector-memory T cell program links peripheral EBV priming to CNS autoimmunity

**DOI:** 10.64898/2026.09.28.753741

**Authors:** Nikolaj Pagh Kristensen, Thanos Tsaktanis, Sina Frischholz, Jasim K. Benotmane, Lucia Klotz, Philipp Hilgendorf, Myriam Grotz, Mariana V. Sousa, Lasse F. Voss, Stefan Schattgen, Vanessa Lang, Hannah Reimann, Cosima Storr, Roland Lang, Bernd Spriewald, Felix B. Engel, Martin Böttcher, Dimitrios Mougiakakos, Stefanie Schreiber, Dieter H. Heiland, Simone Mader, Paul G. Thomas, Sine Reker Hadrup, Veit Rothhammer, Kilian Schober

## Abstract

T and B cell–mediated immunosurveillance of the central nervous system (CNS) can become maladaptive in multiple sclerosis (MS). Epstein–Barr virus (EBV) is a likely driver of MS, yet how exactly EBV-specific T cells contribute remains unclear. We generated an antigen-resolved atlas of blood and cerebrospinal fluid (CSF) T cells from untreated MS patients and controls. T cell clones carrying a GZMK^+^ effector-memory program in blood were CSF-biased, particularly in MS, and associated with EBV reactivity. In CSF, EBV-reactive GZMK^+^ CD8^+^ T cells engaged B lineage cells. Activated CD8^+^ T cells promoted atypical B cell and plasmablast formation, whereas GZMK-enriched effector-memory cells induced IL-6 and IL-8 release from myeloid cells. Together, these findings identify an infection-imprinted, CNS-biased state and support a model in which EBV-specific CD8^+^ T cells contribute to inflammation through a non-canonical mode of B cell and myeloid cell activation.

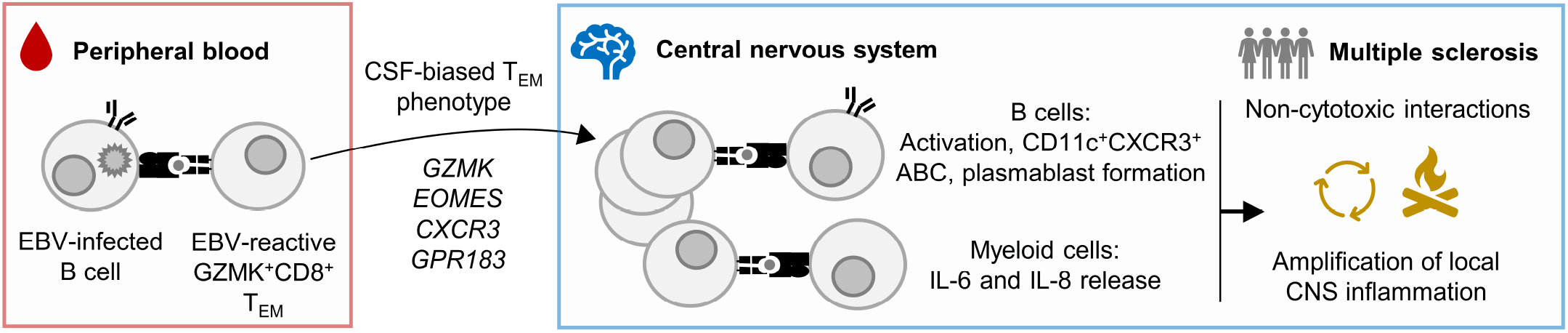

## Main text

The central nervous system (CNS), once considered largely isolated from peripheral immunity, is now recognized as a site of active immune surveillance^1^. Antigen-experienced T cells can access CNS compartments to provide protection against infection, but this capacity may become maladaptive in inflammatory disease.

Multiple sclerosis (MS) is a chronic autoimmune disease of the CNS, affecting 3.1 million individuals worldwide (www.atlasofms.org, September 2026). The most important environmental determinant of MS is Epstein-Barr virus (EBV) infection, which is a necessary but insufficient prerequisite with an MS odds ratio (OR) of 26.5 compared to EBV-seronegative individuals^2,3^. Genetic susceptibility suggests a central role for CD4^+^ T cells, most prominently through HLA-DR15^4^. However, CD8^+^ rather than CD4^+^ T cells constitute a dominant component of inflammatory infiltrates in the cerebrospinal fluid (CSF) and brain parenchyma of MS patients^5–10^. B cells also contribute to disease pathogenesis, as evidenced by intrathecal IgG synthesis and the clinical efficacy of B cell-depleting therapies^11–14^. Despite these advances, how peripheral T cell responses are linked to CNS autoimmunity is poorly understood^15–17^.

Clonal expansion of T cells in the CSF is a hallmark of early MS^9,18,19^. T cell clones can be detected in both peripheral blood (PB) and CSF^10,20–22^. However, it is unclear whether CNS entry of T cells reflects stochastic sampling from the peripheral repertoire, or whether distinct phenotypic profiles, antigen specificities and functional programs predispose individual clones to populate the CSF. Furthermore, the cellular interaction partners of such CNS-biased clones – particularly for CD8^+^ T cells – remain elusive.

EBV is a strong candidate to shape CNS-biased T cell programs. EBV-specific T cells are abundant, long-lived, repeatedly stimulated, and several studies have reported enrichment of EBV-specific T cells in the CSF and CNS tissue of MS patients^5,6,20,22^. EBV establishes latency in B cells, which can act as antigen-presenting cells (APCs), populate CNS inflammatory niches, and provide a plausible substrate for recurrent T cell activation^12–14^. Nonetheless, a mechanistic framework that integrates EBV-specific T cell priming in the periphery with CNS-directed immune responses is lacking, with molecular mimicry between EBV and CNS antigens being more evident for antibodies than for T cells^15,16^. Previous studies have typically addressed clonal expansion, antigen specificity or tissue phenotypes in isolation, and the absence of sufficiently deep, paired PB–CSF datasets has limited the ability to connect these dimensions at single-cell resolution.

Here, we generated a paired PB–CSF single-cell atlas of CD4^+^ and CD8^+^ T cells integrating single-cell RNA sequencing (scRNA-seq), T cell receptor (TCR)-seq, Cellular Indexing of Transcriptomes and Epitopes by Sequencing (CITE-seq) and high-dimensional epitope mapping (jointly referred to as sc-seq), and leveraged transcriptional profiles of physically interacting CSF cells together with published spatial transcriptomic datasets from MS lesions and control brain tissue. We identified a conserved GZMK^+^ effector-memory program in peripheral blood that predicted clonal accumulation in the CSF and was strongly enriched among EBV-specific T cells across CD4^+^ and CD8^+^ compartments, particularly in MS. Within the CNS, GZMK^+^ CD8^+^ T cells preferentially associated with B lineage cells and were found in proximity to myeloid cells. Functionally, GZMK-enriched CD8^+^ T cells promoted atypical B cell^23^ activation and plasmablast formation, as well as IL-6 and IL-8 release by myeloid cells. Overall, these findings support a model in which EBV-specific GZMK^+^ CD8^+^ T cells contribute to a pro-inflammatory circuit during early MS pathogenesis.

### A paired PB and CSF atlas of single T cells

We generated a paired single-cell dataset from PB and CSF of untreated MS patients at diagnosis and non-inflammatory controls (NIC) (**Fig. S1a, Table S1**). The cohort included five MS patients (5/5 female; mean age 31.6 years), five NIC (2/5 female; mean age 36.6 years), and two inflammatory controls (IC; 1x ulcerative colitis and 1x non-MS neuropathy-like condition; 2/2 female; mean age 34.5 years). All MS patients were EBV-seropositive and showed high anti-EBNA IgG titers, consistent with previous observations^24^ (**Fig. S1b**). Three of five MS donors showed markedly higher cell counts in CSF compared to controls (**Fig. S1c**). PB samples were cryopreserved and up to 25,000 CD4^+^ and 25,000 CD8^+^ T cells were sorted per donor for scRNA-seq with paired scTCR-seq and CITE-seq, including hashtag antibodies for donor demultiplexing and surface markers for phenotypic annotation (**Fig. S1a**). CSF samples were processed freshly and without T cell enrichment. In total, the dataset contained 72,343 CD4^+^ and 65,497 CD8^+^ T cells from PB, 36,352 CD4^+^ and 16,535 CD8^+^ T cells from CSF, and 29,281 additional immune cells, including mucosal-associated invariant T (MAIT) cells from PB as well as monocytes, microglia (MG)-like and border-associated macrophage (BAM)-like cells, NK cells, plasmacytoid and conventional dendritic cells (pDC and cDC), B cells, plasmablasts (PBla) and multiplets consisting of T cells interacting with different antigen-presenting cell types from CSF (**Fig. S1d-h**). CSF immune cell type frequencies tended to be higher in MS patients compared to NIC in both absolute and relative numbers (**Fig. S1i**), in line with previous reports^25^.

### GZMK^hi^ IFN^lo^ CD8^+^ T_EM_ cell clones are CSF-biased

Our deep single-cell data enabled us to investigate whether the compartmental distribution of clones between PB and CSF is stochastic or determined by defined phenotypic programs. We first focused on CD8^+^ T cells and identified 23 Leiden clusters based on scRNA-seq and CITE-seq (**Fig. 1a-b, Fig. S2**), spanning naïve (T_N_), central memory (T_CM_), effector memory (T_EM_), effector (T_E_) or effector memory re-expressing CD45RA (T_EMRA_), activated (T_Act_) and CSF-enriched (T_CSF_) states, as well as γδ T cells. T_N_ cells were CD45RA^+^ CD62L^+^ with stem-like gene expression (*BACH2, TCF7*). T_CM_ cells were mostly CD45RA^-^ CD62L^+^. T_EM_ cells were predominantly CD45RA^-^ CD62L^-^ and *GZMK*^hi^. T_E_/T_EMRA_ comprised *GZMB*^hi^ CD45RA^+/-^ CD62L^-^ cells. T_Act_ cells were defined by cell-cycle (*CDKN1B*) and activation markers (*CD69*). T_CSF_ clusters comprised stem-like, *GZMK*^hi^ and tissue-resident (T_RM_)-like populations. Overall, PB CD8^+^ T cells covered the expected differentiation spectrum, whereas T_CSF_ cells were predominantly *GZMK*^hi^ (**Fig. 1a-b, Fig. S2**).

**Figure 1:**
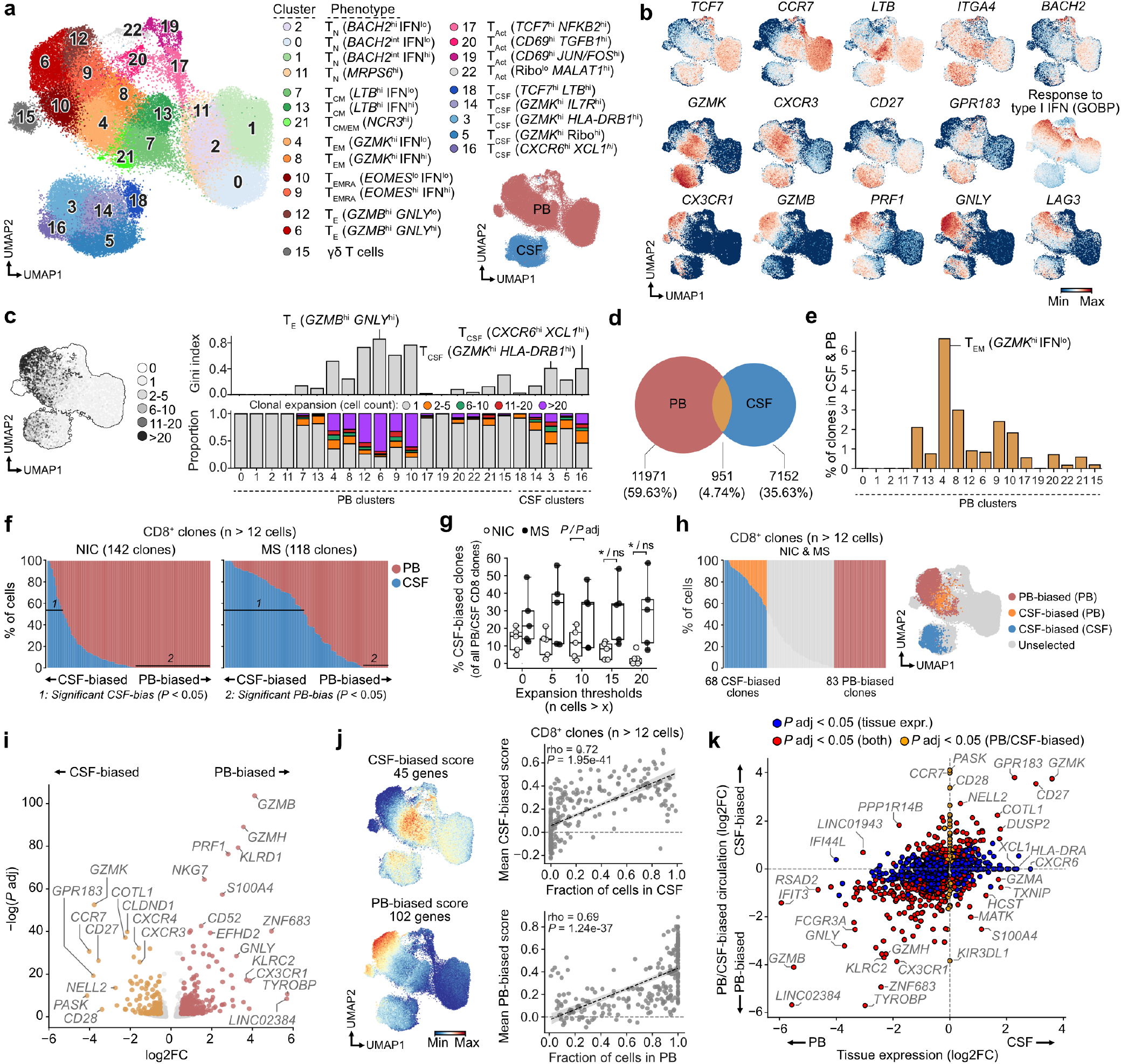
Hyperexpanded CD8^+^ T cell clones in MS are more strongly biased toward the CSF than in NIC. (a) UMAP of 63,769 and 16,252 CD8^+^ T cells from PB and CSF, respectively, with Leiden clusters and compartment annotations. Five NIC, five MS and two IC donors are shown. For brevity, PB origin is not explicitly indicated, whereas CSF-derived populations are labeled T_CSF_. (b) Transcriptomic features related to T cell differentiation, function and phenotype. (c) UMAP of absolute clonal expansion together with side-by-side quantifications of clonal unevenness (Gini index) and proportion of cells in different absolute expansion bins. (d) Venn diagram of CD8^+^ T cell clones shared between PB and CSF. (e) Percentage of CD8^+^ clones shared between CSF and PB among clones in each PB Leiden cluster. (f) Annotation of highly CSF- and PB-biased CD8^+^ T cells. Columns depict hyperexpanded CD8^+^ T cell clones and their relative intraclonal abundance in CSF (blue) or PB (red). A threshold of n > 12 cells was used to define hyperexpanded clones at a fixed statistical threshold (%CSF > 53.8%; see Fig. S3f). (g) Percentage of CSF-biased CD8^+^ T cell clones out of all CD8^+^ clones in PB and CSF across different expansion thresholds. (h) UMAP with highly CSF- and PB-biased CD8^+^ T cells highlighted. (i) Differential gene expression analysis of cells annotated in (h). (j) Gene scores related to the differentially expressed genes found in (i), together with Spearman correlations depicting the positive relationships between CSF- and PB-biased gene scores and the fraction of cells in CSF and PB, respectively, among hyperexpanded CD8^+^ T cell clones. (k) XY graph comparing the log2 fold change values of CSF- and PB-biased gene sets with log2 fold change values of genes upregulated among CSF CD8^+^ T cells compared to PB CD8^+^ T cells. Statistical thresholds to annotate CSF-biased CD8^+^ T cells in (f) and (g) were computed by permutation of a binomial distribution using the relative proportions of total CSF and PB cells as estimates of the expected frequencies. *P* values were calculated using a Wilcoxon test and adjusted using the Benjamini-Hochberg procedure. *, **, ***, **** correspond to *P* < 0.05, *P* < 0.01, *P* < 0.001, and *P* < 0.0001, respectively.

We next harnessed scTCR-seq information to assess the clonal architecture across compartments. T_CSF_ cells showed more evenly distributed clonotype frequencies than PB T_E_/T_EMRA_ cells, although CSF did contain many expanded clones (**Fig. 1c**). Most clones were compartment-restricted, with only 4.74% being shared (**Fig. 1d**). Shared clones were highly enriched in the *GZMK*^hi^ IFN^lo^ T_EM_ C4 in PB (**Fig. 1e**) and the *GZMK*^hi^ *HLA-DRB1*^hi^ C3 in CSF (**Fig. S3a**).

MS patients had more CD8^+^ T cells in CSF compared to NIC both in relative and absolute numbers (**Fig. S3b-c**). Furthermore, sister cells belonging to PB/CSF-shared clonotypes were more commonly found occupying the CSF in MS patients compared to NIC, which was particularly evident for expanded T cell clonotypes (**Fig. S3d**). However, individual clones could be biased to either the PB or the CSF compartment (**Fig. S3e**), arguing against ‘passive spillover’ from PB to CSF. To quantify compartmental skewing in a systematic manner, we focused on hyperexpanded clones (>12 cells), where compartmentalization can be assessed with sufficient statistical power. Here, a CSF fraction of 0% (‘PB-biased’) or >53.8% (‘CSF-biased’) deviated statistically significantly from random expectation (**Fig. S3f**). Expanded clones from MS patients were more CSF-biased than clones from NIC, which also held true when alternative thresholds were applied to define clonal expansion (**Fig. 1f-g**). Re-analysis of published data^10^ supported this pattern, with donors showing high CSF content predominantly among relapsing MS patients (**Fig. S3g-i**). In summary, MS patients possessed expanded CD8^+^ T cell clones with a disproportionately large fraction of cells found in CSF.

We hypothesized that clonal compartmentalization was associated with distinct phenotypes in PB. To investigate this, we focused on clones with at least one cell in PB and compared PB cells from PB-biased with PB cells from CSF-biased clones (**Fig. 1h**). PB-biased clones were characterized by a terminal effector signature (*GZMB, GNLY, CX3CR1*) (**Fig. 1i**). PB-biased clones were also enriched for *ZNF683* (Hobit), which is associated with T_RM_ cells in human tissues but more highly expressed by peripheral effector cells^26,27^ (**Fig. 1i, Fig. S2c-d**). In contrast, CSF-biased clones displayed an effector memory program marked by *GZMK* together with *CCR7, CD27* or *CD28* (**Fig. 1i**). The CSF-biased program also included *CXCR3* and *GPR183*, both implicated in CNS-homing^12,28^.

We then derived gene scores from PB/CSF signatures and applied them to all cells. The CSF-biased score was enriched in *GZMK*^hi^ T_EM_ and T_CM_ subsets in PB (C8, C4, C17, C7) (**Fig. S3j-k**). Importantly, higher CSF-biased and PB-biased scores in PB were strongly correlated with increased representation of the same clones in the CSF and PB, respectively (**Fig. 1j**). Overall, these findings link distinct peripheral phenotypes to T cell accumulation in the CNS.

Finally, we asked to what extent the peripheral CSF-biased program simply reflected the transcriptional changes accompanying tissue entry. As expected, PB-biased genes were generally more highly expressed in PB than in CSF, whereas CSF-biased genes were enriched in CSF (**Fig. 1k, Fig. S3l**). Some genes deviated from this pattern: for example, *CCR7* or *CD28* were enriched in PB cells of CSF-biased clones compared to PB cells of PB-biased clones, but not globally higher in CSF compared to PB (**Fig. 1k, Fig. S3m)**. Conversely, *GZMA* or *CXCR6* were enriched in CSF compared to PB, but not part of the CSF-biased program in PB (**Fig. 1k**).

Overall, clonal expansion of CD8^+^ T cells in the CSF was non-stochastic and instead predicted by a *GZMK*^hi^ T_EM_-like program already present in PB, indicating that defined peripheral T cell states predispose clones to CNS accumulation.

### EBV-specific CD8^+^ T cells are enriched for the CSF-biased transcriptional program

We next asked which antigens are recognized by CSF-biased T cell clones. To answer this question, we profiled virus-specific CD8^+^ T cells using highly multiplexed DNA-barcoded peptide-human leukocyte antigen (pHLA) multimers covering 113 epitopes across nine HLA-I heavy chains and ten viruses (**Fig. 2a**). PB samples from the same donors as in Fig. 1 were sampled to generate a separate multimer dataset, in which 84,710 pHLA^+^ CD8^+^ T cells were sorted and subjected to scRNA-seq with scTCR-seq and CITE-seq including a pHLA DNA barcode readout (**Fig. 2a, Fig. S4a**).

**Figure 2:**
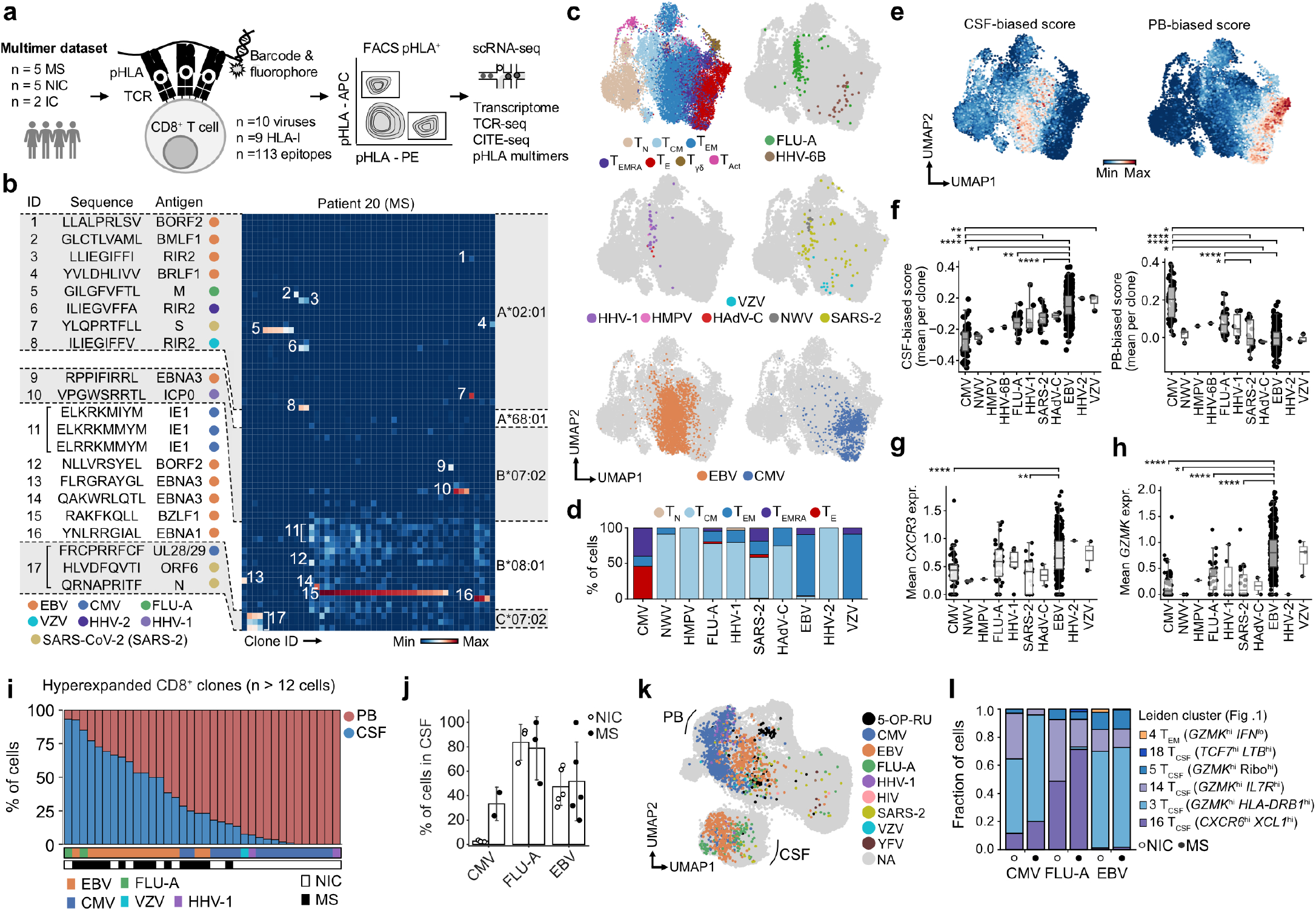
EBV-specific CD8^+^ T cells recapitulate the CSF-biased program and are detected in the CSF as GZMK^hi^ CD8^+^ T cells in both MS and NIC. (a) Experimental setup deploying a large panel of pHLA multimers for the discovery and characterization of antigen-specific CD8^+^ T cell clones in the same cohort of donors as Fig. 1. (b) Example heatmap for single-cell annotation of antigen specificities. Individual cells are grouped and summarized per clonotype. (c) UMAP of phenotypes and antigen specificities after integration and label transfer by scANVI. Major PB CD8^+^ T cell cluster annotations depicted in Fig. 1a were used as the reference for label transfer. (d) Stacked bar graphs showing the phenotypic distribution of different virus specificities. (e) UMAP depicting CSF-biased and PB-biased gene scores. (f-h) Transcriptomic summary statistics per clonotype grouped by virus specificity. See Fig. S4b for clone (n) numbers per condition, i.e., number of clones by virus specificity. (f) Boxplots showing the mean CSF-biased score per clone and the mean PB-biased score per clone of different virus specificities. (g) Mean normalized *CXCR3* gene expression per clone. (h) Mean normalized *GZMK* gene expression per clone. (i) Antigen specificities of hyperexpanded CD8^+^ T cell clones originally depicted in the paired PB/CSF dataset in Fig. 1. Antigen annotations were expanded using computational methods; see Methods. (j) Column plots depicting the frequency of cells in CSF among antigen-specific clones discovered in the paired PB/CSF dataset depicted in Fig. 1. (k) UMAP (Fig. 1) depicting all annotated antigen specificities. (l) Stacked bar graphs detailing the phenotypic distribution in CSF of antigen-specific CD8^+^ T cells. Dots represent donors. Antigen annotations were based solely on multimer binding in panels (c-h) and were extended using computational methods in panels (i-l). *P* values were calculated using Dunn’s test and adjusted using the Benjamini-Hochberg procedure. *, **, ***, **** correspond to *P* < 0.05, *P* < 0.01, *P* < 0.001, and *P* < 0.0001, respectively.

Overall, we detected 3,973 epitope-specific T cells and 443 clones across 83 unique antigen-specific immune responses observed for 47 epitopes including particularly large numbers of clones recognizing the lytic EBV antigen BZLF1 (**Fig. S4b-d**). Some clones bound multiple targets, which represented biologically valid cross-reactivity, exemplified by a clone from patient 20 recognizing previously described^29^ homologous HLA-A*02:01 restricted epitopes from EBV, HHV-2 and VZV (**Fig. 2b**). Detection of EBV- and Cytomegalovirus (CMV)-specific T cell responses was consistent with serological testing from the same donors, except for HLA-C*07:02/FRC (CMV), which bound to clones also in CMV-seronegative donors. HLA-C*07:02 is known to bind to killer-cell immunoglobulin-like receptors (KIR)-expressing effector T cells in an epitope-independent manner^30^. We therefore excluded HLA-C*07:02 responses from downstream analyses.

We then asked how antigen specificity relates to phenotypic programs in PB. Mapping phenotypic labels from the original dataset (Fig. 1) onto the multimer dataset revealed that SARS-CoV-2 (SARS-2)- and influenza-A (FLU)-specific T cells predominantly occupied T_CM_ states, whereas CMV-specific cells were enriched among T_E_ cells (**Fig. 2c-d, Fig. S4e-g**). In contrast, EBV- and Varicella Zoster virus (VZV)-specific T cells were highly enriched among T_EM_ cells (**Fig. 2c-d, Fig. S4e-g**). On a cluster level, 86% of EBV-specific cells mapped to one of the two *GZMK*^hi^ T_EM_ clusters. 47.3% of EBV-specific cells thereby adopted the *GZMK*^hi^ IFN^hi^ T_EM_ phenotype of C8, substantially more than other virus-specificities (<6%) (**Fig. S4e**).

EBV-specific clones showed highly elevated CSF-biased scores compared to other viral specificities, whereas PB-biased scores were enriched in CMV-specific clones (**Fig. 2e-f**). EBV-specific clones also expressed high levels of *CXCR3* and *GZMK*, core components of the CSF-biased signature (**Fig. 2g-h**). Among EBV-specific T cells, clones targeting latency-associated antigens (EBNA1, EBNA3) showed the highest CSF-biased scores and the highest expression of *CXCR3* and *GZMK* (**Fig. S4h-i**). Thus, while multiple viral specificities can adopt CSF-biased phenotypes, EBV-specific T cells form the most dominant compartment with enrichment of this program.

In our original sc-seq dataset of PB/CSF T cells (Fig. 1), we ‘rediscovered’ 182 clones with mapped antigen specificity through our multimer-annotated sc-seq dataset (Fig. 2), including 145 EBV-, 20 CMV- and 8 FLU-specific clones. Notably, this included 95 clones (64 for EBV, 16 for CMV, 8 for FLU) that we also identified in CSF. To enhance antigen specificity profiling in our original dataset further, we followed a computational approach and integrated predicted epitope-specific TCRs from a commercial tool (ImmuneWatch DETECT), from a public TCR database (VDJdb^31^) and from ‘clumps’ of EBV-specific TCRs that share phenotypic and clonotypic features^32^. This led to a total of 536 antigen-mapped clones in our initial dataset (**Fig. S5a**) allowing us to directly annotate the originally observed hyperexpanded clones to validate their suspected EBV-specificity. CSF-biased, hyperexpanded clones (n>12 cells, Fig. 1), were frequently EBV-specific, whereas PB-biased clonotypes recognized CMV (**Fig. 2i, Fig. S5b-c**). EBV-specific CD8^+^ T cells showed extensive CSF representation in both MS and NIC, with an average of 47.2% and 51.7% of cells located in the CSF, while CMV-specific cells had a CSF fraction of only 2.8% (NIC) to 33.2% (MS) (**Fig. 2j**). Interestingly, antigen-mapped FLU-specific CD8^+^ T cells showed an even higher CSF localization than EBV clones (83.6% in NIC, 78.7% in MS) (**Fig. 2j**) likely due to their tendency to adopt T_RM_-like features in CSF which EBV-specific CD8 T cells did not (**Fig. 2k-l**).

To further validate EBV specificity, we used an orthogonal approach through mapping TCRs expanding in cultures with autologous EBV-transformed B-lymphoblastoid cell lines (B-LCL)^6^ to our single-cell dataset (**Fig. S5d**). CD8^+^ T cell clones previously identified by pHLA multimers preferentially expanded in B-LCL cultures (**Fig. S5e**). B-LCL-reactive CD8^+^ clones were enriched among CSF-biased clones in both MS and NIC (**Fig. S5f-g**). The combination of epitope assignment through multimers (mostly EBV specificity) and B-LCL-reactivity explained almost 50% of reactivities of CSF-biased clones in MS (**Fig. S5h**). Thus, CNS-homing is an intrinsic feature of EBV-specific clones that is pronounced in MS, but also observed in NIC.

In CSF, EBV-specific T cells were found especially among *GZMK*^hi^ *HLA-DRB1*^hi^ cells (C3 in our original dataset from Fig. 1) (**Fig. 2k-l**), possibly indicating local T cell activation. CMV-specific T cells were also found in these clusters while FLU-specific cells were additionally prominent in the *CXCR6*^hi^ *XCL*1^hi^ T_RM_-like C16 (**Fig. 2k-l**).

Together, these findings demonstrate that the characteristic trait of EBV-specific CD8^+^ T cells is not CNS localization *per se*, but their convergence on a CSF-biased effector-memory program in PB, an activated phenotype within the CNS, as well as local clonal expansion.

### The CNS-homing transcriptional program is conserved in EBV-reactive CD4^+^ T cells

We next asked whether this EBV antigen-linked, CSF-biased transcriptional program is conserved in CD4^+^ T cells. We identified 21 CD4^+^ T cell Leiden clusters spanning T_N_, T_CM_, T_EM_, cytolytic (T_C_), T_Act_, regulatory (T_Reg_) and T_CSF_ states (**Fig. S6a-e**).

In PB, T_CM_/T_EM_ clusters were enriched for *GZMK, CXCR3* and *ITGA4*, whereas T_C_ cells expressed *PRF1* and *GZMB* and represented the most clonal population (**Fig. S6e-f**). In CSF, CD4^+^ T cells comprised stem-like *TCF7*^*hi*^ states, activated and *CXCR3*^hi^ populations (**Fig. S6a-e**). Among these, *CXCR3*^hi^ *GZMK*^hi^ *EOMES*^hi^ *FOXP3*^lo^ cells lacked cytotoxic molecules, consistent with a regulatory type 1 (Tr1) phenotype^33,34^. Activated cells (C1) and Tr1 cells (C12) represented the most expanded and clonal populations in CSF (**Fig. S6f**).

1.9% of non-naïve CD4^+^ T cell clones were shared between PB and CSF (**Fig. S6g**). Like previously observed for CD8^+^ T cells, CD4^+^ T cell clones were highly compartmentalized (**Fig. S6h**). Among PB clusters, T_CM/EM_ C3 (*PASK*^int^ *FTH1*^hi^) showed the greatest clonal overlap with CSF clones, followed by T_EM_ C9 (*TNF*^hi^ *TNFRSF4*^hi^) (**Fig. S6i**), which were characterized by high *CXCR3* expression (**Fig. S6e**). CD4^+^ clones showed increased CSF representation in MS patients compared to controls (**Fig. S6j–l**), consistent with CD8^+^ clones.

To assess phenotypic PB signatures of CSF- or PB-biased CD4^+^ T cells, we again focused on significantly compartmentalized clones, with >83.3% of sister cells in CSF across different expansion thresholds. Because CD4^+^ clones are generally less expanded than CD8^+^ clones, a lower threshold of five cells was used to allow statistical assessment of both PB- and CSF-biased populations (**Fig. S7a-b**). MS patients contained more expanded CD4^+^ clones with high CSF representation than NIC (**Fig. S7b-c**). Next, we used CSF-biased and PB-biased clones from PB to define CSF- and PB-biased gene scores (**Fig. S7d-e**). As for CD8^+^ clones, the CSF-biased score expression in PB CD4^+^ clones correlated with their representation in CSF (**Fig. S7f**). Consistently, the CSF-biased program was highly conserved across CD4^+^ and CD8^+^ T cells, with *GZMK, CCR7* and *CD27* as top genes (**Fig. S7g**). While the PB-biased score was highly enriched in T_C_ C18, the CSF-biased score in PB was dominant in T_CM/EM_ C3 (**Fig. S8a**), which had also shown the highest clonal overlap with CSF (Fig. S6i). Thus, the CSF-biased transcriptional program identified in CD8^+^ T cells is conserved in CD4^+^ T cells and similarly predicts clonal accumulation in the CSF.

To assess antigen specificity, particularly EBV reactivity, we mapped CD4^+^ T cell clones that were reactive to autologous EBV-transformed B-LCL to our single-cell dataset (Fig. S5). Four B-LCL-reactive CD4^+^ clones were found among hyperexpanded CD4^+^ clones. The two B-LCL-reactive clones found among MS patients were both CSF-biased and located exclusively in CSF (**Fig. S8b**). Among all clones, B-LCL-reactive CD4^+^ clones were enriched among the CSF *CXCR3*^hi^ *GZMK*^hi^ C12 and *CXCR3*^hi^ *KLRB1*^hi^ C11 cells (**Fig. S8c**) and were enriched for a Tr1/Th1 phenotype (**Fig. S8d**).

Overall, EBV-reactive CD4^+^ T cells displayed a *CXCR3*^hi^ Tr1/Th1-like phenotype in the CSF with a capacity to become CSF-biased in both MS and NIC donors consistent with our observation of EBV-specific, CSF-biased CD8^+^ T cells.

### EBV-specific GZMK^+^ CD8^+^ T cells preferentially interact with B lineage cells in the CSF

Having defined the phenotype and antigen specificity of CSF-biased T cells, we next asked which cellular partners they encounter within the CNS compartment. In the CSF, we captured diverse immune cell populations along with numerous cellular multiplets, comprising T cells (marked by fully recombined TCRs), B cells, PBla, cDC, pDC, NK cells, monocytes or MG/BAM-like cells (**Fig. 3a, Fig. S9a-c**). We considered that CSF multiplets may reflect serendipitously captured yet physiologically relevant cellular interactions^35,36^. In support of this, we found that T cell-containing multiplets were enriched for professional APC partners compared to other cell types such as NK cells (**Fig. S9d)**. Furthermore, we observed that T cell-containing multiplets far outnumbered multiplets without T cells, i.e., APC–APC multiplets (**Fig. S9e-f)**. T–APC multiplets tended to be more frequent in MS patients compared to NIC (**Fig. S9g**). To evaluate our findings at the protein level we used imaging flow cytometry, which confirmed the presence of rare CD3^+^ CD19^+^ and CD3^+^ CD14^+^ multiplets in CSF of MS patients (**Fig. S10a-e**). Additionally, we observed a consistently increased frequency of T cell-containing multiplets in PB of MS patients compared to NIC (**Fig. S10f-j)** suggestive of ongoing immune cell perturbation in MS patients. Collectively, these findings indicate that CSF multiplets captured distinct physically interacting immune cells of relevance to MS.

**Figure 3:**
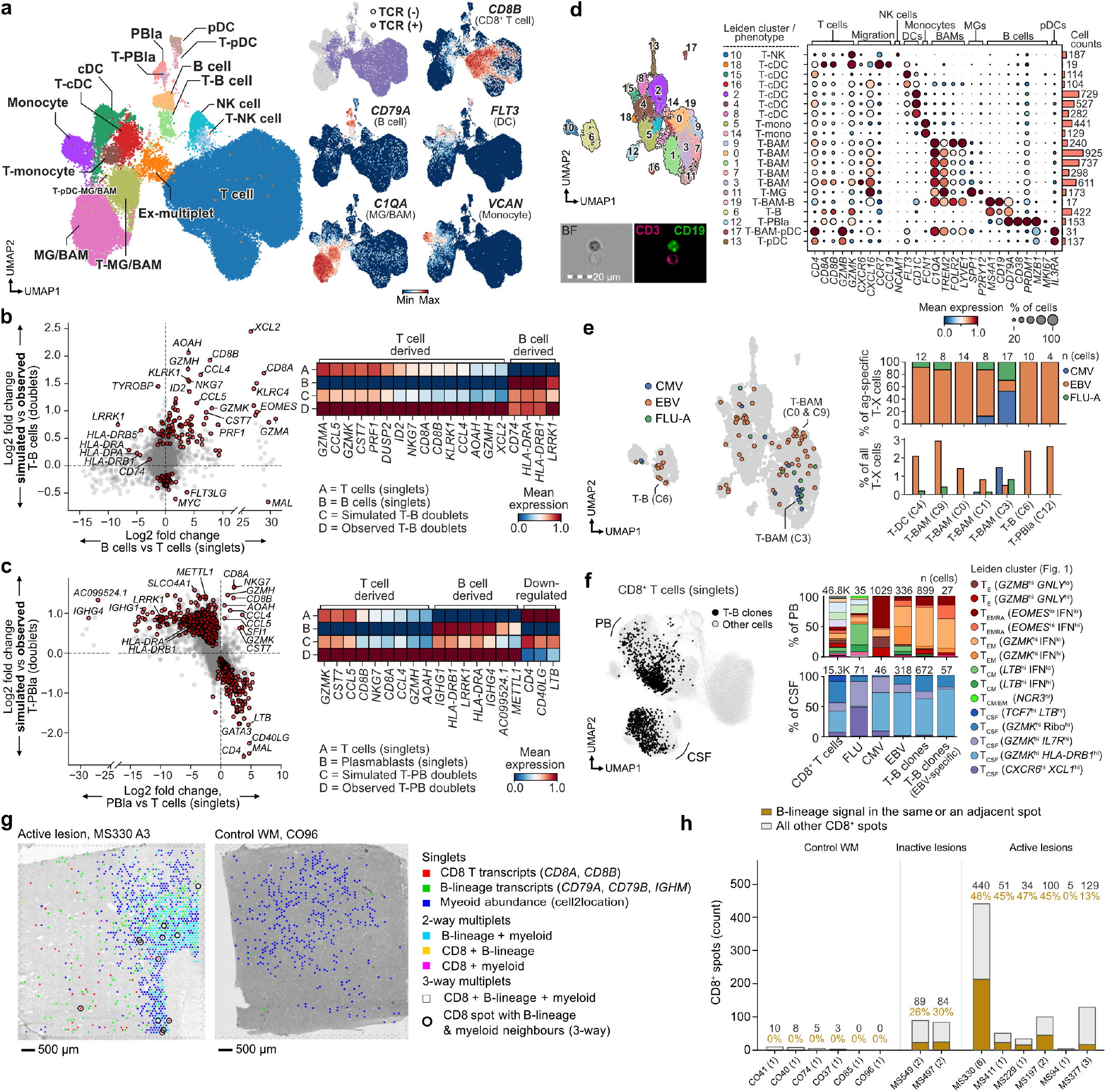
EBV-specific GZMK^+^ CD8^+^ T cells preferentially interact with B lineage cells in the CSF. (a) UMAP of subclusters of CSF immune cells alongside manual cluster annotations with features and marker genes characterizing individual cell subsets. (b) Comparison of differentially expressed genes. T and B cell-specific expression (x-axis) versus differential gene expression in observed multiplets compared with simulated doublets mixed from singlet parent populations (y-axis). Red indicates differentially expressed genes on both axes. A matrix plot accompanies each plot as a summary of noteworthy differentially expressed genes. (c) As in (b), but for PBla and T cells, comparing observed T–PBla with simulated T–PBla on the y-axis. (d) UMAP and gene expression summary of observed T cell-containing multiplets. The inset shows an example of CSF T–B interactions identified by imaging flow cytometry also shown in Fig. S10. (e) Antigen specificity of clones involved in T–APC interactions. Antigen specificities were derived from pHLA multimers and computational mapping of matching clones to TCR databases as detailed in Fig. 2. (f) UMAP and phenotypic distribution (Fig. 1) highlighting singlet cells belonging to clones interacting with B cells in CSF. Only donors with detectable B cells or T–B multiplets in CSF are represented (four MS patients and five NICs). (g) Side-by-side Visium sections from active lesions in postmortem MS patients and control white matter (WM) in postmortem healthy controls without neuropathological changes. Grayscale represents the H&E image. Positive spots per section: CD8 52 and 0; B-lineage 510 and 3; myeloid 879 and 421; ringed 12 and 0. (h) Number of CD8^+^ spots per donor. X-axis indicates different donors. 24 Visium sections were analyzed across inactive and active lesions in MS patients and control WM. Numbers in parentheses indicate the number of sections for each donor. Colored bars and fonts represent the proportion of CD8^+^ spots co-occurring or adjacent to B cell transcript-positive spots. Data in (g-h) originate from Lerma-Martin et al. 2024 and Schroeder et al. 2025.

We hypothesized that interacting T cells and APCs represent only a narrow subset of the T cell and APC phenotypes present in the CSF. To investigate the particular phenotype of T cells interacting with each subset of APCs, we compared outputs from two differential gene expression analyses (**Fig. S11a**): First, we identified differentially expressed genes (DEGs) between parent singlet populations to map T cell and APC-derived genes. We then compared gene expression profiles of observed doublets with simulated doublets that were generated by computational mixing of singlet profiles to identify the genes characteristic of T cells and APCs interacting with each other. This approach enabled us to investigate which T cell or APC-derived genes are enriched in physically observed multiplets.

We first focused on T–B interactions because B cells and PBla are central to MS pathogenesis and may harbor latent EBV episomes, EBV antigens, or EBV-homologous CNS antigens^12–14,37,38^. Compared with simulated doublets, observed T–B interactions were enriched for both *CD8A* and *CD8B* but not for *CD4*, suggesting that CD8^+^ T cells, rather than CD4^+^ T cells, interact with B cells in the CSF (**Fig. 3b**). Furthermore, we observed key genes of CSF-biased T cell clones enriched among T–B multiplets, including *GZMK* and *EOMES*. Genes of *NKG7, PRF1*, and additional non-cytotoxic granzymes like *GZMH* and *GZMA* were also enriched among observed T–B multiplets while cytotoxic *GZMB* and *GNLY* were not enriched (**Fig. 3b**). Among the few enriched B cell-specific genes, *LRRK1* is linked to B cell activation and antibody production^39^, while HLA-DR enrichment indicated enhanced antigen presentation (**Fig. 3b**).

Observed T–PBla multiplets were also enriched for *LRRK1* and HLA-DR genes on the PBla side, and *GZMK, PRF1, GZMH, NKG7, CD8A*, and *CD8B* on the T cell side, compared to simulated T–PBla multiplets (**Fig. 3c**). Furthermore, we observed significantly fewer *CD4* and *CD40LG* transcripts than expected from simulated T–PBla multiplets, indicating that also PBla primarily interact with CD8^+^ T cells in the CSF (**Fig. 3c**). T–PBla were furthermore enriched for multiple additional B cell-specific genes including *IGHG1* and *IGHG4*, as well as *METTL1* compared to simulated T– PBla multiplets. *METTL1* is upregulated in recently activated, post-immunization B cells and associated with multiple germinal center processes including class switching of memory B cells^40^.

DEGs in observed T–B and T–PBla compared to simulated multiplets were not associated with apoptosis in pathway analyses (**Fig. S11b-e**). Instead, in T–B multiplets we observed an enrichment of pathways related to antigen-receptor signaling and activation, and in T–PBla multiplets an enrichment of pathways related to glycoprotein biosynthesis, protein transport, response to type I interferon, and metabolism (**Fig. S11b-e**).

We also observed multiplets consisting of T and myeloid cells, which were less consistently enriched for CD8^+^ T cells compared to interactions involving B lineage cells. Observed T–MG/BAM-like multiplets were enriched for *GZMA, GZMK, GZMH, CCL5, NKG7, EOMES* as well as *CXCL16* and *CXCR6* (**Fig. S12a**). Notably, *CXCL16* was expressed by MG/BAMs, whereas *CXCR6* was expressed by the specific subset of T cells interacting with MG/BAMs, highlighting a greater participation of *CXCR6*^hi^ T_RM_ cells and the CXCR6–CXCL16 axis in T– MG/BAM-like interactions in the CSF. Pathways enriched among T–MG/BAM-like multiplets were related to microglial cell activation and antigen presentation on HLA class II (**Fig. S12b-c**).

Overall, these results suggest that B cells and plasmablasts preferentially interacted with CD8^+^ *GZMK*^hi^ T cells in the CSF and that these interactions were associated with pro-inflammatory activation rather than cytotoxic killing.

To resolve the heterogeneity among interacting T– APCs, we subclustered all multiplets into 20 T–APC clusters (**Fig. 3d**). CD4^+^ T cells interacted with many APCs including cDCs (e.g., C15), MGs (C11), and pDCs (C13), whereas CD8^+^ T cells were specifically enriched in T–B (C6), T–PBla (C12), and individual cDC (C18) and BAM-like (C3) multiplets. *GZMB* was restricted to pDC multiplets, consistent with its expression in pDCs^41^, whereas high expression of *GZMK* generally marked CD8-dominant multiplets. *CXCR6* expression was specific to T–BAM C3, indicating that the above-described interaction between *CXCR6*^hi^ T_RM_-like cells and target *CXCL16*^hi^ BAM cells constituted a smaller subset of all T–MG/BAM multiplets.

Mapping of identified antigen-specificities onto our reclustered T–APC multiplets yielded dozens of EBV-, CMV- and FLU-specific cells (**Fig. 3e**). Strikingly, among cells with known specificity, T–B and T–PBla multiplets contained 100% EBV-specific T cells (10/10 and 4/4 cells, respectively). T–BAM multiplets, on the other hand, contained a mixture of antigen-specificities, from which CMV- and FLU-specific CD8_+_ T cells were most enriched among the *CXCR6*^hi^ T–BAMs (C3) (**Fig. 3e**). This was consistent with the enrichment of FLU-specific cells among *CXCR6*^hi^ T_RM_-like singlet cells in the CSF (Fig. 2l).

Finally, we examined the phenotypes of CSF and PB singlets belonging to clones that interacted with APCs in CSF. As singlets in CSF, T cell clones identified in T– B and T–PBla multiplets were enriched for *GZMK*^hi^ cells, and most closely resembled CMV- and EBV-specific cells (**Fig. 3f, Fig. S13a**). In PB, the same clonotypes were enriched for *GZMK*^hi^ T_EM_ phenotypes, corresponding to EBV-specific cells (**Fig. 3f, Fig. S13a**).

Cells in PB and CSF belonging to T–MG/BAM clones adopted both *GZMK*^hi^ and T_RM_-like phenotypes, consistent with their respective antigen-specificities (**Fig. S13b**). Interestingly, 62% (60/97 cells) of all observed FLU-specific singlets in CSF were of the *CXCR6*^hi^ T_RM_ phenotype, but among FLU-specific clones interacting with MG/BAMs this number rose to 100% (32/32 cells), supporting the role of the CXCR6– CXCL16 axis in shaping this interaction (**Fig. S13b)**.

We finally investigated whether CD8^+^ T cells interact with B cells also in the brain parenchyma. To this end, we re-analyzed published spatial transcriptomics data of MS patients^42,43^. Active lesions contained more CD8^+^ T cells than control white matter (WM) (**Fig. 3g-h, Fig. S14a-d**). CD8^+^ T cells in active lesions had a B lineage cell in their spot neighborhood more often than other lymphocyte subsets from the same donor, a comparison that was descriptive only in inactive lesions (two donors) and not estimable in control WM, which had too few CD8^+^ spots per section to test (**Fig. 3h, Fig. S14e**). In addition, *GZMK* represented the leading granzyme in lesions (**Fig. S14f-h**).

In summary, T–B and T–PBla interactions in the CSF were highly enriched for EBV-specific *GZMK*^hi^ CD8^+^ T cells, whereas T–MG/BAM interactions involved more diverse antigen specificities, with FLU-specific CD8^+^ T cells highlighting a particular role of the CXCR6– CXCL16 axis. In addition, active CNS lesions of MS patients were enriched for CD8^+^ T–B interactions and *GZMK* expression.

### GZMK^+^ CD8^+^ T_EM_ cells promote atypical B cell activation and plasmablast formation

Our findings of CD8^+^ T–B and T–BAM-like interactions prompted us to investigate the functional consequences of the GZMK^+^ effector-memory program in these interactions. To be able to sort T_EM_ cells with high intracellular GZMK expression, we used CCR5 as a surface marker^44^. GZMK was highly enriched in CCR5^+^ T_EM_ cells from healthy-donor PB, which in turn expressed considerably less GZMB than CCR5^-^ T_EM_ or T_E_ cells (**Fig. 4a-b**).

**Figure 4:**
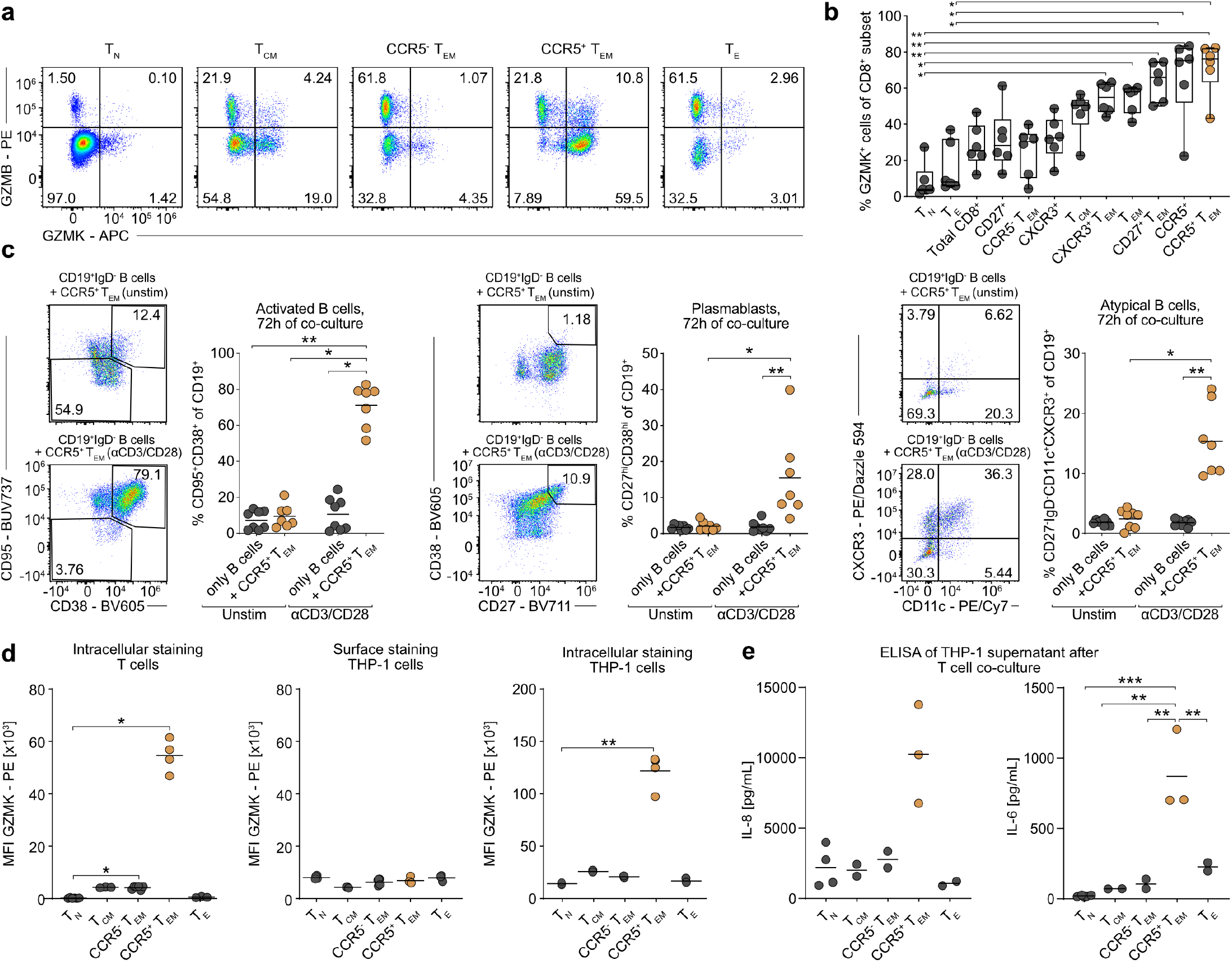
GZMK^+^ T_EM_ cells influence the pro-inflammatory activity and activation state of target cells. (a) Representative flow cytometry data from one donor, showing GZMK and GZMB expression in CD8^+^ T cell subsets directly *ex vivo*. (b) GZMK expression in CD8^+^ T cell subsets. Dots represent independent donors (n = 6). (c) Representative flow cytometry plots and quantification of B cell activation, plasmablast formation and atypical B cell formation after 72 h co-culture of sorted CD19^+^IgD^−^ B cells with unstimulated or αCD3/CD28-activated sorted CD8^+^ T cells. Dots represent 1-2 technical replicates from four donors across two independent experiments. (d) Detection of intracellular GZMK in T cells, and surface and intracellular GZMK in THP-1 cells, after 4 h of co-culture of sorted CD8^+^ T cell subsets with THP-1 cells from one experiment. Dots represent 1-2 technical replicates from two donors. (e) IL-6 and IL-8 detected in supernatants after 48 h of co-culture of sorted CD8^+^ T cell subsets with THP-1 cells measured by ELISA. Dots represent technical replicates from one donor; representative of three experiments. *P* values were calculated using Dunn’s test and adjusted using the Benjamini-Hochberg procedure. *, **, ***, **** correspond to *P* < 0.05, *P* < 0.01, *P* < 0.001, and *P* < 0.0001, respectively.

Next, we tested whether CD8^+^ T cells, like CD4^+^ T cells, could influence B cell maturation and activity. We co-cultured sorted CCR5^+^ or CCR5^−^ T_EM_ and T_E_ cells with IgD^−^ antigen-experienced B cells for up to 72 hours, with or without αCD3/αCD28 stimulation (**Fig. 4c, Fig. S15a**). Activated CD8^+^ T cells formed T–B doublets *in vitro* and induced B cell activation (CD38^+^CD95^+^), plasmablast formation (CD27^hi^ CD38^hi^) and a CD11c^+^ CXCR3^+^ atypical B cell-like phenotype^23^ in the absence of exogenous IL-21 (**Fig. 4c, Fig. S15b-d**). Although all tested CD8^+^ T cell subsets could induce these B cell responses *in vitro* (**Fig. S15d**), *GZMK*^hi^ T_EM_ cells were the subset preferentially interacting with B cells *in situ* (Fig. 3). Thus, this broadly shared functional capacity may become biologically relevant specifically through the localization of CCR5^+^ T_EM_ cells within T–B interactions. At the same time, the fact that CCR5^−^ T_EM_ cells could also promote B cell activity argues against GZMK itself being the direct mediator of these effects.

GZMK has previously been linked to transendothelial diapedesis^44^. However, T cell-specific knockout of *Gzmk* did not impair CNS migration in a rat model of experimental autoimmune encephalitis (EAE)^28^, whereas *Cxcr3* and *Gpr183* – two components of our CSF-biased program – emerged as migration-relevant molecules (**Fig. S16a**), consistent with their established role in T cell tissue migration^45,46^. We therefore examined potential local effector functions of GZMK after CNS entry. Recombinant GZMK induced opsonization of target cells (THP-1, HUVECs and HEK293T). However, in contrast to recent reports^47,48^, this did not efficiently progress to terminal complement complex formation (**Fig. S16b-c**).

We next turned to a well-established extracellular inflammatory function of GZMK: activation of protease-activated receptor (PAR)-dependent responses in macrophages, endothelial and epithelial cells^49,50^. We co-cultured sorted CD8^+^ T cell subsets with THP-1 cells, quantified intra- and extracellular GZMK by flow cytometry, and measured IL-6 and IL-8 release (**Fig. 4d-e, Fig. S16d-g**). In contrast to recombinant GZMK, sorted GZMK^hi^ CCR5^+^ CD8^+^ T_EM_ cells only led to intracellular GZMK accumulation within THP-1 cells (**Fig. 4d, Fig. S16d-f**). A lack of target cell death (**Fig. S16g**) was thereby consistent with the non-cytotoxic phenotype of *GZMK*^hi^ T_EM_ cells *in situ* (Fig. 3). Instead, GZMK-enriched CCR5^+^ CD8^+^ T_EM_ cells, but not CCR5^-^ T_EM_ or T_E_ cells, selectively induced IL-6 and IL-8 production by THP-1 cells (**Fig. 4e**).

Together, activated CD8^+^ T cells, including GZMK^+^ T_EM_ cells, could promote atypical B cell and plasmablast formation, and GZMK^+^ CD8^+^ T_EM_ cells selectively induced IL-6 and IL-8 release by myeloid cells.

## Discussion

EBV is the strongest environmental risk factor for MS, yet how peripheral antiviral immunity leads to CNS autoimmunity remains unresolved. Our data support a model in which EBV-specific T cells acquire a GZMK^+^ T_EM_ differentiation state that accumulates in the CSF and contributes to local inflammation through interactions with B and myeloid cells. Our findings add an orthogonal layer to the relationship between EBV and MS: while previous work has focused on TCR recognition of CNS antigens^14,51^, our data suggest that the differentiation state and tissue tropism of EBV-specific T cells are important determinants of CNS autoimmunity. This model remains fully compatible with molecular mimicry and epitope spreading, including antibody-mediated mechanisms for which cross-reactivity is firmly established between the EBV antigen EBNA1 and the self-antigen GlialCAM^52^.

Resolving tissue-associated T cell differentiation requires integration of clonal identity, phenotype, antigen specificity and localization. Previous studies have largely addressed these dimensions in isolation, precluding intraclonal comparisons across tissues^16^. Our findings extend earlier observations that CCR5^+^ T_EM_ cells preferentially accumulate in the CNS of MS patients^53^. Although our data cannot determine the directionality of migration, our analyses linked a distinct differentiation program, characterized by *GZMK, CXCR3, GPR183* and *CD27* expression, to preferential CSF accumulation and EBV specificity. Independent support for the biological relevance of this program comes from the observation that CXCR3 is a strong regulator of CNS migration in a preclinical MS model^28^. More broadly, our analytical strategy provides a framework for dissecting tissue-associated clonal differentiation programs in human T cell responses beyond multiple sclerosis.

Our antigen-resolved atlas demonstrates that pathogen specificity is closely linked to tissue-associated differentiation. Previous studies have identified individual EBV-specific T cells in the CNS of patients with MS, but their limited scale precluded systematic comparisons across pathogens and differentiation states^5,22^. Our study substantially expands the current landscape of antigen-specific T cell responses in the human CNS, highlighting that pathogen specificity shapes distinct tissue-associated differentiation programs. Influenza-specific T cells showed similarly strong CSF enrichment as EBV-specific cells but occupied a more T_RM_-like differentiation state. In contrast, EBV-specific T cells possessed a CSF-biased effector-memory program already in peripheral blood while also constituting the dominant virus-specific population among expanded CSF clones.

The CSF-biased differentiation program is defined not only by location, but also by cellular interactions within the CNS. Independent experimental approaches in our study converged on interactions between CD8^+^ and B lineage cells. This suggests that non-cytotoxic CD8^+^ T cell functions may contribute more broadly to (CNS) inflammation than previously appreciated. This interpretation is supported by the presence of pro-inflammatory GZMK but absence of the cytotoxicity mediator GZMB^54,55^. Consistently, B cells in T–B multiplets lacked apoptotic signatures and instead displayed enhanced antigen presentation and cellular activation. Further evidence comes from a recent report that anti-CD20 treatment leads to an indirect reduction of GZMK^+^ CD8^+^ T_EM_ cells^56^. Of note, the spatial co-occurrence signal in brain lesions reflects shared tissue neighborhoods rather than demonstrated cell-to-cell contact, and analysis of more active lesion sections from more donors is required for replication.

Our *in vitro* functional assays recapitulated key cellular interactions predicted by the *ex vivo* analyses. Activated CD8^+^ T_EM_ cells promoted B cell differentiation toward an atypical CXCR3^+^ CD11c^+^ phenotype and plasmablast formation^12,23^. Further, GZMK-enriched CCR5^+^ CD8^+^ T_EM_ cells selectively induced IL-6 and IL-8 release from myeloid cells. These findings mirror recent observations in systemic lupus erythematosus (SLE), where GZMK^+^ CD8^+^ T cells promote atypical B cell differentiation^57^. Tissue-associated PD-1^+^ CD8^+^ T_EM_ cells can produce IL-21 and provide B cell help in chronic inflammatory settings^58–60^ and myeloid-derived IL-6 promotes differentiation of such IL-21-producing ‘helper CD8^+^ T cells’^61^. While the precise molecular mediators *in vivo* remain to be validated as part of complex multicellular networks, our data support a model in which the CSF-biased effector-memory program amplifies local B and myeloid cell responses, establishing an inflammatory feed-forward loop.

This model is compatible with evidence implicating EBV-infected B cells in CNS immune niches^12–14^, but does not require that interacting B cells remain EBV-infected or that every interacting T cell retains EBV specificity. The detection of EBV in B cells in the context of MS and SLE remains challenging despite technological progress^22,62–64^, but anti-CD20 treatment leads to dramatic reduction of peripheral EBV viral titers^65^. Our findings are consistent with the possibility that a major contribution of EBV occurs during peripheral immune priming, where the exceptionally large and prolonged antiviral response establishes a tissue-oriented differentiation program that subsequently participates in CNS inflammation^16,66^. Such a model places pathogen-driven immune imprinting within a broader context of host genetics and environmental factors. In this regard, it is intriguing that polymorphisms in *EOMES*, a central regulator of the differentiation program identified here^67^, are strongly linked to both MS susceptibility and EBV DNAemia, providing orthogonal human genetic support for this biological axis^68–70^. How similar tissue-associated differentiation programs operate across different autoimmune disorders, and even during ‘inflammaging’^71^, remain open questions.

Together, our findings identify a tissue-oriented, EBV-associated effector-memory program that links peripheral antiviral immunity to local inflammatory niches in the CNS.

## Methods

### Patient samples

Ethics approval (identifier 23-225-Bp) was granted by the local Ethics Committee of the Medical Faculty of the University Hospital of Erlangen, Friedrich-Alexander University Erlangen-Nürnberg, Germany. Samples were collected after informed consent of the donors. The cohort is described in detail in **Table S1**. The cohort included five MS patients (5/5 female; mean age 31.6 years), five NIC (2/5 female; mean age 36.6 years), and two inflammatory controls (IC; 1x ulcerative colitis and 1x non-MS neuropathy-like condition; 2/2 female; mean age 34.5 years). All MS patients were EBV-seropositive and showed high anti-EBNA IgG titers, consistent with previous observations^24^ (**Fig. S1b**). Three of five MS donors showed markedly higher cell counts in CSF compared to controls (**Fig. S1c**). All MS patients were untreated at the time of sampling. Patients were HLA genotyped using next generation sequencing at the University Hospital of Erlangen.

CSF samples were obtained via lumbar puncture during diagnostic procedures or during routine clinical care.

Peripheral blood mononuclear cells (PBMCs) were isolated from citrated peripheral blood by density-gradient centrifugation using BioColl density medium (BioSell; BS.L 6115). PBMCs were cryopreserved in heat-inactivated fetal calf serum (FCS) (anprotec; AC-SM-0027) + 10% dimethyl sulfoxide (DMSO) in liquid nitrogen.

### Cell cultures

All cell lines were cultured at 37 °C, 5% CO_2_ unless otherwise stated. THP-1 cells were a gift from Prof. Dr. Matthias Tenbusch (University Hospital of Erlangen). THP-1 cells were cultured in RPMI 1640 Medium (RPMI, Life Technologies; 21875091) with 1% L-glutamine (Fisher Scientific; 25030024) and 1% penicillin–streptomycin (Life Technologies; 15140122), and 10% heat-inactivated FCS (anprotec; AC-SM-0027). THP-1 cells were split regularly to a starting density of 10^5^ cells/mL. HEK293T cells were cultured in DMEM (Life Technologies, 11995073) supplemented by 10% heat-inactivated FCS, 0.05 mg/mL gentamicin (Life Technologies; 15750060), 0.05 mM β-mercaptoethanol (Life Technologies; 31350010), 1.1915 g/L Hydroxyethyl piperazine ethane sulfonic acid (HEPES; Carl Roth; HN77.3), 0.2 g/L L-glutamine, 100 U/mL penicillin, and 100 μg/mL streptomycin.

Primary human umbilical vein endothelial cell (HUVEC) cultures were obtained from Prof. Felix B. Engel (University Hospital of Erlangen). Freshly isolated HUVEC were cultured at 37 °C, 7.5% CO_2_ in Endothelial Cell Growth Medium (Promocell; C-22110).

Cryopreserved PBMC samples were thawed and rested overnight (37 °C; 5% CO_2_) in complete RPMI (cRPMI) containing RPMI, 10% heat-inactivated FCS (anprotec; AC-SM-0027), 0.05 mM β-mercaptoethanol (Life Technologies; 31350010), 1.1915 g/L HEPES (Carl Roth; HN77.3), 0.2 g/L L-glutamine (Fisher Scientific; 25030024), 100 U/mL penicillin, 100 μg/mL streptomycin (Life Technologies; 15140122), 0.05 mg/mL gentamicin (Life Technologies; 15750060) at a density of 1-2 × 10^6^ PBMCs/mL.

### Flow cytometry

The following antibodies were used for sorting live CD14^-^CD19^-^CD3^+^CD4^+^ T cells and live CD14^-^CD19^-^ CD3^+^CD8^+^ T cells for sc-seq: TotalSeq™-C0072 anti-human CD4 (300567; 1:200), TotalSeq™-C0046 CD8a (344753, 1:200), TotalSeq™-C0156 CD95/Fas (305651, 1:200), TotalSeq™-C0147 CD62L (304851, 1:588), TotalSeq™-C0063 CD45RA (304163, 1:588), anti-CD3-BV510 (317331, 1:100), anti-CD19-FITC (302205, 1:100), and anti-CD8α-APC (301049, 1:100) from BioLegend, along with CD14-APC/Cyanine7 (557831, 1:100) from BD Biosciences, and anti-CD4-PE (12-0019-42, 1:100) from Thermo Fisher. Sorting live CD14^-^CD19^-^CD3^+^CD8^+^pHLA^+^ cells for sc-seq used the following antibodies: TotalSeq™-C0072 anti-human CD4 (300567; 1:200), TotalSeq™-C0046 CD8a (344753, 1:200),

TotalSeq™-C0156 CD95/Fas (305651, 1:200), TotalSeq™-C0147 CD62L (304851, 1:588), TotalSeq™-C0063 CD45RA (304163, 1:588), TotalSeq™-C0148 anti-human CD197/CCR7 (353251, 1:250), TotalSeq™-C0140 anti-human CD183/CXCR3 (353747, 1:333), TotalSeq™-C0366 anti-human CD184/CXCR4 (306533, 1:200), TotalSeq™-C1484 anti-human GPR183 (EBI2) (368923, 1:200), TotalSeq™-C0154 anti-human CD27 (302853, 1:333), anti-CD3-BV510 (317331, 1:100), anti-CD19-FITC (302205, 1:100), anti-CD8α-BV421 (301035, 1:200), CD14-APC/Cyanine7 (557831, 1:100) from BioLegend and anti-CD4-FITC (555346, 1:50) from BD Biosciences. Wherever necessary, we included ‘hashtag’-antibodies from BioLegend to multiplex samples for sc-seq, e.g., TotalSeq™-C0251 anti-human Hashtag 1 (394661). Up to 8 hashtag-positive samples were used at a time.

The following antibodies were used for surface phenotyping of T cells: anti-CD8a-BV421 (301035; 1:200), anti-CD45RA-PerCP-Cy5.5 (304121, 1:200), anti-CD62L-APC/Cy7 (304813; 1:100), anti-CD27-BV711 (302833; 1:100), anti-CXCR3-PE-Dazzle™ 594 (353735; 1:100), anti-CCR5-BV605 (359145; 1:50), and anti-GZMK-APC (370509, 1:20) from BioLegend, as well as anti-GZMA-FITC (568389, 1:20), anti-GZMB-PE (561142, 1:20), anti-CD3-PE-Cy7 (737657,1:200), and anti-CD4-BV786 (740962; 1:400) from BD Biosciences. A ZombieAqua or Zombie NIR Fixable Viability Kit (BioLegend; 4231017/423102; 1:500) was used for viability staining. Additional antibodies were used and reported as part of specific in vitro experiments.

2-8 million overnight-rested PBMCs were stained in 96-well V-bottom plates at 4 °C and washed in FACS buffer (phosphate-buffered saline (PBS) + 0.5% bovine serum albumin (BSA)). For surface staining, cells were incubated with surface antibodies and viability dye for 30 min at 4 °C unless otherwise specified and washed 3 times prior to acquisition or cell sorting in FACS buffer.

For intracellular staining, cells were fixed and permeabilized using a BD Cytofix/Cytoperm Kit (BD Biosciences, 554714) for human samples, then incubated for 60 min with intracellular antibodies in PermWash.

Cells were acquired on a Cytek Northern Lights flow cytometer. Cells were sorted using a MoFlo Astrios EQ (1 and 2) at the Core Unit for Cell Sorting and Immunomonitoring in Erlangen.

### Sample processing for sc-seq

CSF was shortly after lumbar puncture measured (volume in mL) and spun down at 300 g, 10 min, at 4 °C. Supernatants were measured and carefully removed. Cells were transferred in approximately 0.5 mL leftover volume to sterile-filtered 2% BSA saturated Eppendorf tubes and resuspended in an equal volume of filtered cRPMI. CSF immune cells were then transported on ice within the hospital grounds (< 30 min) and spun down upon arrival to check for visible erythrocyte contamination and cell counting. Visible erythrocyte contamination of CSF immune cells served as a strict exclusion criterion of the sample and patient.

Cells were counted by first resuspending each CSF sample in 80 µL sterile-filtered 0.5% BSA (in PBS). 6 µL were mixed with 6 µL Trypan Blue (T8154, Thermo Fisher) and counted using a Neubauer chamber to ensure that no more than 25,000 cells were injected into the sc-seq cartridges from 10x Genomics.

### Single-cell sequencing

CSF and sorted immune cell populations were brought up to 1 mL of sterile-filtered 0.5% BSA (in PBS) and spun down at 600 g, 10 min at 4 °C before injection into the Next GEM microcontroller cartridge (10x Genomics). Further processing and sc-seq library construction included scRNA-seq, scTCR-seq, and CITE-seq and was done following manufacturer protocols (Chromium Next GEM Single Cell 5’ Reagent Kits v2 (Dual Index)). Sequencing was performed by Novogene Munich, Germany on an Illumina NovaSeq platform with PE150 strategy.

### Preprocessing of sc-seq data

All sequencing runs were separately processed by the Cell Ranger ‘multi’-command (version 9.0.1, 10x Genomics) with the gene expression reference GRCh38 (version 2020A, 10x Genomics), the VDJ reference vdj-GRCh38 (version 5.0.0, 10x Genomics), and customized feature barcode references. The remaining single-cell processing was conducted via Scanpy (v.1.11.4), Scirpy (v.0.22.3), Muon (v0.1.7), and Scrublet (v. 0.2.3)^72^. Low-quality cells with < 200 number of genes and ≤ 400 unique molecular identifier (UMI) counts were pruned, before UMI count normalization (CP10K), and transformation using the shifted logarithm. Low-quality cells with more than 10% mitochondrial genes were removed as well. Multiplets were annotated by running Scrublet for individual samples. Normalization of feature barcode-associated UMI counts was performed using dsb normalization implemented in Muon. Hashtag sample demultiplexing was performed using Hash Solo^73^ implemented in Scanpy. Hashtag-negative cells and heterotypic doublets consisting of cells with different hashtag labels were removed. Two CSF samples were multiplexed without hashtag antibodies. Sample demultiplexing was performed using scSplit^74^ and heterotypic doublets were discarded.

### Clonotype annotation

TCR clonotype annotation, i.e. annotation of TRA and TRB chains of the TCR was performed using Scirpy v0.22.3. Clonotypes were defined as sharing 1 TRA and 1 TRB amino acid sequence. Both primary and secondary chains were considered. Singlets and multiplets were annotated separately.

### Batch correction and feature selection

Individually preprocessed samples were concatenated and assessed for batch effects defined as sample-specific clusters unique to a Next GEM cartridge lane. No clear batch effects were evident in the initial clustering and UMAP projections of CD4^+^ and CD8^+^ T cells from PB and CSF. In contrast, pHLA multimer-binding CD8 T cells sorted from all donors and sequenced across two different Next GEM cartridge lanes exhibited clear batch effects. For pHLA multimer-binding, CD8^+^ T cells batches were corrected using scVI^75^. A soft batch correction was performed for other analyses by selecting highly variable genes (HVGs) seen in multiple batches. This mimics ‘SelectIntegrationFeatures’ in Seurat^76^. A batch was here defined by the specific Next GEM cartridge lane and by sample donor ID.

### Sc-seq analysis

Leiden clusters and UMAP projections were calculated using Scanpy. TRA and TRB-related genes were removed from the list of HVGs. All cells were initially clustered at high Leiden cluster resolution (res. 4, 5000 HVGs) and annotated as either CD4, CD8, CD4/CD8 or MAIT cells according to the expression of *CD4, CD8A, CD8B*, CD4 protein, and CD8 protein. Protein expression was prioritized, and voting was performed per observed clonotype. T cells labeled and sorted from PB were fully identifiable by protein expression alone. All CSF T cells were subclustered (res. 2, 5000 HVGs) and remaining negative cells in the CSF were annotated based on gene expression including CD4 or CD8 annotation of each CSF T cell cluster followed by intra-clonotype label voting to annotate co-receptor negative cells. MAIT cells and CD4/CD8 T cells were excluded from further analysis.

Whenever subsequent analyses concerned singlets only, we removed multiplets before continuing by discarding cells with a high number of genes detected (≥ 3000 genes), or a high total UMI count (≥ 4000), or a high Scrublet score (≥ 0.5). An initial round of Leiden clustering at high resolution (res.4.0, 5000 HVGs) and UMAP projection was used to identify and exclude small outlier clusters with high % of doublets discerned by the number of genes, number of UMI counts, and by the Scrublet score, as well as by a high frequency of two-chain TCRs.

CD4^+^ and CD8^+^ singlet T cells were analyzed separately after gene-count renormalization. Leiden clusters were then generated at medium resolution (res. 1.8, 6500 HVGs). Clusters were manually annotated according to the expression of known subset-specific markers including CITE-seq markers CD4, CD8A, CD45RA, CD62L, and CD95^77^. Differential gene expression analyses by Scanpy (scanpy.tl.rank_genes_groups) were performed using a Wilcoxon test adjusted for multiple comparisons and a Benjamini-Hochberg procedure considering genes expressed in more than 15% of either immune cell population.

### Annotation of CSF- and PB-biased clonotypes

Statistical thresholds to annotate CSF-biased and PB-biased clonotypes were individually computed for CD4^+^ and CD8^+^ T cell clonotypes. To compute thresholds, we used permutation of a binomial distribution using the relative proportions of total CSF and PB T cells as estimates of the expected frequencies under random sampling of the combined PB and CSF datasets. We computed such thresholds for different expansion thresholds, and selected expansion thresholds where both CSF-biased and PB-biased clonotypes could be identified at *P* < 0.05.

### DNA barcode-labeled pHLA multimer libraries

DNA-barcoded pHLA multimer libraries were designed as described by Povlsen et al.^78^. In brief, biotinylated, single-stranded DNA-barcodes compatible with the 5’ capture chemistry from 10x Genomics were purchased from LCG Biosearch Technologies (Denmark) or Integrated DNA Technologies (USA) and dissolved to 100 μM in barcode dilution buffer (PBS with 1 mg/mL herring-DNA, 2 mM EDTA and 0.5% (wt/vol) BSA) before long-term storage at −20 °C. For a working solution, the DNA barcodes were further diluted to 0.54 μM in barcode dilution buffer. DNA barcodes were conjugated to 160 nM APC- or PE-labeled, streptavidin-dextran backbones from Fina Biosolutions LLC (USA) in a 1:1 molar ratio and incubated between 0.5-24 h at 4 °C.

Recombinant pHLA folded with a UV-cleavable peptide was produced following published work^79,80^. In brief, recombinant β2 microglobulin (B2M) and heavy chains of HLA-A*01:01, A*68:01, B*07:02, B*08:01, B*15:01, B*35:01, and C*07:02 were expressed in *Escherichia coli* using pET series expression plasmids. Denatured heavy chains and B2M were subsequently isolated as inclusion bodies and refolded in the presence of a UV-cleavable peptide. Refolded, recombinant HLA proteins were biotinylated (BirA biotin-protein ligase standard reaction kit, Avidity LLC, Aurora, CO) and purified using HPLC (Waters Corporation, USA). HLA-A*02:01 and HLA-A*24:02 were folded and purified as empty disulfide-stabilized complexes as described previously^81^. All HLA-I folded monomers were quality-checked for their concentration, UV degradation, and biotinylation efficiency and stored at −80 °C.

All viral epitope sequences were published previously^82–85^. 2 µg lyophilized, crude synthesis product for each peptide was purchased from Biosynth (USA) or from Synpeptide (China), and resuspended in pure dimethyl sulfoxide (DMSO) to a stock concentration of 10 mM and stored at −20 °C.

When loading peptides into HLA-I monomers, peptides were first diluted to 200-400 μM in fresh PBS. Recombinant loadable-pHLA molecules were diluted to 100-150 µg/mL (~2-4 μM). Diluted peptides and empty loadable-pHLAs were mixed 1:1 (v/v) and exposed to UV-light (366 nm) for 1 h to facilitate peptide exchange. Empty, loadable monomers, HLA-A*02:01 and A*24:02, were loaded for 30 min at room temperature (RT). Peptide-loaded pHLA monomers (50-100 µg/mL, i.e., ~1-1.5 μM) were spun at 3300 g, 4 °C, 5 min to pellet aggregates. Refolded pHLA monomers were conjugated to DNA-barcode labeled backbones in a 16-24 molar ratio and incubated for 30 min at 4 °C before addition of barcode freezing buffer (PBS with 9 µM D-biotin, 625 µg/mL herring DNA, 12 mM EDTA, 3% (wt/vol) BSA and 30% (vol/vol) glycerol) to a final pHLA multimer concentration of 35.2 nM. The resulting multimers, containing 1.4 µM D-biotin, 100 µg/mL herring DNA, 1.9 mM EDTA, 0.5% BSA, and 5% glycerol, were stored at −80 °C.

Each batch of functionalized dextrans and pHLA monomers was tested for its capacity to stain a known antigen-specific population of memory T cells from previously tested anonymous blood donors by titrating the pHLA concentration. HLA-wise pools of pHLA multimers were furthermore tested for their capacity to stain a known antigen-specific population without background before staining patient samples.

EBV-associated peptides were assembled for pHLA multimers on a PE-SA-dextran backbone. Other virus-derived peptides were assembled on an APC-SA-dextran backbone.

### Staining cells with pHLA multimer libraries

Frozen pHLA multimers were thawed on ice and spun for 5 min at 3300 g, 4 °C to pellet aggregates. HLA-specific pHLA multimer pools were made in BSA-saturated reservoirs from 3 μL of each DNA barcode-labeled pHLA multimer. HLA-wise multimer pools were concentrated, if necessary, using BSA-saturated Vivospin 6 columns (100,000 Da PES filter, VS0642, Sartorius) to 25 μL per stain. HLA-wise multimer pools were transferred to BSA-saturated, LoBind Eppendorf tubes, volume-adjusted slightly using sterile-filtered 0.5% BSA, and stored on ice before staining overnight-rested PBMCs. HLA-wise multimer pools were centrifuged at 10,000 g, 2 min, 4 °C, which was repeated in fresh LoBind Eppendorf tubes twice. 4-6 million PBMCs were washed twice and resuspended in 25-100 μL pHLA multimers matching their HLA genotypes depending on the donor. Staining volumes were subsequently adjusted to 100 μL using sterile-filtered FACS buffer. HLA-A*02:01 was stained separately from other HLA alleles due to the size of the HLA-A*02:01 library. Multimers were incubated with PBMCs for 60 min at 4 °C before washing in sterile-filtered FACS buffer. PBMCs were subsequently stained with surface antibodies including fluorescently labeled antibodies, hashtag antibodies, and CITE-seq antibodies. Surface antibodies were spun for 14,000 g, 10 min, 4 °C to remove aggregates before staining. PBMCs were stained at 4 °C for 30 min before washing in sterile-filtered FACS buffer and sorting cells into 100 µL sterile-filtered 0.5% BSA. Cells were transported and stored on ice before cell sorting and sc-seq following manufacturer protocols (Chromium Next GEM Single Cell 5’ Reagent Kits v2 (Dual Index)) with adjusted read depths (GEX: 20,000, TCR: 8,000, CS: 10,000). Sequencing was performed by Novogene Munich, Germany on an Illumina NovaSeq platform with PE150 strategy. Cell Ranger was used for preprocessing multimer-attached DNA barcodes (the same as for TotalSeq-C, CITE-seq antibodies). Subsequent preprocessing was performed using dsb normalization of feature barcode counts. DextraDemixer was used to annotate pMHC multimer-binding clonotypes^86^. To further increase the specificity of DextraDemixer only multimer-annotated clones with 2 or more cells were included in downstream analysis.

### B-LCL transformation and longitudinal monitoring using TCR-seq

1-2 million PBMCs from each donor were transformed using filtered supernatants from the B95-8 parental cell line. Trizol (Thermo Fisher; 15596018) lysates from 500,000 cells were cryopreserved during periods of culturing on day 0 and day 19. RNA was extracted using the manufacturer protocol, and used as input for cDNA synthesis for bulk TCR-seq following published protocols^87^ with the following modifications. First strand synthesis was performed as a double reaction (2 × 20 µL) using up to 600 ng RNA, as well as materials from the SMARTScribe Reverse Transcriptase kit (Cat. no: 639538, Takara) together with a custom template switch oligo and reverse primers matching the constant region of the TRB chain (See table S2 for a list of primers). cDNA was generated by incubation at 42 °C in a thermocycler for 60 min. 2 µL of Uracyl DNA glycosylase (New England Biolabs; M0280L) were added to each reaction and incubated at 37 °C for 40 min. cDNA purification was done using AMPure XP beads (1x ratio) (Beckman Coulter, A63881). Eluted cDNA (27 µL) was used for PCR amplification (PCR 1), which ran as published elsewhere using a Q5 polymerase (NEB; M0491L)^87^. cDNA purification was done using AMPure XP beads (0.8x ratio). Qubit for detection of dsDNA (Thermo Fisher; Q32851) was used to quantify the library. We used 13.5 ng of cDNA as input for further PCR amplification (PCR 2), which added incomplete adapter-sequences, custom barcodes as sample IDs, and heterogeneity spacers via 5’-overhangs on the given primers. PCR amplification was performed as follows: 98 °C for 30 s, 10 cycles of denaturation at 98 °C for 10 s, annealing at 58 °C for 10 s and extension at 72 °C for 50 s. A final extension was performed at 72 °C for 2 min. A double-sided AMPure XP purification (0.5x → 0.8x) was used to purify cDNA after PCR 2. Indexing PCR used Nextera™-compatible indexing primers with unique dual indexes (IDT). A PCR master mix was prepared containing 1× Q5 reaction buffer (10 µL of 5× buffer per reaction), 0.2 mM each dNTP (1 µL of 10 mM dNTP mix), 0.4 µM C01 primer mix (2 µL of 10 µM primer mix) and 0.02 U/µL Q5 High-Fidelity DNA Polymerase (0.5 µL of 2 U/µL polymerase; NEB). PCR amplification at 98 °C for 3 min, 8 cycles of denaturation at 98 °C for 30 s, annealing at 60 °C for 30 s and extension at 72 °C for 30 s. A final extension was performed at 72 °C for 5 min, followed by a hold at 4 °C. Libraries were purified within 1 h of index PCR using a 1.8x single-sided cleanup with AMPure XP magnetic beads. Library quality and concentration were assessed using a TapeStation (Agilent), and sequencing was performed using PE150 configurations by Novogene at read depths of 60-95 reads per cell. Clonotypes based on TRB chains were called using MiXcR^88^, and relative frequencies were studied to compare differences in clonal expansion of T cell clones at day 19 compared to day 0. Any clonotype with positive fold changes at day 19 was defined as LCL-reactive in the sc-seq analyses given exact matching between TRB sequences.

### Imaging cytometry

PB and CSF samples for imaging cytometry were collected from a separate cohort of MS patients and NIC from Magdeburg, Germany.

Patients’ peripheral blood was obtained according to the Declaration of Helsinki upon consent (ethical approval number 61/22 at the Ethical Committee of the Medical Faculty at Otto-von-Guericke University Magdeburg, Germany) and processed within 1 h for peripheral blood mononuclear cell (PBMC) isolation using gradient density centrifugation with Pancoll (PAN Biotech, 1.077 g/mL density) in Leucosep tubes (Greiner bio-one) according to manufacturer’s recommendations. PBMCs were cryo-conserved in medium containing 50% FCS and 10% DMSO until further use.

Patients’ CSF was obtained according to the Declaration of Helsinki upon consent (ethical approval number 07/17 at the Ethical Committee of the Medical Faculty at Otto-von-Guericke University Magdeburg, Germany) and processed within 30 min. Briefly, CSF volume was topped to 15 mL with PBS + 0.04% BSA and centrifuged for 10 min at 300 x g, 4 °C without break. Cells were fixed immediately with 1.5% PFA and directly used for further analysis.

### Staining and analysis

Samples were washed with PBA (PBS + 0.5% BSA +0.01% NaN3) by centrifugation, blocked for unspecific bindings with 3 mg/mL Gamunex (human IgG, Grifols), and stained with specific fluorochrome-conjugated surface antibodies. The following antibodies were used for surface staining: anti-CD3-BV786 (clone OTK3; BD Biosciences; 566781; 1:20), anti-CD14-PE (clone HCD14; BioLegend; 325606; 1:33.3), and anti-CD19-PE/Cy7 (clone HIB19; BioLegend; 302216; 1:20). After washing with PBS + 2% FCS + 0.1% Triton-X, samples were counterstained with 1 µM DAPI and subjected to imaging cytometry on a FlowSight system (Cytek). Recorded data was analyzed using IDEAS 6.2 software (Cytek).

### Re-analysis of spatial transcriptomics data

24 Visium sections were analyzed: 18 sections from Lerma-Martin et al.^42^ (GSE279181; 6 control white matter, 8 chronic active and 4 chronic inactive lesions) and 6 sections of one donor from Schroeder et al.^43^ (Zenodo 15151517; chronic active white matter). In total, 14 donors were re-analyzed (6 active, 2 inactive, 6 control). Raw counts per spot were taken from the DecontX-corrected object (4,613 genes shared by both deposits).

Per-spot cell type abundances were estimated with cell2location in two stages: reference expression signatures per cell type were learned from a harmonized single-nucleus reference of 171,704 nuclei from 24 donors, pooling two published cohorts^42,89^ (donor as batch, study cohort as covariate, 4,000 epochs), and the spatial model was then fitted per section (detection alpha 20, up to 5,000 epochs, 1,000 posterior samples); the 5th posterior percentile of the abundance was used as a conservative per-spot estimate. The run used here was fitted with a reduced reference of 15,482 nuclei across 29 cell types. Only the myeloid abundance tracked its own marker transcripts (*AIF1, C1QB, CSF1R, P2RY12, CD14*) closely enough to be used (depth-stratified area under the receiver operating characteristic (AUROC) of marker-positive against marker-free spots at least 0.65 in 14 of 18 lesion sections). CD8 T cell and B lineage signals were therefore read from the transcripts themselves rather than from the deconvolution.

### A spot counted as positive when it carries

CD8 T, at least one *CD8A, CD8B* transcript; B lineage, at least one *CD79A, CD79B, IGHM* transcript (B cell or IgM-secreting plasma cell); T cell, at least one *CD3E, CD3G, CD2, TRAC* transcript; CD4 (proxy), a T cell spot without a *CD8* transcript; NK, at least one *NKG7, GNLY, KLRD1* transcript and no T cell transcript; myeloid, cell2location myeloid abundance (5th posterior percentile) of at least 0.14 cells per spot. The myeloid threshold was the 95th percentile of the cell2location abundance in control spots without any transcript of *AIF1, C1QB, CSF1R, P2RY12, CD14*. The CD4 arm used T cell spots without a CD8 transcript as a proxy, because the CD4 transcript in this tissue tracks myeloid cells. All rules were fixed before any co-occurrence result was computed.

### Neighborhood co-occurrence

The neighborhood of a spot is the spot itself and its six adjacent spots (100 µm center to center). For each section, the risk difference was calculated as the fraction of target spots containing a neighborhood of interest minus the same fraction among the other lymphoid spots (T cell, NK cells or CD4 T cells) of that section. Its null distribution is a Monte Carlo permutation of the neighborhood of interest in absence of observed differences between CD8^+^ spots and other lymphocyte subsets. Sections were combined per donor by inverse-variance weighting and across donors by DerSimonian-Laird random effects, with the Hartung-Knapp small-sample *P* value reported alongside the normal-approximation *P*. The primary hypothesis, pre-specified in the positive direction, is that in active lesions CD8 T spots have a B lineage spot in their neighborhood more often than the other lymphoid spots (two-sided *P* values); the same test additionally requiring a myeloid spot is secondary and Benjamini-Hochberg adjusted. Inactive lesions, control white matter and the CD4-proxy comparison are descriptive and not formally tested. A section was not estimable with fewer than 20 target or background spots, when the background fraction exceeds 0.95 (one active section) or when the null has no variance.

### Granzyme expression

Granzyme expression was read from the full Visium objects, normalized to counts per 10,000 and log1p-transformed per spot. The 4,613-gene deconvolution panel lacked GZMA, GZMB and GZMM. Lesion effects were estimated by ordinary least squares of spot-level normalized expression on two indicator variables, active lesion and inactive lesion, with control white matter as the reference level; standard errors were clustered on section, so precision reflects the 24 sections rather than the 85,745 spots, and 95% confidence intervals are the coefficient ± 1.96 clustered standard errors. The active-versus-control coefficient is informed by 20 sections from 12 donors (14 active, 6 control); *P* values across the eight tested genes were corrected by the Benjamini-Hochberg procedure. Analyses were run in Python 3.10.20 with statsmodels 0.14.6.

### Co-culture of sorted CD8^+^ T cell subsets with B cells

Cryopreserved PBMCs from leukoreduction system chambers (LRSC) of healthy donors were rested in cRPMI (RPMI 1640 Medium (Life Technologies; 21875091), 10% heat-inactivated FCS (anprotec; AC-SM-0027), 0.05 mM β-mercaptoethanol (Life Technologies; 31350010), 1.1915 g/L HEPES (Carl Roth; HN77.3), 0.2 g/L L-glutamine (Fisher Scientific; 25030024), 0.05 mg/mL gentamicin (Life Technologies; 15750060) and 100 U/mL penicillin–streptomycin (Life Technologies; 15140122)). Cells were washed with FACS buffer, counted and stained for 20 min at RT with anti-CD4-PE/Cy7 (BD Biosciences; 560649; 1:200), anti-CD62L-APC/Cy7 (BioLegend; 304813; 1:100), anti-CD45RA-PerCP/Cy5.5 (BioLegend; 304121; 1:400), anti-CCR5-APC (BioLegend, 359122, 1:50), anti-CD8-FITC (Beckman Coulter; A07756; 1:200), anti-CD19-BV421 (BD Biosciences; 562440, 1:400) and anti-IgD-PE (BioLegend; 348204; 1:400). Cells were pregated on CD4^-^ single lymphocytes and sorted into CD19^+^IgD^−^ B cells and CD8^+^ T cell subsets: CCR5^+^ T_EM_ (CD45RA^−^CD62L^−^), CCR5^-^ T_EM_ and T_E_ (CD45RA^+^CD62L^−^) cells.

Sorted cells were resuspended in cRPMI with 50 U/mL IL-2 (Life Technologies; 200-02-250UG), and 1 × 10^5^B cells and 1 × 10^5^T cells of the indicated subsets were combined per well in a 96-well round-bottomed plate. For stimulation, wells were precoated overnight (4 °C) with anti-CD3 (1 µg/mL; BioLegend, 317302) and anti-CD28 (1 µg/mL; BioLegend; 302902). Plates were briefly centrifuged (480 g, 20 s) and incubated for 72 h (37 °C, 5% CO_2_).

At 0 h and 72 h, counting beads (123count eBeads; Life Technologies; 01-1234-42) were added for absolute quantification of cells. Cells were stained for 20 min at RT with anti-CD45RA-PerCP/Cy5.5, anti-CCR5-APC, anti-CD8-FITC, anti-CD19-BV421, anti-IgD-PE, anti-CD27-BV510 (BioLegend; 302835; 1:100), anti-CD38-BV605 (BD Biosciences, 562666; 1:400), anti-CD11c-PE/Cy7 (BD Biosciences; 561356; 1:400), anti-CD95-BUV737 (BD Biosciences; 612790; 1:20), anti-CXCR3-PE/Dazzle 594 (BioLegend; 353735; 1:100) and Zombie NIR viability kit (BioLegend; 423102; 1:1000), then acquired on a Cytek NorthernLights instrument.

### Terminal complement complex formation on HUVEC, HEK293T and THP-1 cells

Complement activation by recombinant GZMK was assessed by detection of terminal complement complex (TCC) formation using a C9 neoantigen antibody (Hycult Biotech; HM2264). HUVEC, HEK293T and THP-1 cells were harvested, centrifuged (300 g, 5 min) and resuspended in the respective serum-free culture medium (Endothelial Cell Growth Medium (Promocell; C-22110) for HUVECs, DMEM for HEK293T, RPMI 1640 for THP-1). Cells were plated at 1 × 10^5^cells per well in 100 µL in a 96-well V-bottomed plate, centrifuged (300 g, 5 min), washed twice in serum-free RPMI (480 g, 5 min), and resuspended in serum-free medium.

Recombinant GZMK was prediluted in protein freezing buffer to a 20× working stock of 4000 nM immediately before use. Cells were incubated in a total volume of 100 µL with 2% C1q-depleted serum (Complement Technology; A300 Lot #26a) alone or supplemented with 50 nM or 100 nM GZMK, or with 20% active or heat-inactivated normal human serum (NHS; Complement Technology; NHS Lot #53b) as positive and negative controls, for 120 min at 37 °C. Serum was thawed immediately before use and kept on ice throughout.

After incubation, cells were centrifuged (300 g, 5 min), resuspended in 200 µL PBS and split into two plates for parallel GZMK and TCC staining.

For GZMK staining, cells were washed once in opsonization buffer and stained with anti-GZMK-APC (1:25) and ZombieAqua viability dye (1:1,000) for 45 min (RT, dark), followed by washing in opsonization buffer.

For TCC staining, cells were washed in PBS, stained with viability dye in PBS for 15 min (ice), and blocked with 10% goat serum (Thermo Fisher; 10000C) in PBS (100 µL per well) for 20 min (ice). Cells were washed in 10% HI-FACS buffer (10% FCS in PBS) and incubated with anti-C9 neoantigen antibody (Hycult Biotech; 1:50) in 25 µL per well for 30 min (4 °C, shaking), followed by PE-conjugated anti-mouse Fc secondary antibody (1:100) in 25 µL per well for 30 min (4 °C). Cells were washed, resuspended in 100 µL FACS buffer and acquired on a Cytek NorthernLights instrument.

### Co-culture of sorted CD8^+^ T cell subsets with THP-1 cells

The human monocytic leukemia cell line THP-1 (provided by Matthias Tenbusch, University Hospital Erlangen) was maintained in RPMI 1640 with 10% FCS, 1% L-glutamine and 1% penicillin–streptomycin (37 °C, 5% CO_2_). Cryopreserved PBMCs from LRSC of healthy donors were rested overnight in cRPMI (RPMI 1640 Medium, 10% heat-inactivated FCS, 0.05 mM β-mercaptoethanol, 1.1915 g/L HEPES, 0.2 g/L L-glutamine, 0.05 mg/mL gentamicin and 100 U/mL penicillin– streptomycin), then stained for 20 min at RT with anti-CD4-PE/Cy7, anti-CD62L-FITC, anti-CD45RA-PerCP/Cy5.5, anti-CD3-BV510 (BioLegend; 317331; 1:100), anti-CD8a-BUV496 (BD Biosciences; 612942; 1:200), anti-CD19-BV421 and anti-CCR5-APC. Cells were pregated on single, CD19^-^CD4^-^CD3^+^ lymphocytes and sorted into five CD8^+^ T cell subsets: T^N^ (CD45RA^+^CD62L^+^), T_CM_ (CD45RA^−^CD62L^+^), CCR5^+^ T_EM_ (CD45RA^−^CD62L^−^), CCR5^-^ T_EM_ and T_E_ (CD45RA^+^CD62L^−^) cells.

THP-1 cells were labeled with CellTrace Violet (CTV; 1 µM, 5 × 10^6^ cells per 100 µL) for 20 min at 37 °C, quenched with medium containing 20% FCS. Sorted T cells and CTV-labeled THP-1 cells were resuspended in RPMI 1640 with 10% human serum (Sigma Aldrich; H4522), 1% L-glutamine and 1% penicillin-streptomycin, and 2 × 10^4^ THP-1 cells and 2 × 10^5^T cells were combined per well (100 µL total volume) in a 96-well V-bottomed plate. Control wells received THP-1 cells alone, with 1 µg/mL LPS, or with recombinant GZMK (50, 100 or 200 nM; Enzo Life Sciences; ALX-201-117A-C010). Plates were briefly centrifuged (480 g, 20 s) and incubated for 4 h (37 °C, 5% CO_2_), alongside 0 h controls.

At 4 h, supernatants were collected and stored at −20 °C, and counting beads were added for absolute quantification. Cells were stained with Zombie NIR viability kit for 15 min. For surface GZMK detection, cells were stained with anti-GZMK-PE (BioLegend; 370512; 1:40) for 45 min at RT and fixed in 2% paraformaldehyde (Science Service; E15710). For intracellular staining, cells were fixed and permeabilized using a BD Cytofix/Cytoperm Kit (BD Biosciences; 554714), then stained with anti-GZMK-PE for 45 min at RT in PermWash. Samples were acquired on a Cytek NorthernLights instrument, and cell counts were normalized to counting beads.

### IL-6 and IL-8 ELISA

Cytokine concentrations in 48 h co-culture supernatants were quantified by sandwich ELISA (R&D systems, DY206 and DY208). High-binding 96-well plates were coated with capture antibody diluted in PBS (50 µL per well) and incubated overnight at 4 °C. Plates were washed three times with wash buffer (PBS, 0.05% Tween-20) and blocked with PBS containing 1% BSA for at least 2 h at RT. After three washes, standards were applied as a 1:2 serial dilution series in PBS/1% BSA starting from the highest concentration, with PBS/1% BSA alone as blank. 50 µL of supernatants were added per well at the appropriate dilution (1:50 for detection of IL-8; 1:5 for the detection of IL-6) in PBS/1% BSA and incubated overnight at 4 °C.

Plates were washed three times and incubated with 50 µL detection antibody (1:363.6) in PBS/1% BSA for 2 h at RT. After three washes, 50 µL of streptavidin-HRP diluted 1:40 in PBS/1% BSA was added and incubated for 20 min at RT, followed by three further washes. Substrate reagents A and B were mixed 1:1, and 50 µL of substrate mix was added per well. The reaction was developed until blue color appeared and stopped with 25 µL of 2 N H_2_SO_4_ per well. Absorbance was measured at 450 nm with reference wavelength correction at 540-570 nm.

### Statistics

Information regarding statistics pertaining to the analysis of sc-seq and spatial transcriptomic data is described separately under their respective sub headers. Ordinary two-sample hypothesis testing was performed using two-sided, non-parametric Mann–Whitney *U* test. For multiple comparisons, we used Dunn’s post hoc test and adjusted for multiple hypothesis testing using Benjamini-Hochberg or Šidák’s correction. Correlation analyses were performed using a non-parametric Spearman correlation. Statistical testing was performed in Python using the scipy v.1.16.2, or GraphPad Prism 10. Significance is defined as follows: * *P* < 0.05, ** *P* < 0.01, *** *P* < 0.001, **** *P* < 0.0001.

### Data analysis and visualization

Data graphs were generated using GraphPad Prism 10 or Python using Matplotlib v3.10.6 and Seaborn v0.13.2 or by using plotting functions integrated in individual modules. Schemes and figures were generated with Affinity Designer (Serif (Europe) Ltd, version 2.6.5).

## Data and code availability

All sc-seq data will be made publicly available after peer review. The source code used to analyze the sc-seq data is available at https://github.com/user-pagh/UKER.MS.EBV.scseq.analysis.

## Author contributions

Conceptualization: N.P.K., K.S., data curation: N.P.K., K.S., formal analysis: N.P.K., J.K.B., S.F., M.B., funding acquisition: N.P.K., K.S., investigation: N.P.K., T.T., J.K.B., S.F., L.K., P.H., M.G., M.V.S., V.L., M.B., C.S., methodology: N.P.K., J.K.B., H.R., B.S., D.M., D.H.H., S.M., P.G.T., project administration: N.P.K., K.S., resources: L.F.V., S.Scha., S.Schr., F.B.E., S.R.H., R.L., V.R., K.S., software: N.P.K., J.K.B., supervision: K.S., V.R., validation: N.P.K., visualization: N.P.K., K.S., writing original draft: N.P.K., K.S., review & editing: all authors.

## Acknowledgments

This work was mainly supported by the Lundbeck Foundation (project R402-2022-1390) to N.P.K. It was also made possible by grants from the German Federal Ministry of Research, Technology and Space (BMFTR, projects 01KI2013, 01KI2516 and 031L0290B), the Else Kröner-Fresenius-Stiftung (project 2020_EKEA.127) and the German Research Foundation (DFG) through the research training group RTG 2504 (project 401821119), the Heisenberg program (project SCHO 1949/1-1), the TRR221/3 (324392634, project A09) and the SFB 1755 CASCAID (550296805; project #13), the Aventis Foundation (project 2024: Life Science Bridge Award), as well as the Yellow4FLAVI project, funded by the European Union, under the Horizon Europe Programme, Grant Agreement N° 101137459 to K.S. Further support came from Scleroseforeningen (A44005) to L.F.V.

The study was funded by the European Union. Views and opinions expressed are, however, those of the authors only and do not necessarily reflect those of the European Union or HADEA. Funding agencies had no influence on the study design or implementation.

We thank all study participants for making this work possible. We thank members of the Schober laboratory for experimental support and critical discussion. Furthermore, we gratefully acknowledge the generous support of the Manfred Roth-Stiftung, Fürth, Germany, and thank the Core Unit Cell Sorting and Immunomonitoring Erlangen.

## Declaration on usage of generative AI

During the preparation of this work, the authors used ChatGPT (OpenAI) for language editing and manuscript refinement including consistency checks. After using this tool, the authors reviewed and edited all text and take full responsibility for the content of the article.

## Conflicts of interest

The authors declare no conflicts of interest.

## Supplementary Materials

Figures S1-S16

Tables S1-S2

## Notes

### Competing Interest Statement

The authors have declared no competing interest.

## References

1. Smyth, L.C.D., and Kipnis, J. (2025). Redefining CNS immune privilege. Nat. Rev. Immunol. 25, 766–775. 10.1038/s41577-025-01175-0.

2. Bjornevik, K., Cortese, M., Healy, B.C., Kuhle, J., Mina, M.J., Leng, Y., Elledge, S.J., Niebuhr, D.W., Scher, A.I., Munger, K.L., et al. (2022). Longitudinal analysis reveals high prevalence of Epstein-Barr virus associated with multiple sclerosis. Science 375, 296–301. 10.1126/science.abj8222.

3. Vietzen, H., Berger, S.M., Kühner, L.M., Furlano, P.L., Bsteh, G., Berger, T., Rommer, P., and Puchhammer-Stöckl, E. (2023). Ineffective control of Epstein-Barr-virus-induced autoimmunity increases the risk for multiple sclerosis. Cell 186, 5705-5718.e13. 10.1016/j.cell.2023.11.015.

4. Moutsianas, L., Jostins, L., Beecham, A.H., Dilthey, A.T., Xifara, D.K., Ban, M., Shah, T.S., Patsopoulos, N.A., Alfredsson, L., Anderson, C.A., et al. (2015). Class II HLA interactions modulate genetic risk for multiple sclerosis. Nat. Genet. 47, 1107–1113. 10.1038/ng.3395.

5. Angelini, D.F., Serafini, B., Piras, E., Severa, M., Coccia, E.M., Rosicarelli, B., Ruggieri, S., Gasperini, C., Buttari, F., Centonze, D., et al. (2013). Increased CD8+ T cell response to Epstein-Barr virus lytic antigens in the active phase of multiple sclerosis. PLoS Pathog. 9, e1003220. 10.1371/journal.ppat.1003220.

6. Gottlieb, A., Pham, H.P.T., Saltarrelli, J.G., and Lindsey, J.W. (2024). Expanded T lymphocytes in the cerebrospinal fluid of multiple sclerosis patients are specific for Epstein-Barr-virus-infected B cells. Proc. Natl. Acad. Sci. U. S. A. 121, e2315857121. 10.1073/pnas.2315857121.

7. Kaufmann, M., Evans, H., Schaupp, A.L., Engler, J.B., Kaur, G., Willing, A., Kursawe, N., Schubert, C., Attfield, K.E., Fugger, L., et al. (2021). Identifying CNS-colonizing T cells as potential therapeutic targets to prevent progression of multiple sclerosis. Med 2, 296-312.e8. 10.1016/j.medj.2021.01.006.

8. Smolders, J., Remmerswaal, E.B.M., Schuurman, K.G., Melief, J., Van Eden, C.G., Van Lier, R.A.W., Huitinga, I., and Hamann, J. (2013). Characteristics of differentiated CD8+ and CD4+ T cells present in the human brain. Acta Neuropathol. 126, 525–535. 10.1007/s00401-013-1155-0.

9. Skulina, C., Schmidt, S., Dornmair, K., Babbe, H., Roers, A., Rajewsky, K., Wekerle, H., Hohlfeld, R., and Goebels, N. (2004). Multiple sclerosis: Brain-infiltrating CD8+ T cells persist as clonal expansions in the cerebrospinal fluid and blood. Proc. Natl. Acad. Sci. U. S. A. 101, 2428–2433. 10.1073/pnas.0308689100.

10. Kavaka, V., Mutschler, L., de la Rosa Del Val, C., Eglseer, K., Gómez Martínez, A.M., Flierl-Hecht, A., Ertl-Wagner, B., Keeser, D., Mortazavi, M., Seelos, K., et al. (2024). Twin study identifies early immunological and metabolic dysregulation of CD8+ T cells in multiple sclerosis. Sci. Immunol. 9, eadj8094. 10.1126/sciimmunol.adj8094.

11. Montalban, X., Hauser, S.L., Kappos, L., Arnold, D.L., Bar-Or, A., Comi, G., de Seze, J., Giovannoni, G., Hartung, H.-P., Hemmer, B., et al. (2017). Ocrelizumab versus Placebo in Primary Progressive Multiple Sclerosis. N. Engl. J. Med. 376, 209–220. 10.1056/NEJMoa1606468.

12. Läderach, F., Piteros, I., Fennell, É., Bremer, E., Last, M., Schmid, S., Rieble, L., Campbell, C., Ludwig-Portugall, I., Bornemann, L., et al. (2025). EBV induces CNS homing of B cells attracting inflammatory T cells. Nature 646, 171– 179. 10.1038/s41586-025-09378-0.

13. Kim, H., Schneider, M., Raach, Y., Karypidis, P., Roux, J., Perdikaris, G., Holdermann, S., Kulsvehagen, L., Lecourt, A.-C., Narr, K., et al. (2026). Myelin antigen capture in the CNS by B cells expressing EBV latent membrane protein 1 leads to demyelinating lesion formation. Cell 189, 603-619.e25. 10.1016/j.cell.2025.12.031.

14. Wang, J., Qiu, Y., Marti, Z., Li, F., Wacker, M., Oldrati, P., Mühlenbruch, L., Jin, L., Zhang, H., Xu, W., et al. (2026). EBV infection and HLA-DR15 jointly drive multiple sclerosis by myelin peptide presentation. Cell 189, 569-584.e14. 10.1016/j.cell.2025.12.046.

15. Münz, C. (2025). Epstein–Barr virus pathogenesis and emerging control strategies. Nat. Rev. Microbiol. 23, 667– 679. 10.1038/s41579-025-01181-y.

16. Wahbeh, F., and Sabatino, J.J. (2025). Epstein-Barr Virus in Multiple Sclerosis: Past, Present, and Future. Neurol. Neuroimmunol. Neuroinflammation 12, e200460. 10.1212/NXI.0000000000200460.

17. Robinson, W.H., Younis, S., Love, Z.Z., Steinman, L., and Lanz, T. V (2024). Epstein-Barr virus as a potentiator of autoimmune diseases. Nat. Rev. Rheumatol. 20, 729–740. 10.1038/s41584-024-01167-9.

18. Babbe, H., Roers, A., Waisman, A., Lassmann, H., Goebels, N., Hohlfeld, R., Friese, M., Schröder, R., Deckert, M., Schmidt, S., et al. (2000). Clonal expansions of CD8+ T cells dominate the T cell infiltrate in active multiple sclerosis lesions as shown by micromanipulation and single cell polymerase chain reaction. J. Exp. Med. 192, 393–404. 10.1084/jem.192.3.393.

19. Jacobsen, M., Cepok, S., Quak, E., Happel, M., Gaber, R., Ziegler, A., Schock, S., Oertel, W.H., Sommer, N., and Hemmer, B. (2002). Oligoclonal expansion of memory CD8+ T cells in cerebrospinal fluid from multiple sclerosis patients. Brain 125, 538–550. 10.1093/brain/awf059.

20. Schneider-Hohendorf, T., Gerdes, L.A., Pignolet, B., Gittelman, R., Ostkamp, P., Rubelt, F., Raposo, C., Tackenberg, B., Riepenhausen, M., Janoschka, C., et al. (2022). Broader Epstein-Barr virus-specific T cell receptor repertoire in patients with multiple sclerosis. J. Exp. Med. 219, e20220650. 10.1084/jem.20220650.

21. Gate, D., Saligrama, N., Leventhal, O., Yang, A.C., Unger, M.S., Middeldorp, J., Chen, K., Lehallier, B., Channappa, D., De Los Santos, M.B., et al. (2020). Clonally expanded CD8 T cells patrol the cerebrospinal fluid in Alzheimer’s disease. Nature 577, 399–404. 10.1038/s41586-019-1895-7.

22. Hayashi, F., Mittl, K., Dandekar, R., Gerdts, J., Hassan, E., Schubert, R.D., Oshiro, L., Loudermilk, R., Greenfield, A., Augusto, D.G., et al. (2026). Antigen specificity of clonally enriched CD8+ T cells in multiple sclerosis. Nat. Immunol. 27, 490–502. 10.1038/s41590-025-02412-3.

23. Eisenbarth, S.C., Batista, F., Cyster, J., Elsner, R., Kelsoe, G., Lund, F.E., Pillai, S., Sanz, I., Shlomchik, M., Toellner, K.M., et al. (2025). A roadmap for defining “extrafollicular” B cell responses. Immunity 58, 2627–2645. 10.1016/j.immuni.2025.08.007.

24. Vietzen, H., Kühner, L.M., Berger, S.M., Ponleitner, M., Graninger, M., Pistorius, C., Jungbauer, C., Reindl, M., Saucke, H., Kauth, F., et al. (2025). Early identification of individuals at risk for multiple sclerosis by quantification of EBNA-1381-452-specific antibody titers. Nat. Commun. 16, 6416. 10.1038/s41467-025-61751-9.

25. Ostkamp, P., Deffner, M., Schulte-Mecklenbeck, A., Wünsch, C., Lu, I.-N., Wu, G.F., Goelz, S., De Jager, P.L., Kuhlmann, T., Gross, C.C., et al. (2022). A single-cell analysis framework allows for characterization of CSF leukocytes and their tissue of origin in multiple sclerosis. Sci. Transl. Med. 14, eadc9778. 10.1126/scitranslmed.adc9778.

26. Mackay, L.K., Minnich, M., Kragten, N.A.M., Liao, Y., Nota, B., Seillet, C., Zaid, A., Man, K., Preston, S., Freestone, D., et al. (2016). Hobit and Blimp1 instruct a universal transcriptional program of tissue residency in lymphocytes. Science 352, 459–463. 10.1126/science.aad2035.

27. Gray, J.I., and Farber, D.L. (2022). Tissue-Resident Immune Cells in Humans. Annu. Rev. Immunol. 40, 195– 220. 10.1146/annurev-immunol-093019-112809.

28. Kendirli, A., de la Rosa, C., Lämmle, K.F., Eglseer, K., Bauer, I.J., Kavaka, V., Winklmeier, S., Zhuo, L., Wichmann, C., Gerdes, L.A., et al. (2023). A genome-wide in vivo CRISPR screen identifies essential regulators of T cell migration to the CNS in a multiple sclerosis model. Nat. Neurosci. 26, 1713–1725. 10.1038/s41593-023-01432-2.

29. Chiu, C., McCausland, M., Sidney, J., Duh, F.-M., Rouphael, N., Mehta, A., Mulligan, M., Carrington, M., Wieland, A., Sullivan, N.L., et al. (2014). Broadly reactive human CD8 T cells that recognize an epitope conserved between VZV, HSV and EBV. PLoS Pathog. 10, e1004008. 10.1371/journal.ppat.1004008.

30. Schlott, F., Steubl, D., Ameres, S., Moosmann, A., Dreher, S., Heemann, U., Hösel, V., Busch, D.H., and Neuenhahn, M. (2018). Characterization and clinical enrichment of HLA-C*07:02-restricted Cytomegalovirus-specific CD8+ T cells. PLoS One 13, e0193554. 10.1371/journal.pone.0193554.

31. Goncharov, M., Bagaev, D., Shcherbinin, D., Zvyagin, I., Bolotin, D., Thomas, P.G., Minervina, A.A., Pogorelyy, M. V., Ladell, K., McLaren, J.E., et al. (2022). VDJdb in the pandemic era: a compendium of T cell receptors specific for SARS-CoV-2. Nat. Methods 19, 1017–1019. 10.1038/s41592-022-01578-0.

32. Schattgen, S., Vegesana, K., Hazelton, W.D., Minervina, A., Valkiers, S., Slowikowski, K., Smith, N., Villani, A.-C., Thomas, P.G., and Bradley, P. (2025). Diverse modes of T cell receptor sequence convergence define unique functional and cellular phenotypes. bioRxiv. 10.1101/2025.05.31.657155.

33. Pulvirenti, N., Righetti, C., Clemente, F., Serafini, B., Cordiglieri, C., Vasco, C., Maioli, S., Pietroboni, A., Galeota, E., Rosicarelli, B., et al. (2025). Regulatory T-cells in multiple sclerosis produce IL-10 in the central nervous system but are activated by Epstein-Barr Virus. bioRxiv. 10.1101/2024.07.30.605745.

34. Bonnal, R.J.P., Rossetti, G., Lugli, E., De Simone, M., Gruarin, P., Brummelman, J., Drufuca, L., Passaro, M., Bason, R., Gervasoni, F., et al. (2021). Clonally expanded EOMES+ Tr1-like cells in primary and metastatic tumors are associated with disease progression. Nat. Immunol. 22, 735–745. 10.1038/s41590-021-00930-4.

35. Giladi, A., Cohen, M., Medaglia, C., Baran, Y., Li, B., Zada, M., Bost, P., Blecher-Gonen, R., Salame, T.M., Mayer, J.U., et al. (2020). Dissecting cellular crosstalk by sequencing physically interacting cells. Nat. Biotechnol. 38, 629–637. 10.1038/s41587-020-0442-2.

36. Burel, J.G., Pomaznoy, M., Lindestam Arlehamn, C.S., Weiskopf, D., da Silva Antunes, R., Jung, Y., Babor, M., Schulten, V., Seumois, G., Greenbaum, J.A., et al. (2019). Circulating T cell-monocyte complexes are markers of immune perturbations. Elife 8, e46045. 10.7554/eLife.46045.

37. Jelcic, I., Al Nimer, F., Wang, J., Lentsch, V., Planas, R., Jelcic, I., Madjovski, A., Ruhrmann, S., Faigle, W., Frauenknecht, K., et al. (2018). Memory B Cells Activate Brain-Homing, Autoreactive CD4+ T Cells in Multiple Sclerosis. Cell 175, 85-100.e23. 10.1016/j.cell.2018.08.011.

38. Ochs, J., Nissimov, N., Torke, S., Freier, M., Grondey, K., Koch, J., Klein, M., Feldmann, L., Gudd, C., Bopp, T., et al. (2022). Proinflammatory CD20+ T cells contribute to CNS-directed autoimmunity. Sci. Transl. Med. 14, eabi4632. 10.1126/scitranslmed.abi4632.

39. Morimoto, K., Baba, Y., Shinohara, H., Kang, S., Nojima, S., Kimura, T., Ito, D., Yoshida, Y., Maeda, Y., Sarashina-Kida, H., et al. (2016). LRRK1 is critical in the regulation of B-cell responses and CARMA1-dependent NF-κB activation. Sci. Rep. 6, 25738. 10.1038/srep25738.

40. Wang, S., Han, H., Qian, Y., Ruan, X., Lin, Z., Li, J., Chen, B., Lai, Y., Wang, Z., Li, M., et al. (2024). Aberrant METTL1-mediated tRNA m7G modification alters B-cell responses in systemic autoimmunity in humans and mice. Nat. Commun. 15, 10599. 10.1038/s41467-024-54941-4.

41. Jahrsdörfer, B., Vollmer, A., Blackwell, S.E., Maier, J., Sontheimer, K., Beyer, T., Mandel, B., Lunov, O., Tron, K., Ulrich Nienhaus, G., et al. (2010). Granzyme B produced by human plasmacytoid dendritic cells suppresses T-cell expansion. Blood 115, 1156–1165. 10.1182/blood-2009-07-235382.

42. Lerma-Martin, C., Badia-I-Mompel, P., Ramirez Flores, R.O., Sekol, P., Schäfer, P.S.L., Riedl, C.J., Hofmann, A., Thäwel, T., Wünnemann, F., Ibarra-Arellano, M.A., et al. (2024). Cell type mapping reveals tissue niches and interactions in subcortical multiple sclerosis lesions. Nat. Neurosci. 27, 2354–2365. 10.1038/s41593-024-01796-z.

43. Schroeder, A., Loth, M.L., Luo, C., Yao, S., Yan, H., Zhang, D., Piya, S., Plowey, E., Hu, W., Clemenceau, J.R., et al. (2025). Scaling up spatial transcriptomics for large-sized tissues: uncovering cellular-level tissue architecture beyond conventional platforms with iSCALE. Nat. Methods 22, 1911–1922. 10.1038/s41592-025-02770-8.

44. Herich, S., Schneider-Hohendorf, T., Rohlmann, A., Khaleghi Ghadiri, M., Schulte-Mecklenbeck, A., Zondler, L., Janoschka, C., Ostkamp, P., Richter, J., Breuer, J., et al. (2019). Human CCR5high effector memory cells perform CNS parenchymal immune surveillance via GZMK-mediated transendothelial diapedesis. Brain 142, 3411– 3427. 10.1093/brain/awz301.

45. Balashov, K.E., Rottman, J.B., Weiner, H.L., and Hancock, W.W. (1999). CCR5+ and CXCR3+ T cells are increased in multiple sclerosis and their ligands MIP-1α and IP-10 are expressed in demyelinating brain lesions. Proc. Natl. Acad. Sci. U. S. A. 96, 6873–6878. 10.1073/pnas.96.12.6873.

46. Chen, K.Y., De Giovanni, M., Xu, Y., An, J., Kirthivasan, N., Lu, E., Jiang, K., Brooks, S., Ranucci, S., Yang, J., et al. (2025). Inflammation switches the chemoattractant requirements for naive lymphocyte entry into lymph nodes. Cell 188, 1019-1035.e22. 10.1016/j.cell.2024.11.031.

47. Donado, C.A., Theisen, E., Zhang, F., Nathan, A., Fairfield, M.L., Rupani, K.V., Jones, D., Johannes, K.P., Accelerating Medicines Partnership RA/SLE Network, Raychaudhuri, S., et al. (2025). Granzyme K activates the entire complement cascade. Nature 641, 211–221. 10.1038/s41586-025-08713-9.

48. Lan, F., Li, J., Miao, W., Sun, F., Duan, S., Song, Y., Yao, J., Wang, X., Wang, C., Liu, X., et al. (2025). GZMK-expressing CD8+ T cells promote recurrent airway inflammatory diseases. Nature 638, 490–498. 10.1038/s41586-024-08395-9.

49. Cooper, D.M., Pechkovsky, D. V, Hackett, T.L., Knight, D.A., and Granville, D.J. (2011). Granzyme K activates protease-activated receptor-1. PLoS One 6, e21484. 10.1371/journal.pone.0021484.

50. Kaiserman, D., Zhao, P., Rowe, C.L., Leong, A., Barlow, N., Joeckel, L.T., Hitchen, C., Stewart, S.E., Hollenberg, M.D., Bunnett, N., et al. (2022). Granzyme K initiates IL-6 and IL-8 release from epithelial cells by activating protease-activated receptor 2. PLoS One 17, e0270584. 10.1371/journal.pone.0270584.

51. Thomas, O.G., Rykaczewska, U., Galešić, M., van der Burgt, R.T.M., Hallén, N., Ferro, F., Bronge, M., Marti, Z., Li, Y., Riqué, A.H., et al. (2026). Anoctamin-2-specific T cells link Epstein-Barr virus to multiple sclerosis. Cell 189, 585-602.e38. 10.1016/j.cell.2025.12.032.

52. Lanz, T. V., Brewer, R.C., Ho, P.P., Moon, J.S., Jude, K.M., Fernandez, D., Fernandes, R.A., Gomez, A.M., Nadj, G.S., Bartley, C.M., et al. (2022). Clonally expanded B cells in multiple sclerosis bind EBV EBNA1 and GlialCAM. Nature 603, 321–327. 10.1038/s41586-022-04432-7.

53. Sato, W., Tomita, A., Ichikawa, D., Lin, Y., Kishida, H., Miyake, S., Ogawa, M., Okamoto, T., Murata, M., Kuroiwa, Y., et al. (2012). CCR2+CCR5+ T Cells Produce Matrix Metalloproteinase-9 and Osteopontin in the Pathogenesis of Multiple Sclerosis. J. Immunol. 189, 5057–5065. 10.4049/jimmunol.1202026.

54. Niessl, J., Müller, T.R., Constantz, C., Cai, C., Nilsén, V., Rivera Ballesteros, O., Adamo, S., Kammann, T., Mouchtaridi, E., Gao, Y., et al. (2025). Tissue origin and virus specificity shape human CD8+ T cell cytotoxicity. Sci. Immunol. 10, eadq4881. 10.1126/sciimmunol.adq4881.

55. Fransen, N.L., Hsiao, C.C., Van Der Poel, M., Engelenburg, H.J., Verdaasdonk, K., Vincenten, M.C.J., Remmerswaal, E.B.M., Kuhlmann, T., Mason, M.R.J., Hamann, J., et al. (2020). Tissue-resident memory T cells invade the brain parenchyma in multiple sclerosis white matter lesions. Brain 143, 1714–1730. 10.1093/brain/awaa117.

56. Farooq, R., Cutler, B., Pisa, M., Balaram Kuttikkatte, S., Palace, J., DeLuca, G.C., Gold, R., Theis, F.J., Fugger, L., and Attfield, K.E. (2026). Multimodal discovery of a pathogenic B cell-dependent T cell state in multiple sclerosis. Brain, Epub ahead of print. 10.1093/brain/awag303.

57. Wu, C., Jiang, S., Chen, Z., Li, T., Gu, X., Dai, M., Du, F., Ye, Y., Tang, L., Wang, M., et al. (2025). Single-cell transcriptomics reveal potent extrafollicular B cell response linked with granzyme K+ CD8 T cell activation in lupus kidney. Ann. Rheum. Dis. 84, 451–466. 10.1136/ard-2024-225876.

58. Xiao, L., Jia, L., Bai, L., He, L., Yang, B., Wu, C., and Li, H. (2016). Phenotypic and functional characteristics of IL-21-expressing CD8(+) T cells in human nasal polyps. Sci. Rep. 6, 30362. 10.1038/srep30362.

59. Higashioka, K., Yoshimura, M., Sakuragi, T., Ayano, M., Kimoto, Y., Mitoma, H., Ono, N., Arinobu, Y., Kikukawa, M., Yamada, H., et al. (2021). Human PD-1hiCD8+ T Cells Are a Cellular Source of IL-21 in Rheumatoid Arthritis. Front. Immunol. 12, 654623. 10.3389/fimmu.2021.654623.

60. Shen, J., Luo, X., Wu, Q., Huang, J., Xiao, G., Wang, L., Yang, B., Li, H., and Wu, C. (2018). A Subset of CXCR5+CD8+ T Cells in the Germinal Centers From Human Tonsils and Lymph Nodes Help B Cells Produce Immunoglobulins. Front. Immunol. 9, 2287. 10.3389/fimmu.2018.02287.

61. Yang, R., Masters, A.R., Fortner, K.A., Champagne, D.P., Yanguas-Casás, N., Silberger, D.J., Weaver, C.T., Haynes, L., and Rincon, M. (2016). IL-6 promotes the differentiation of a subset of naive CD8+ T cells into IL-21–producing B helper CD8+ T cells. J. Exp. Med. 213, 2281–2291. 10.1084/jem.20160417.

62. Younis, S., Moutusy, S.I., Rasouli, S., Jahanbani, S., Pandit, M., Wu, X., Acharya, S., Sharpe, O., Wijeratne, T.U., Harris, M.L., et al. (2025). Epstein-Barr virus reprograms autoreactive B cells as antigen-presenting cells in systemic lupus erythematosus. Sci. Transl. Med. 17, eady0210. 10.1126/scitranslmed.ady0210.

63. Younis, S., Rasouli, S., Loeffler, J.W., Sattarnezhad, N., Courtney, Y., Moutusy, S.I., Jahanbani, S., Pandit, M., Tomczak, A., Wong, H.H., et al. (2026). EBV reprograms autoreactive anti-CNS B cells as antigen presenting cells in multiple sclerosis. bioRxiv. 10.64898/2026.02.11.701910.

64. Orr, N., and Steinman, L. (2025). Epstein-Barr virus and the immune microenvironment in multiple sclerosis: Insights from high-dimensional brain tissue imaging. Proc. Natl. Acad. Sci. U. S. A. 122, e2425670122. 10.1073/pnas.2425670122.

65. Bjornevik, K., Mahler, J.V., Bilodeau, P.A., Rød, B.E., Romanow, G., Mikami, T., Anderson, M., Bobrowski-Khoury, N., Zhu, H., Nguyen, J., et al. (2026). CD4+ T cells reactive to Epstein-Barr virus late lytic antigens are enriched in individuals with multiple sclerosis. Sci. Transl. Med. 18, eadz6566. 10.1126/scitranslmed.adz6566.

66. Sarkkinen, J., Yohannes, D.A., Kreivi, N., Dürnsteiner, P., Elsakova, A., Huuhtanen, J., Nowlan, K., Kurdo, G., Linden, R., Saarela, M., et al. (2025). Altered immune landscape of cervical lymph nodes reveals Epstein-Barr virus signature in multiple sclerosis. Sci. Immunol. 10, eadl3604. 10.1126/sciimmunol.adl3604.

67. Xie, T., Du, Y., Wang, Q., Zhang, H., Wei, K., Chi, X., Bai, X., Fu, Y., Peng, Z., Zhu, Y., et al. (2026). A unique CD4+ T cell subset expressing granzyme K is regulated by transcription factor EOMES and important for T cell-mediated intestinal inflammation. Nat. Immunol. 27, 1212– 1223. 10.1038/s41590-026-02479-6.

68. Nyeo, S.S., Cumming, E.M., Burren, O.S., Pagadala, M.S., Gutierrez, J.C., Ali, T.A., Kida, L.C., Chen, Y., Chu, H., Hu, F., et al. (2026). Population-scale sequencing resolves determinants of persistent EBV DNA. Nature 650, 664–672. 10.1038/s41586-025-10020-2.

69. Schmidt, A., Alawathurage, T.M., David, F.S., Ogawa, Y., Frach, L., Richter, S., Schaefer, M., Mathey, C.M., Henne, S.K., Japan COVID-19 Task Force, et al. (2026). Host control of persistent Epstein-Barr virus infection. Nature 653, 444–454. 10.1038/s41586-026-10274-4.

70. Yasumizu, Y., Kim, N., Rivier, C.A., Moon, J., Kojima, S., Chen, H.-L., Buitrago-Pocasangre, N., Silberfeld, A., Quinn, E., Vaughn, S., et al. (2025). A Genetically Driven Immunologic Mechanism Underlying the Link between EBV and Multiple Sclerosis. medRxiv. 10.64898/2025.12.11.25342083.

71. Mogilenko, D.A., Shpynov, O., Andhey, P.S., Arthur, L., Swain, A., Esaulova, E., Brioschi, S., Shchukina, I., Kerndl, M., Bambouskova, M., et al. (2021). Comprehensive Profiling of an Aging Immune System Reveals Clonal GZMK+ CD8+ T Cells as Conserved Hallmark of Inflammaging. Immunity 54, 99-115.e12. 10.1016/j.immuni.2020.11.005.

72. Wolock, S.L., Lopez, R., and Klein, A.M. (2019). Scrublet: Computational Identification of Cell Doublets in Single-Cell Transcriptomic Data. Cell Syst. 8, 281-291.e9. 10.1016/j.cels.2018.11.005.

73. Bernstein, N.J., Fong, N.L., Lam, I., Roy, M.A., Hendrickson, D.G., and Kelley, D.R. (2020). Solo: Doublet Identification in Single-Cell RNA-Seq via Semi-Supervised Deep Learning. Cell Syst. 11, 95-101.e5. 10.1016/j.cels.2020.05.010.

74. Xu, J., Falconer, C., Nguyen, Q., Crawford, J., McKinnon, B.D., Mortlock, S., Senabouth, A., Andersen, S., Chiu, H.S., Jiang, L., et al. (2019). Genotype-free demultiplexing of pooled single-cell RNA-seq. Genome Biol. 20, 290. 10.1186/s13059-019-1852-7.

75. Lopez, R., Regier, J., Cole, M.B., Jordan, M.I., and Yosef, N. (2018). Deep generative modeling for single-cell transcriptomics. Nat. Methods 15, 1053–1058. 10.1038/s41592-018-0229-2.

76. Hao, Y., Stuart, T., Kowalski, M.H., Choudhary, S., Hoffman, P., Hartman, A., Srivastava, A., Molla, G., Madad, S., Fernandez-Granda, C., et al. (2024). Dictionary learning for integrative, multimodal and scalable single-cell analysis. Nat. Biotechnol. 42, 293–304. 10.1038/s41587-023-01767-y.

77. Frischholz, S., Schuster, E.-M., Grotz, M., Schülein, C., Benz, J., Kocher, K., Klotz, L., Varga, S., Hiltner, T., Alsalameh, R., et al. (2026). Metabolic quiescence of naive-like memory T cells precedes and maintains antigen-specific T cell memory. Nat. Immunol. 27, 452–462. 10.1038/s41590-026-02421-w.

78. Povlsen, H.R., Bentzen, A.K., Kadivar, M., Jessen, L.E., Hadrup, S.R., and Nielsen, M. (2023). Improved T cell receptor antigen pairing through data-driven filtering of sequencing information from single cells. Elife 12, e81810. 10.7554/eLife.81810.

79. Rodenko, B., Toebes, M., Hadrup, S.R., van Esch, W.J.E., Molenaar, A.M., Schumacher, T.N.M., and Ovaa, H. (2006). Generation of peptide-MHC class I complexes through UV-mediated ligand exchange. Nat. Protoc. 1, 1120–1132. 10.1038/nprot.2006.121.

80. Bakker, A.H., Hoppes, R., Linnemann, C., Toebes, M., Rodenko, B., Berkers, C.R., Hadrup, S.R., Van Esch, W.J.E., Heemskerk, M.H.M., Ovaa, H., et al. (2008). Conditional MHC class I ligands and peptide exchange technology for the human MHC gene products HLA-A1, -A3, -A11, and -B7. Proc. Natl. Acad. Sci. U. S. A. 105, 3825–3830. 10.1073/pnas.0709717105.

81. Saini, S.K., Tamhane, T., Anjanappa, R., Saikia, A., Ramskov, S., Donia, M., Svane, I.M., Jakobsen, S.N., Garcia-Alai, M., Zacharias, M., et al. (2019). Empty peptide-receptive MHC class I molecules for efficient detection of antigen-specific T cells. Sci. Immunol. 4, eaau9039. 10.1126/sciimmunol.aau9039.

82. Kristensen, N.P., Dionisio, E., Bentzen, A.K., Tamhane, T., Kemming, J.S., Nos, G., Voss, L.F., Hansen, U.K., Lauer, G.M., and Hadrup, S.R. (2024). Simultaneous analysis of pMHC binding and reactivity unveils virus-specific CD8 T cell immunity to a concise epitope set. Sci. Adv. 10, eadm8951. 10.1126/sciadv.adm8951.

83. Kristensen, N.P., Heeke, C., Tvingsholm, S.A., Borch, A., Draghi, A., Crowther, M.D., Carri, I., Munk, K.K., Holm, J.S., Bjerregaard, A.-M., et al. (2022). Neoantigen-reactive CD8+ T cells affect clinical outcome of adoptive cell therapy with tumor-infiltrating lymphocytes in melanoma. J. Clin. Invest. 132, e150535. 10.1172/JCI150535.

84. Saini, S.K., Hersby, D.S., Tamhane, T., Povlsen, H.R., Amaya Hernandez, S.P., Nielsen, M., Gang, A.O., and Hadrup, S.R. (2021). SARS-CoV-2 genome-wide T cell epitope mapping reveals immunodominance and substantial CD8+ T cell activation in COVID-19 patients. Sci. Immunol. 6, eabf7550. 10.1126/sciimmunol.abf7550.

85. Vita, R., Blazeska, N., Marrama, D., Shackelford, D., Zalman, L., Foos, G., Zarebski, L., Chan, K., Reardon, B., Fitzpatrick, S., et al. (2025). The Immune Epitope Database (IEDB): 2024 update. Nucleic Acids Res. 53, D436–D443. 10.1093/nar/gkae1092.

86. An, Y., Drost, F., Bonafonte-Pardàs, I., Grotz, M., Schober, K., and Schubert, B. (2026). DextraDemixer enables accurate identification of antigen-specific T cells from pMHC multimer experiments. bioRxiv. 10.64898/2026.06.23.733339.

87. Turchaninova, M.A., Davydov, A., Britanova, O. V., Shugay, M., Bikos, V., Egorov, E.S., Kirgizova, V.I., Merzlyak, E.M., Staroverov, D.B., Bolotin, D.A., et al. (2016). High-quality full-length immunoglobulin profiling with unique molecular barcoding. Nat. Protoc. 11, 1599–1616. 10.1038/nprot.2016.093.

88. Bolotin, D.A., Poslavsky, S., Mitrophanov, I., Shugay, M., Mamedov, I.Z., Putintseva, E. V, and Chudakov, D.M. (2015). MiXCR: software for comprehensive adaptive immunity profiling. Nat. Methods 12, 380–381. 10.1038/nmeth.3364.

89. Absinta, M., Maric, D., Gharagozloo, M., Garton, T., Smith, M.D., Jin, J., Fitzgerald, K.C., Song, A., Liu, P., Lin, J.-P., et al. (2021). A lymphocyte-microglia-astrocyte axis in chronic active multiple sclerosis. Nature 597, 709–714. 10.1038/s41586-021-03892-7.

