## Supplementary Figures for "A pro-inflammatory effector-memory T cell program links peripheral EBV priming to CNS autoimmunity"

[1] Mikrobiologisches Institut – Klinische Mikrobiologie, Immunologie und Hygiene, Universitätsklinikum Erlangen, Friedrich-Alexander Universität Erlangen-Nürnberg, Erlangen, Germany; [2] Department of Neurology, University Hospital Erlangen, Friedrich-Alexander University Erlangen-Nürnberg, Germany; [3] Department of Neurosurgery, University Hospital Erlangen, Friedrich-Alexander-Universität Erlangen-Nürnberg, Erlangen, Germany; [4] Department of Health Technology, Technical University of Denmark, Kongens Lyngby, Denmark; [5] Department of Host–Microbe Interactions, St. Jude Children's Research Hospital, Memphis, TN, United States; [6] Department of Internal Medicine 5 - Hematology and Oncology, Friedrich-Alexander-Universität Erlangen-Nürnberg (FAU) and Universitätsklinikum Erlangen; Erlangen, Germany; [7] Department of Nephropathology, Institute of Pathology, Experimental Renal and Cardiovascular Research, Friedrich-Alexander-Universität Erlangen-Nürnberg (FAU), Erlangen, Germany; [8] Department of Cardiology, Friedrich-Alexander-Universität Erlangen-Nürnberg (FAU), Erlangen, Germany; [9] Health Campus for Inflammation, Immunity and Infection (GCI3), Otto von Guericke University of Magdeburg, Magdeburg, Germany; [10] Department of Hematology, Oncology and Cell Therapy, Otto von Guericke University of Magdeburg, Magdeburg, Germany; [11] Department of Neurology, Otto von Guericke University, Magdeburg, Germany; [12] German Center for Neurodegenerative Diseases (DZNE), Magdeburg, Germany; [13] Department of Translational Immunology, Department of Medicine 3-Rheumatology and Immunology, Uniklinikum Erlangen und Friedrich-Alexander-Universität (FAU) Erlangen-Nürnberg, Nikolaus-Fiebiger-Center, Erlangen, Germany; [14] Deutsches Zentrum Immuntherapie, Friedrich-Alexander-Universität (FAU) Erlangen-Nürnberg and Uniklinikum Erlangen, 91054 Erlangen, Germany; [15] Immunology and Vaccine Development Program, Vaccine and Infectious Disease Division, Fred Hutchinson Cancer Center, Seattle, WA, United States; [16] FAU Profile Center Immunomedicine, FAU Erlangen-Nürnberg, Erlangen, Germany; corresponding author: Kilian Schober;

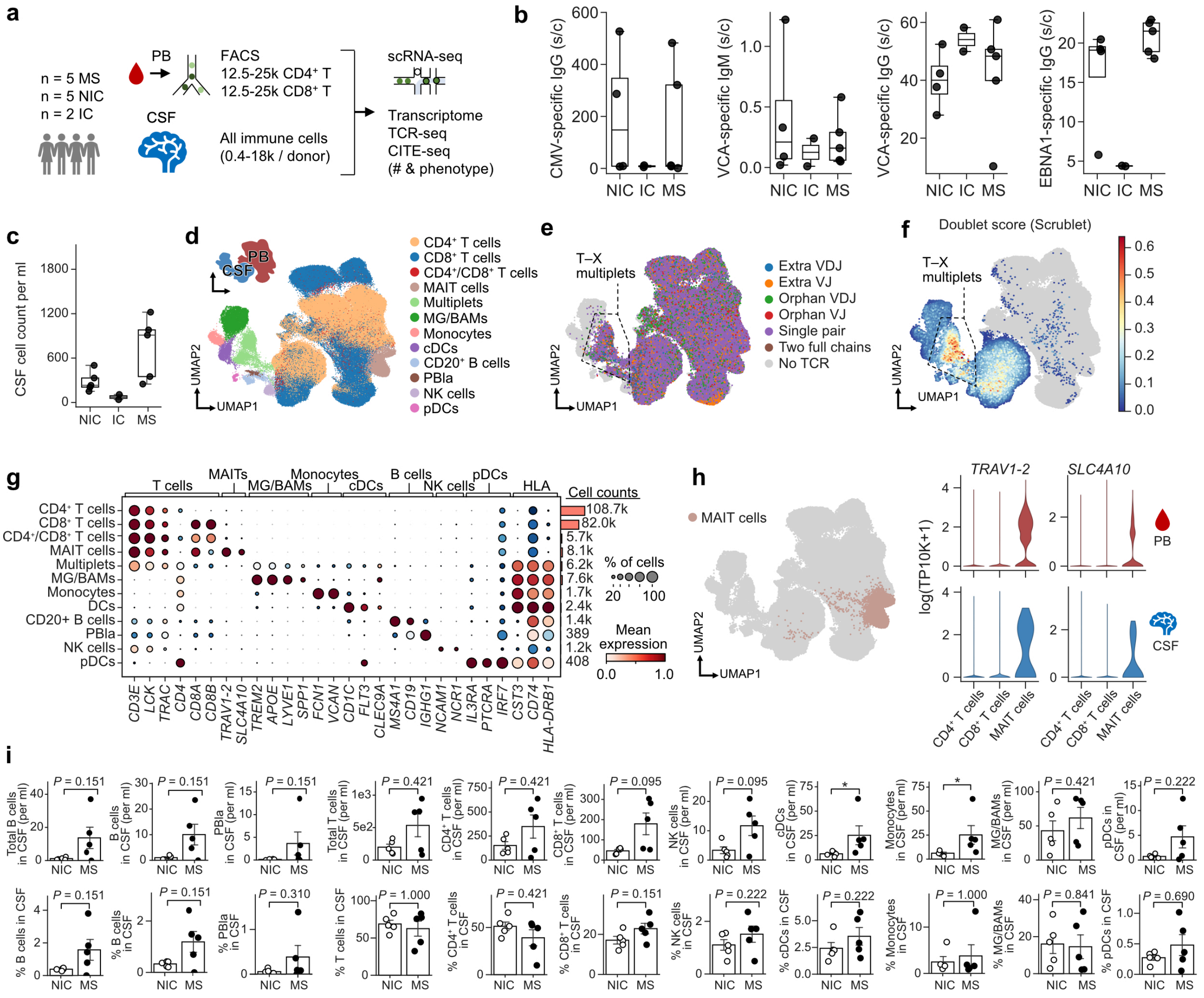

**Supplementary figure 1: Single-cell transcriptomic analysis of leukocytes from paired CSF and PB samples.** (a) Experimental setup. Five NIC, five MS and two IC were prospectively sampled after informed consent during diagnostic or therapeutic procedures involving a lumbar puncture. Suspected MS cases were subsequently confirmed. Cryopreserved PBMCs from PB and fresh CSF samples were compared using multimodal single-cell sequencing (10x Genomics) combining scRNA-seq, scTCR-seq, and CITE-seq. (b) Serological summary data. Four donors were CMV-positive. All donors were EBV-positive. (c) CSF leukocyte counts per mL of CSF. (d) UMAP and manually annotated clusters for the collective PB/CSF dataset. (e) Expression of recombined TCR alpha (TRA) and TCR beta (TRB) chains as measured by scTCR-seq (10x Genomics). T–X multiplets represent T cells interacting with different non-T cells. (f) Doublet scores measured by Scrublet indicative of T–X multiplets. (g) Summary of marker genes associated with each manually annotated phenotypic cluster including T–X multiplets. (h) Presence of MAIT cells in CSF as detected by sharing of TCRs found among MAIT cell clusters in PB. (i) Quantification of cells per mL and percentage of cells using reclustered CSF data and further annotated Leiden clusters detailed in Fig. 3a. *P* values were calculated using a Wilcoxon test. *, $P < 0.05$.

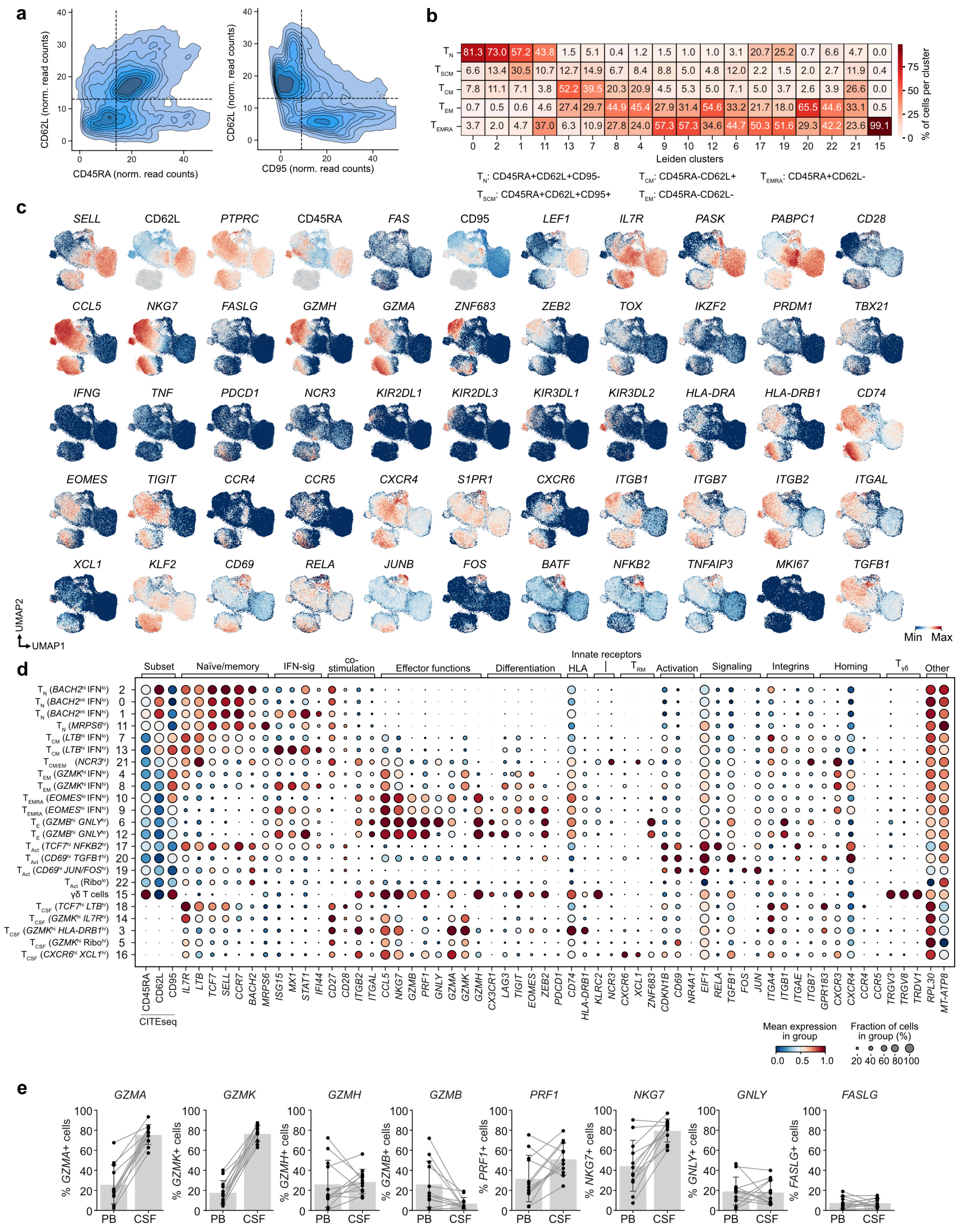

a
CD62L (norm. read counts)
CD45RA (norm. read counts)
CD95 (norm. read counts)
b
TN 81.3 73.0 57.2 43.8 1.5 5.1 0.4 1.2 1.5 1.0 1.0 3.1 20.7 25.2 0.7 6.6 4.7 0.0
TSCM 6.6 13.4 30.5 10.7 12.7 14.9 6.7 8.4 8.8 5.0 4.8 12.0 2.2 1.5 2.0 2.7 11.9 0.4
TCM 7.8 11.1 7.1 3.8 52.2 39.5 20.3 20.9 4.5 5.3 5.0 7.1 5.0 3.7 2.6 3.9 26.6 0.0
TEM 0.7 0.5 0.6 4.6 27.4 29.7 44.9 45.4 27.9 31.4 54.6 33.2 21.7 18.0 65.5 44.6 33.1 0.5
TEMRA 3.7 2.0 4.7 37.0 6.3 10.9 27.8 24.0 57.3 57.3 34.6 44.7 50.3 51.6 29.3 42.2 23.6 99.1
0 2 1 11 13 7 8 4 9 10 12 6 17 19 20 22 21 15
Leiden clusters
% of cells per cluster
TN: CD45RA+CD62L+CD95-
TSCM: CD45RA+CD62L+CD95+
TCM: CD45RA-CD62L+
TEM: CD45RA-CD62L-
TEMRA: CD45RA+CD62L-
c
SELL CD62L PTPRC CD45RA FAS CD95 LEF1 IL7R PASK PABPC1 CD28
CCL5 NKG7 FASLG GZMH GZMA ZNF683 ZEB2 TOX IKZF2 PRDM1 TBX21
IFNG TNF PDCD1 NCR3 KIR2DL1 KIR2DL3 KIR3DL1 KIR3DL2 HLA-DRA HLA-DRB1 CD74
EOMES TIGIT CCR4 CCR5 CXCR4 S1PR1 CXCR6 ITGB1 ITGB7 ITGB2 ITGAL
XCL1 KLF2 CD69 RELA JUNB FOS BATF NFKB2 TNFAIP3 MKI67 TGFB1
UMAP2
UMAP1
Min Max
d
Subset
Naïve/memory
IFN-sig
co-stimulation
Effector functions
Differentiation
Innate receptors
HLA
TRM
Activation
Signaling
Integrins
Homing
Tγδ
Other
TN (BACH2hi IFNlo) 2
TN (BACH2int IFNlo) 0
TN (BACH2int IFNhi) 1
TN (MRPS6hi) 11
TCM (LTBhi IFNlo) 7
TCM (LTBhi IFNhi) 13
TCM/EM (NCR3hi) 21
TEM (GZMKhi IFNlo) 4
TEM (GZMKhi IFNhi) 8
TEMRA (EOMESlo IFNlo) 10
TEMRA (EOMEShi IFNhi) 9
TE (GZMBhi GNLYhi) 6
TE (GZMBhi GNLYlo) 12
TAct (TCF7hi NFKB2hi) 17
TAct (CD69hi TGFB1hi) 20
TAct (CD69hi JUN/FOShi) 19
TAct (Ribolo) 22
γδ T cells 15
TCSF (TCF7hi LTBhi) 18
TCSF (GZMKhi IL7Rhi) 14
TCSF (GZMKhi HLA-DRB1hi) 3
TCSF (GZMKhi Ribohi) 5
TCSF (CXCR6hi XCL1hi) 16
CD45RA CD62L CD95 IL7R LTB TCF7 SELL CCR7 BACH2 MRPS6 ISG15 MX1 STAT1 IFI44 CD27 CD28 ITGB2 ITGAL CCL5 NKG7 GZMB PRF1 GNLY GZMA GZMK GZMH CX3CR1 LAG3 TIGIT EOMES ZEB2 PDCD1 CD74 HLA-DRB1 KLRC2 NCR3 CXCR6 XCL1 ZNF683 CDKN1B CD69 NR4A1 EIF1 RELA TGFB1 FOS JUN ITGA4 ITGB1 ITGAE ITGB7 GPR183 CXCR3 CXCR4 CCR4 CCR5 TRGV3 TRGV8 TRDV1 RPL30 MT-ATP8
CITEseq
Mean expression in group
0.0 0.5 1.0
Fraction of cells in group (%)
20 40 60 80 100
e
GZMA GZMK GZMH GZMB PRF1 NKG7 GNLY FASLG
% GZMA+ cells
% GZMK+ cells
% GZMH+ cells
% GZMB+ cells
% PRF1+ cells
% NKG7+ cells
% GNLY+ cells
% FASLG+ cells
0 20 40 60 80 100
PB CSF

**Supplementary figure 2: Phenotypic characterization and annotation of CSF and PB CD8$^+$ T cells.** (a) Normalized CITE-seq count values displayed as a density plot. Threshold values to define virtual gates for $T_N$, $T_{SCM}$, $T_{CM}$, $T_{EM}$, and $T_{EMRA}$ are indicated as dashed lines. (b) Quantification and annotation of Leiden clusters according to the virtual gates shown in (a). (c) Representative UMAPs for selected genes related to CD8$^+$ T cell differentiation, phenotype and function. (d) Quantification of representative genes and manual annotation of Leiden clusters. (e) Frequency of transcript-positive cells in PB and CSF per donor. Five MS and five NIC are shown. Genes were manually curated based on their association with CD8$^+$ T cell effector functions.

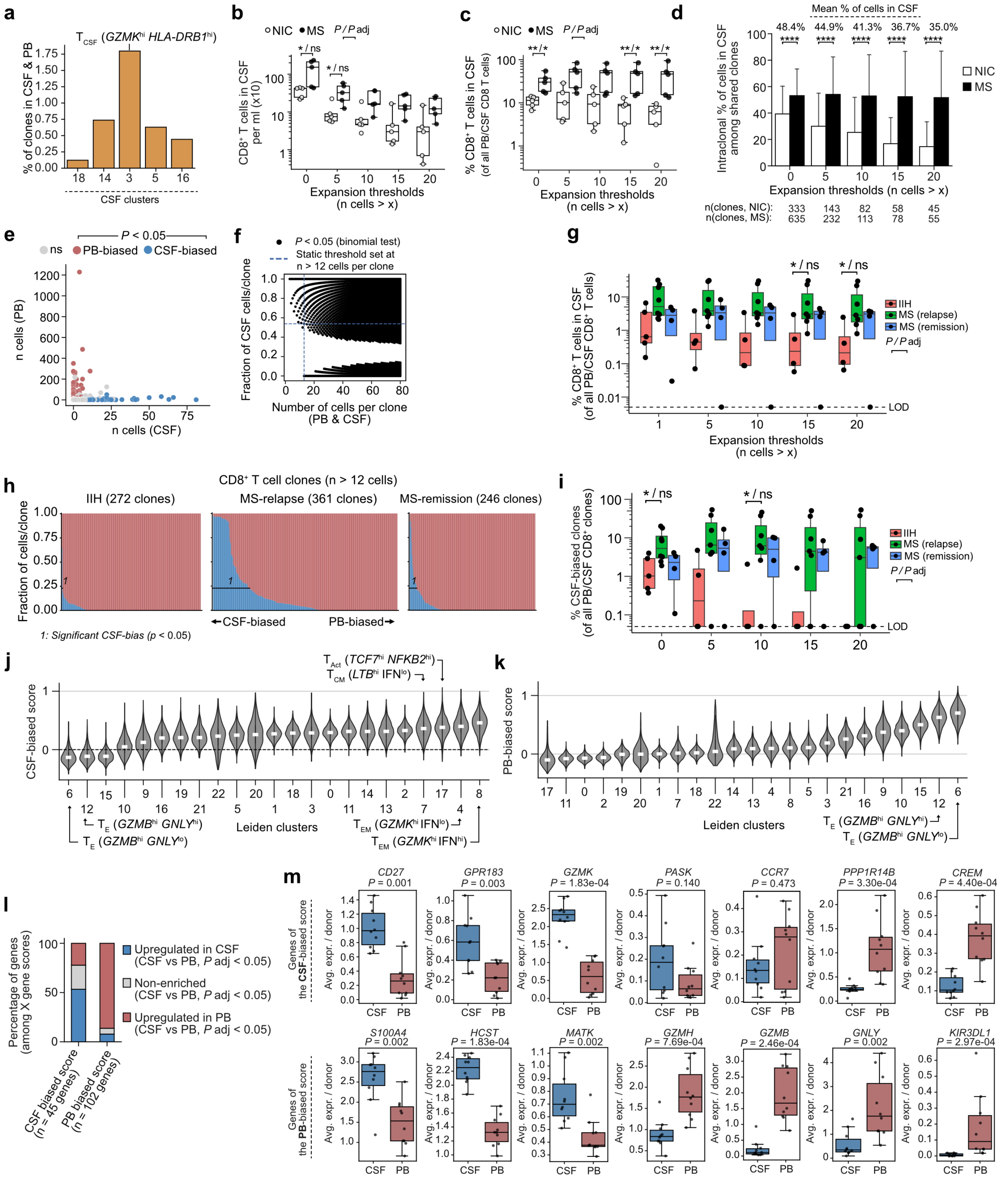

a
b
c
d
e
f
g
h
i
j
k
l
m
$T_{CSF}$ (GZMK$^{hi}$ HLA-DRB1$^{hi}$)
% of clones in CSF & PB
CSF clusters
NIC
MS
P / P adj
* / ns
CD8+ T cells in CSF per ml (x10)
Expansion thresholds (n cells > x)
** / *
% CD8+ T cells in CSF (of all PB/CSF CD8 T cells)
Mean % of cells in CSF
48.4%
44.9%
41.3%
36.7%
35.0%
****
Intraclonal % of cells in CSF among shared clones
n(clones, NIC): 333 143 82 58 45
n(clones, MS): 635 232 113 78 55
P < 0.05
ns
PB-biased
CSF-biased
n cells (PB)
n cells (CSF)
P < 0.05 (binomial test)
Static threshold set at n > 12 cells per clone
Fraction of CSF cells/clone
Number of cells per clone (PB & CSF)
IIH
MS (relapse)
MS (remission)
% CD8+ T cells in CSF (of all PB/CSF CD8+ T cells)
LOD
CD8+ T cell clones (n > 12 cells)
IIH (272 clones)
MS-relapse (361 clones)
MS-remission (246 clones)
Fraction of cells/clone
←CSF-biased
PB-biased→
1: Significant CSF-bias (p < 0.05)
% CSF-biased clones (of all PB/CSF CD8+ clones)
$T_{Act}$ (TCF7$^{hi}$ NFKB2$^{hi}$)
$T_{CM}$ (LTB$^{hi}$ IFN$^{lo}$)
CSF-biased score
$T_{E}$ (GZMB$^{hi}$ GNLY$^{hi}$)
$T_{E}$ (GZMB$^{hi}$ GNLY$^{lo}$)
Leiden clusters
$T_{EM}$ (GZMK$^{hi}$ IFN$^{lo}$)
$T_{EM}$ (GZMK$^{hi}$ IFN$^{hi}$)
PB-biased score
Percentage of genes (among X gene scores)
Upregulated in CSF (CSF vs PB, P adj < 0.05)
Non-enriched (CSF vs PB, P adj < 0.05)
Upregulated in PB (CSF vs PB, P adj < 0.05)
CSF biased score (n = 45 genes)
PB biased score (n = 102 genes)
Genes of the CSF-biased score
Genes of the PB-biased score
Avg. expr. / donor
CSF
PB
CD27 P = 0.001
GPR183 P = 0.003
GZMK P = 1.83e-04
PASK P = 0.140
CCR7 P = 0.473
PPP1R14B P = 3.30e-04
CREM P = 4.40e-04
S100A4 P = 0.002
HCST P = 1.83e-04
MATK P = 0.002
GZMH P = 7.69e-04
GZMB P = 2.46e-04
GNLY P = 0.002
KIR3DL1 P = 2.97e-04

**Supplementary figure 3: Additional analyses of $CD8^+$ T cells in NIC and MS.** (a) Percentage of $CD8^+$ clones shared between CSF and PB among clones in each CSF Leiden cluster. (b) Number of $CD8^+$ T cells in CSF normalized to CSF sample volume across different absolute clonal expansion thresholds. (c) Percentage of $CD8^+$ T cells in CSF among all $CD8^+$ T cells in PB and CSF across different absolute expansion thresholds. (d) Intraclonal percentage of cells in CSF out of all discovered PB and CSF cells belonging to the same clone. Only clones with cells in both PB and CSF are considered. Number of clones displayed is indicated beneath each set of columns. Column plots depict the mean intraclonal percentage ± SD. (e) XY graph depicting absolute cell counts per clone in CSF and PB. CSF- and PB-biased $CD8^+$ T cell clones are highlighted. (f) XY graph depicting the fraction of CSF cells per clone as a function of the total number of CSF/PB cells discovered for the given clone. Black dots indicate statistically significant bias toward CSF, i.e., high CSF fractions, or toward PB, i.e., low CSF fractions, as determined by a binomial test at the given degree of expansion. Blue dashed lines indicate the thresholds used to annotate CSF-biased clones for further analysis: n > 12 cells (vertical line) and a CSF fraction of 0.538 (horizontal line). (g-i) Re-analysis of published data from Kavaka et al. 2024 (GSE276167). (g) Percentage of $CD8^+$ T cells in CSF among all $CD8^+$ T cells in PB and CSF across different absolute expansion thresholds. (h) Columns depicting hyperexpanded $CD8^+$ T cell clones and their relative intraclonal abundance in CSF (blue) or PB (red). IIH, idiopathic intracranial hypertension (equivalent to NIC in our study). n > 12 cells was used to define hyperexpanded clones at a fixed statistical threshold (%CSF > 53.8%). (i) Percentage of CSF-biased $CD8^+$ T cell clones out of all observed clones in PB and CSF at different absolute expansion thresholds. (j) Expression of the CSF-biased score per Leiden cluster. (k) Expression of the PB-biased score per Leiden cluster. (l) Percentage of CSF- and PB-biased genes that were upregulated in CSF (blue), non-enriched (gray), or upregulated in PB. Up- and downregulated genes as well as their *P* values were derived from the analysis shown in Fig. 1k. (m) Average gene expression per donor among select CSF- and PB-biased genes. Unless otherwise stated, *P* values were calculated using a Wilcoxon test and adjusted using the Benjamini-Hochberg procedure. *, **, ***, **** correspond to $P < 0.05$, $P < 0.01$, $P < 0.001$, and $P < 0.0001$, respectively.

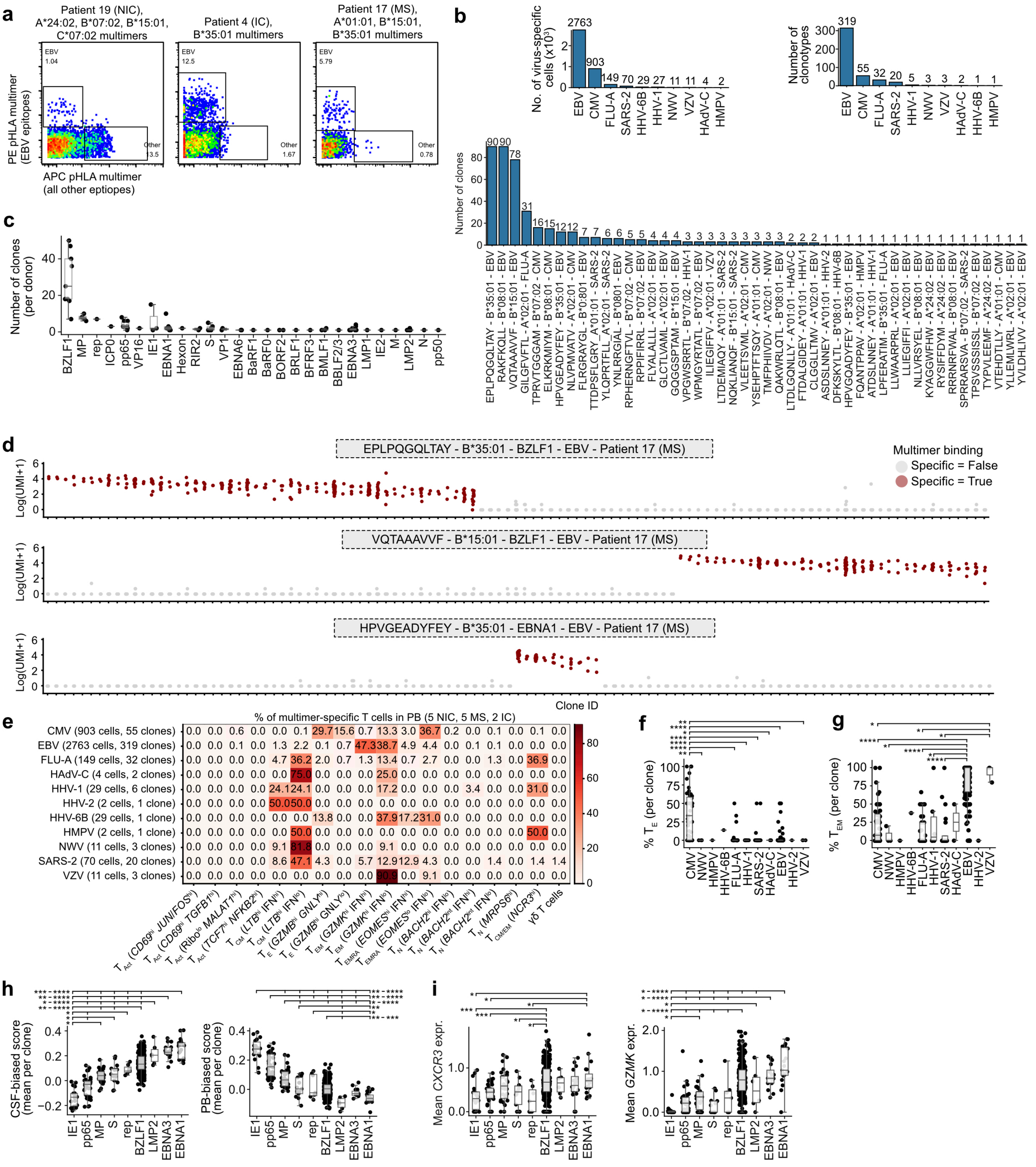
a
Patient 19 (NIC), A*24:02, B*07:02, B*15:01, C*07:02 multimers
Patient 4 (IC), B*35:01 multimers
Patient 17 (MS), A*01:01, B*15:01, B*35:01 multimers
EBV 1.04
EBV 12.5
EBV 5.79
Other 3.5
Other 1.67
Other 0.78
PE pHLA multimer (EBV epitopes)
APC pHLA multimer (all other eptiopes)
b
No. of virus-specific cells (x10³)
Number of clonotypes
Number of clones
c
Number of clones (per donor)
d
EPLPQGQLTAY - B*35:01 - BZLF1 - EBV - Patient 17 (MS)
VQTAAAVVF - B*15:01 - BZLF1 - EBV - Patient 17 (MS)
HPVGEADYFEY - B*35:01 - EBNA1 - EBV - Patient 17 (MS)
Log(UMI+1)
Multimer binding
Specific = False
Specific = True
Clone ID
e
% of multimer-specific T cells in PB (5 NIC, 5 MS, 2 IC)
CMV (903 cells, 55 clones)
EBV (2763 cells, 319 clones)
FLU-A (149 cells, 32 clones)
HAdV-C (4 cells, 2 clones)
HHV-1 (29 cells, 6 clones)
HHV-2 (2 cells, 1 clone)
HHV-6B (29 cells, 1 clone)
HMPV (2 cells, 1 clone)
NWV (11 cells, 3 clones)
SARS-2 (70 cells, 20 clones)
VZV (11 cells, 3 clones)
γδ T cells
f
% T_E (per clone)
g
% T_EM (per clone)
h
CSF-biased score (mean per clone)
PB-biased score (mean per clone)
i
Mean CXCR3 expr.
Mean GZMK expr.

**Supplementary figure 4: Information supporting pHLA multimer-based discovery of antigen-specific $CD8^+$ T cell clones.** (a) Example flow cytometry gates used for sorting $multimer^+$ $CD8^+$ T cells for sc-seq. (b) Bar graphs depicting summary statistics of discovered antigen-specific $CD8^+$ T cells and $CD8^+$ T cell clones by virus specificity and epitope. (c) Boxplots depicting the number of clones per donor discovered for each antigen specificity. (d) Example categorical scatterplot depicting shifted log-transformed UMI counts of individual DNA barcode-labeled pHLA multimers with antigen-specific clones recognizing the highlighted epitope shown in red. (e) Heatmap detailing the phenotypic distribution of cells discovered for each virus specificity. Percentages are calculated horizontally, i.e., as the percentage of each virus specificity. (f) Percentage of $T_E$ $CD8^+$ T cells for each virus specificity. (g) Percentage of $T_{EM}$ $CD8^+$ T cells for each virus specificity. Dots represent clones in (f-g). (h) Mean CSF-biased scores per clone across individual antigens. (i) Mean *CXCR3* and mean *GZMK* gene expression per antigen specificity. Antigens shown in (h-i) each had more than five annotated antigen-specific clones. Virus specificities for each included antigen were: IE1 (CMV), pp65 (CMV), MP (FLU), S (SARS-2), rep (SARS-2), BZLF1 (EBV), LMP2 (EBV), EBNA3 (EBV), EBNA1 (EBV). *P* values were calculated using Dunn's test and adjusted using the Benjamini-Hochberg procedure. *, **, ***, **** correspond to $P < 0.05$, $P < 0.01$, $P < 0.001$, and $P < 0.0001$, respectively.

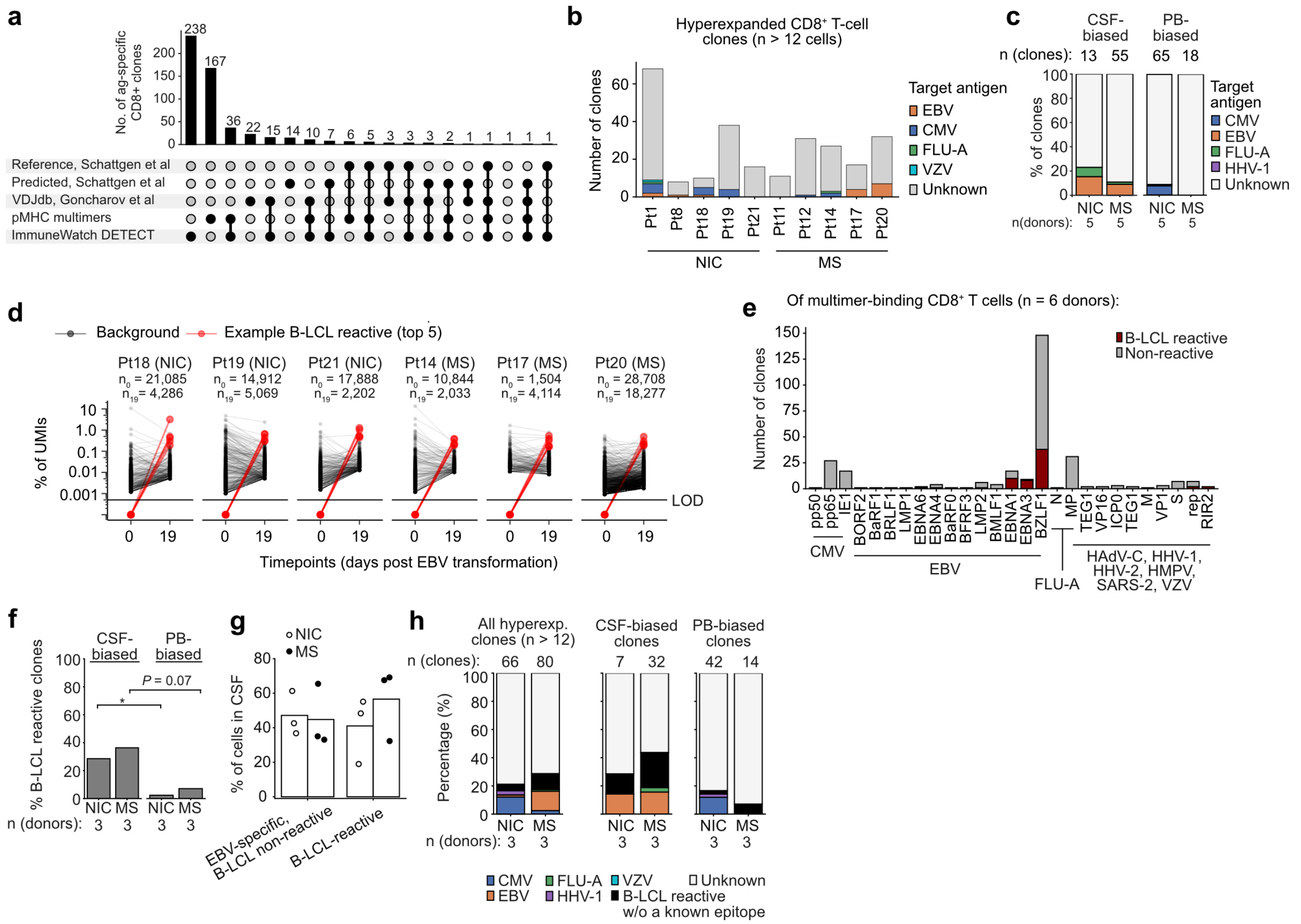


**Supplementary figure 5: Information supporting pHLA multimer-based discovery of antigen-specific CD8⁺ T cell clones.** (a) UpSet plot detailing additional antigen annotation using computational methods, including the ImmuneWatch Detect tool (https://www.immunewatch.com/solutions/imw-detect), cross-referencing VDJdb, as well as using sets of EBV-specific CD8⁺ T cell clones gathered by Schattgen et al. 2025. (b) Antigen specificities discovered among the original hyperexpanded CD8⁺ T cell clones from PB and CSF using pHLA multimers and public databases. (c) Antigen specificities discovered among the original CSF-biased and PB-biased CD8⁺ T cell clones across NIC and MS. (d) Percentage of unique UMIs per TRB CDR3 region detected by bulk TCR-seq of EBV-transformed PBMC cultures on day 0 and day 19, respectively. The top five clones were selected as examples. B-LCL reactive clones were annotated as clones exhibiting a positive fold change in percentage UMI readouts between day 0 and day 19. (e) Correspondence between antigen-specific clones discovered among matched donors (MS, n = 3; NIC, n = 3) and clones detected by bulk TCR-seq. (f) Percentage of B-LCL reactive clones among CSF-biased and PB-biased CD8⁺ T cell clones discovered in the original paired PB and CSF dataset. (g) Percentage of cells in CSF of EBV-specific CD8⁺ T cells split by B-LCL reactivity. (h) Relative proportion of annotated hyperexpanded, CSF-biased and PB-biased CD8⁺ T cell clones in the original paired PB and CSF dataset among donors where bulk B-LCL reactivity was studied. *P* values were calculated using Fisher's exact test. *, **, ***, **** correspond to $P < 0.05$, $P < 0.01$, $P < 0.001$, and $P < 0.0001$, respectively.

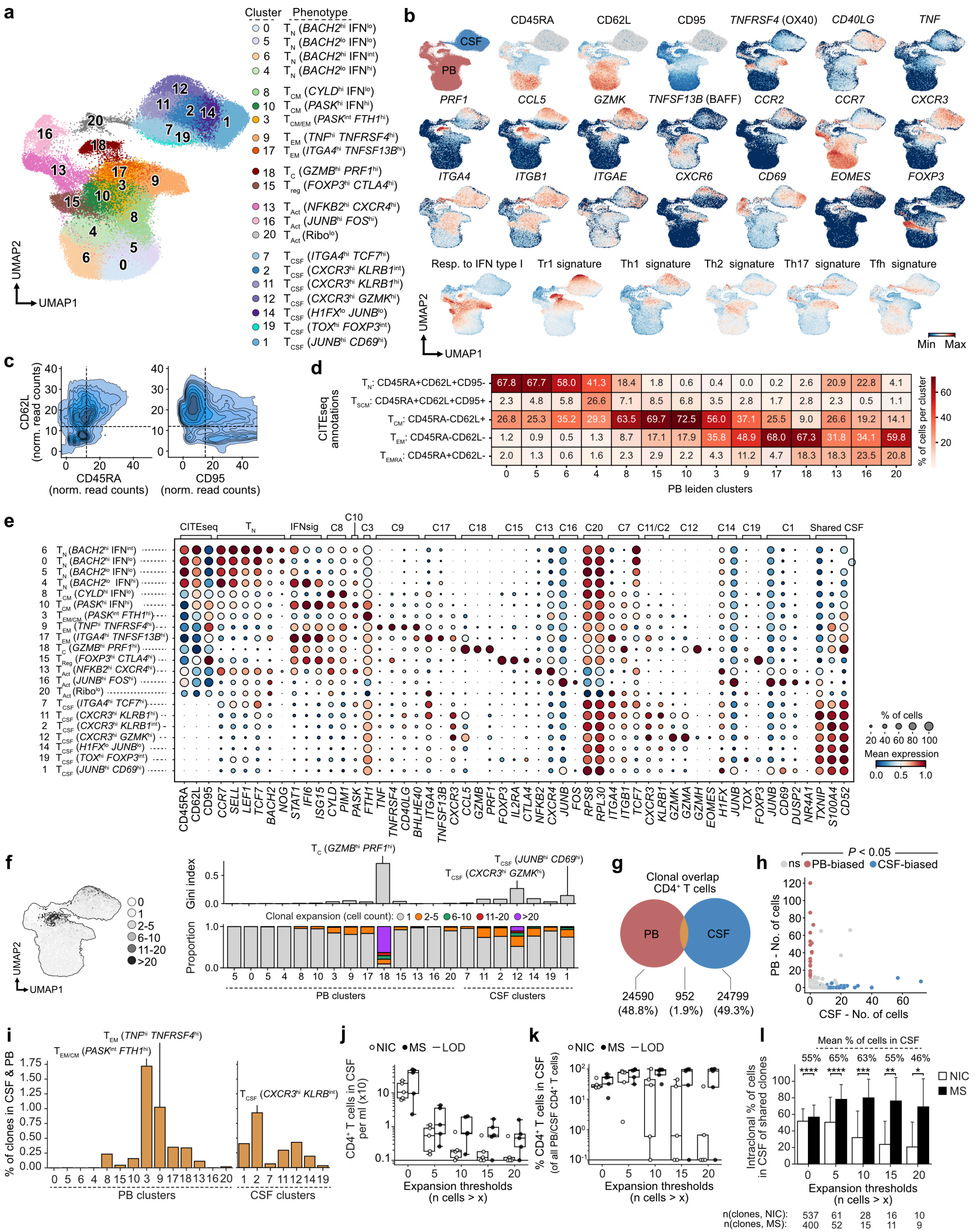

a
Cluster Phenotype
0 TN (BACH2hi IFNlo)
5 TN (BACH2lo IFNlo)
6 TN (BACH2hi IFNint)
4 TN (BACH2lo IFNhi)
8 TCM (CYLDhi IFNlo)
10 TCM (PASKhi IFNhi)
3 TCM/EM (PASKint FTH1hi)
9 TEM (TNFhi TNFRSF4hi)
17 TEM (ITGA4hi TNFSF13Bhi)
18 TC (GZMBhi PRF1hi)
15 Treg (FOXP3hi CTLA4hi)
13 TAct (NFKB2hi CXCR4hi)
16 TAct (JUNBhi FOShi)
20 TAct (Ribolo)
7 TCSF (ITGA4hi TCF7hi)
2 TCSF (CXCR3hi KLRB1int)
11 TCSF (CXCR3hi KLRB1hi)
12 TCSF (CXCR3hi GZMKhi)
14 TCSF (H1FXlo JUNBlo)
19 TCSF (TOXhi FOXP3int)
1 TCSF (JUNBhi CD69hi)
UMAP2
UMAP1
b
CSF
PB
CD45RA
CD62L
CD95
TNFRSF4 (OX40)
CD40LG
TNF
PRF1
CCL5
GZMK
TNFSF13B (BAFF)
CCR2
CCR7
CXCR3
ITGA4
ITGB1
ITGAE
CXCR6
CD69
EOMES
FOXP3
Resp. to IFN type I
Tr1 signature
Th1 signature
Th2 signature
Th17 signature
Tfh signature
Min Max
c
CD62L (norm. read counts)
CD45RA (norm. read counts)
CD95 (norm. read counts)
d
CITEseq annotations
TN: CD45RA+CD62L+CD95-
TSCM: CD45RA+CD62L+CD95+
TCM: CD45RA-CD62L+
TEM: CD45RA-CD62L-
TEMRA: CD45RA+CD62L-
% of cells per cluster
PB leiden clusters
e
CITEseq
TN
IFNsig
C8
C10
C3
C9
C17
C18
C15
C13
C16
C20
C7
C11/C2
C12
C14
C19
C1
Shared CSF
% of cells
20 40 60 80 100
Mean expression
0.0 0.5 1.0
CD45RA CD62L CD95 CCR7 SELL LEF1 TCF7 BACH2 NOG STAT1 IFI6 ISG15 CYLD PIM1 PASK FTH1 TNF TNFRSF4 CD40LG BHLHE40 ITGA4 TNFSF13B CXCR3 CCL5 GZMB PRF1 FOXP3 IL2RA CTLA4 NFKB2 CXCR4 JUNB FOS RPS8 RPL30 ITGA4 ITGB1 TCF7 CXCR3 KLRB1 GZMK GZMA GZMH EOMES H1FX JUNB TOX FOXP3 JUNB CD69 DUSP2 NR4A1 TXNIP S100A4 CD52
f
0
1
2-5
6-10
11-20
>20
UMAP2
UMAP1
Gini index
TC (GZMBhi PRF1hi)
TCSF (JUNBhi CD69hi)
TCSF (CXCR3hi GZMKhi)
Clonal expansion (cell count): 1 2-5 6-10 11-20 >20
Proportion
PB clusters
CSF clusters
g
Clonal overlap CD4+ T cells
PB
CSF
24590 (48.8%)
952 (1.9%)
24799 (49.3%)
h
P < 0.05
ns PB-biased CSF-biased
PB - No. of cells
CSF - No. of cells
i
TEM (TNFhi TNFRSF4hi)
TEM/CM (PASKint FTH1hi)
TCSF (CXCR3hi KLRB1int)
% of clones in CSF & PB
PB clusters
CSF clusters
j
NIC MS LOD
CD4+ T cells in CSF per ml (x10)
Expansion thresholds (n cells > x)
k
NIC MS LOD
% CD4+ T cells in CSF (of all PB/CSF CD4+ T cells)
Expansion thresholds (n cells > x)
l
Mean % of cells in CSF
55% 65% 63% 55% 46%
**** **** *** ** *
NIC
MS
Intraclonal % of cells in CSF of shared clones
Expansion thresholds (n cells > x)
n(clones, NIC): 537 61 28 16 10
n(clones, MS): 400 52 15 11 9

**Supplementary figure 6: Phenotypic and clonal diversity of PB and CSF CD4$^+$ T cells.** (a) UMAP of 69,173 and 36,032 CD4$^+$ T cells from PB and CSF, respectively, with Leiden clusters and compartment annotations. Five NIC, five MS and two IC donors are shown. For brevity, PB origin is not explicitly indicated, whereas CSF-derived populations are labeled $T_{CSF}$. (b) Transcriptomic features related to CD4$^+$ T cell differentiation, function and phenotype. (c) Normalized CITE-seq count values displayed as a density plot. Threshold values to define virtual gates for $T_N$, $T_{SCM}$, $T_{CM}$, $T_{EM}$, and $T_{EMRA}$ are indicated as dashed lines. (d) Quantification and annotation of Leiden clusters according to virtual gates shown in (c). (e) Gene expression summary of individually annotated Leiden clusters using general marker genes as well as selected marker genes for each cluster. (f) UMAP of absolute clonal expansion together with side-by-side quantifications of clonal unevenness (Gini index) and proportion of cells in different absolute expansion bins. (g) Venn diagram of CD4$^+$ T cell clones shared between PB and CSF. (h) XY graph depicting absolute cell counts per clone in CSF and PB. CSF- and PB-biased CD4$^+$ T cell clones are highlighted. (i) Percentage of CD4$^+$ clones shared between CSF and PB among clones in each Leiden cluster. (j) Number of CD4$^+$ T cells in CSF normalized to CSF sample volume across different absolute clonal expansion thresholds. (k) Percentage of CD4$^+$ T cells in CSF among all CD4$^+$ T cells in PB and CSF across different absolute expansion thresholds. (l) Intraclonal percentage of cells in CSF out of all discovered PB and CSF cells belonging to the same clone. Only clones with cells in both PB and CSF are considered. Number of clones displayed is indicated beneath each set of columns. Column plots depict the mean intraclonal percentage ± SD. Statistical thresholds to annotate CSF-biased CD4$^+$ T cells in (h) were computed by permutation of a binomial distribution using the relative proportions of total CSF and PB cells as estimates of the expected frequencies. $P$ values in (l) were calculated using a Wilcoxon test and adjusted using the Benjamini-Hochberg procedure. *, **, ***, **** correspond to $P < 0.05$, $P < 0.01$, $P < 0.001$, and $P < 0.0001$, respectively.

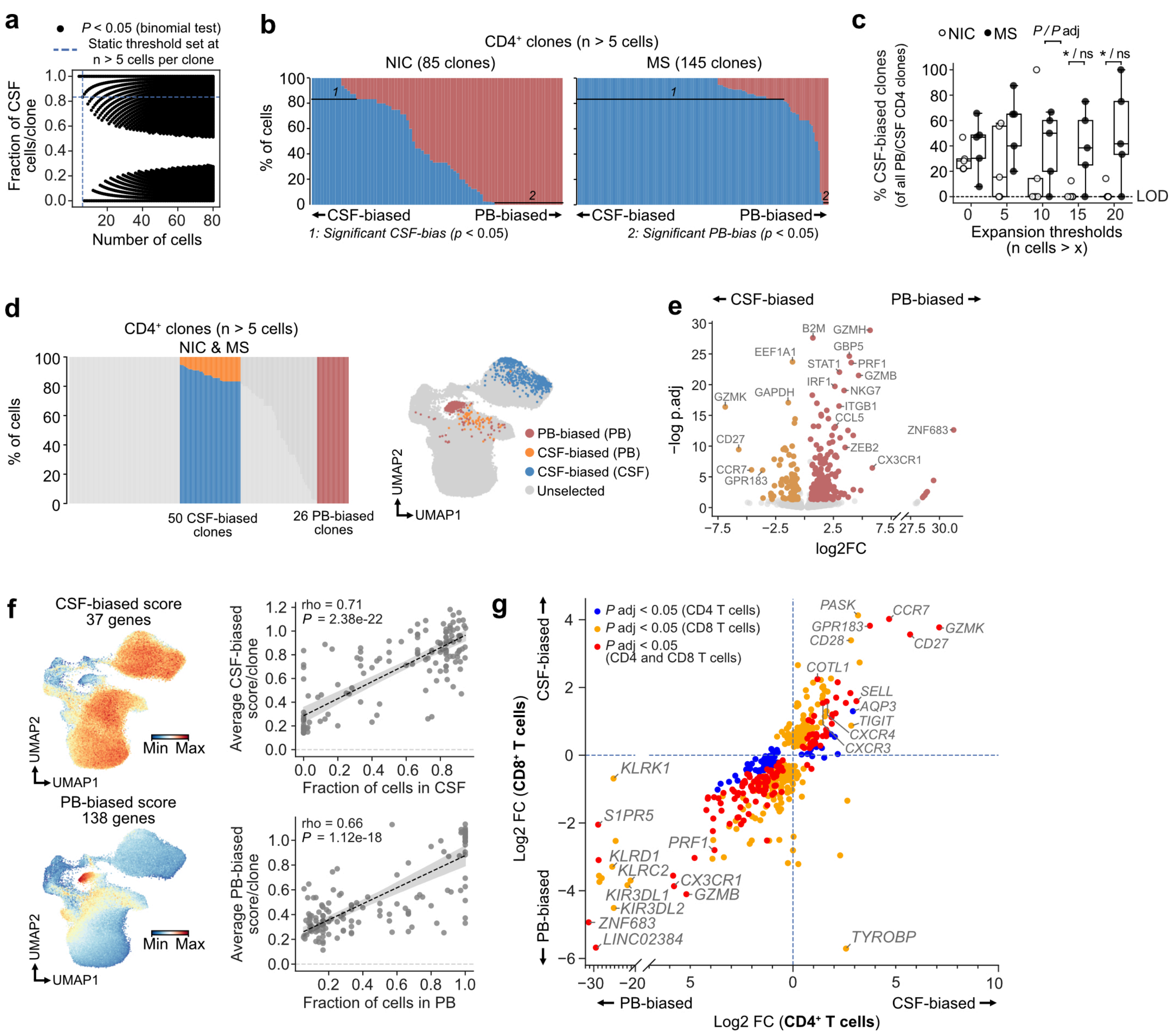

**Supplementary figure 7: Expanded CSF-biased CD4⁺ T cell clones in PB are more frequent in MS patients and share transcriptomic features with CSF-biased CD8⁺ T cells, including *GZMK* expression.** (a) XY graph depicting the fraction of CSF cells per clone as a function of the total number of CSF and PB cells discovered for the given clone. Black dots indicate statistically significant bias toward either CSF, i.e., high CSF fractions, or PB, i.e., low CSF fractions. Blue dashed lines indicate thresholds used to annotate CSF-biased clones at n > 5 cells. (b) Annotation of CSF- and PB-biased CD4⁺ T cells. Columns depict expanded CD4⁺ T cell clones and their relative intraclonal abundance in CSF (blue) or PB (red). A threshold of n > 5 cells was used to define expanded clones at a fixed statistical threshold (%CSF > 83.3%) according to the XY graph in (a). (c) Percentage of CSF-biased CD4⁺ T cell clones out of all CD4⁺ T cell clones discovered across different expansion thresholds. (d) Column plots and UMAP highlighting highly CSF- and PB-biased CD4⁺ T cells. Orange indicates CSF-biased CD4⁺ T cells in PB. (e) Differential gene expression analysis of CSF- and PB-biased CD4⁺ T cells from PB. (f) Gene scores related to the differentially expressed genes found in (e), together with Spearman correlations depicting the positive relationships between CSF- and PB-biased gene scores and the fraction of cells in CSF and PB, respectively, among expanded CD4⁺ T cell clones. (g) XY graph comparing the log2 fold change values of differentially expressed genes among CSF- and PB-biased

CD4$^+$ T cells with the corresponding log2 fold change values of differentially expressed genes among CSF- and PB-biased CD8$^+$ T cells. Statistical thresholds to annotate CSF-biased CD4$^+$ T cells in (a-b) were computed by permutation of a binomial distribution using the relative proportions of total CSF and PB cells as estimates of the expected frequencies. $P$ values were calculated using a Wilcoxon test and adjusted using the Benjamini-Hochberg procedure. *, **, ***, **** correspond to $P < 0.05$, $P < 0.01$, $P < 0.001$, and $P < 0.0001$, respectively.

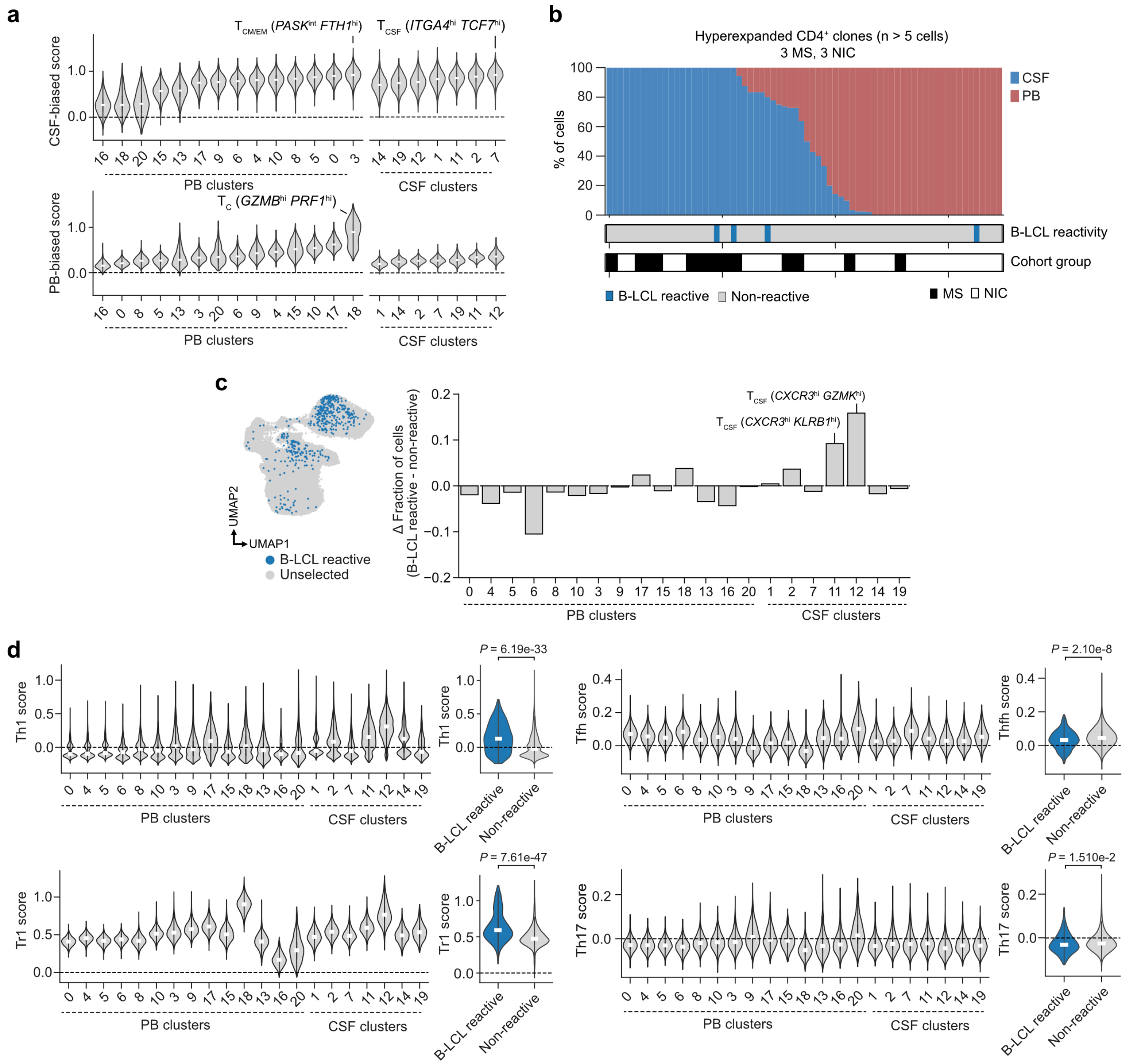


**Supplementary figure 8: B-LCL reactive CD4+ T cells are enriched among *CXCR3*hi clusters in the CSF and resemble Th1 and Tr1-like cells.** (a) CSF- and PB-biased scores calculated per cell and summarized by cluster. (b) B-LCL reactivity among CD4+ expanded T cells visualized as bar graphs in six donors (3 MS, 3 NIC) used in the LCL-reactivity assay. Additional annotations indicate B-LCL reactivity and donor groups. (c) B-LCL reactive cells, highlighted on a UMAP and summarized as column plots of residual B-LCL reactivity per cluster after subtraction of the non-reactive CD4+ T cell fraction. (d) Th1, Tfh, Tr1 and Th17 scores per cell summarized across compartments and phenotypic clusters, together with summarized expression among B-LCL reactive and non-reactive CD4+ T cells. Th1, Tfh and Th17 scores were derived from Ostkamp et al. 2022, and the Tr1 score from Bonnal et al. 2021. *P* values were calculated using a Welch's *t* test.

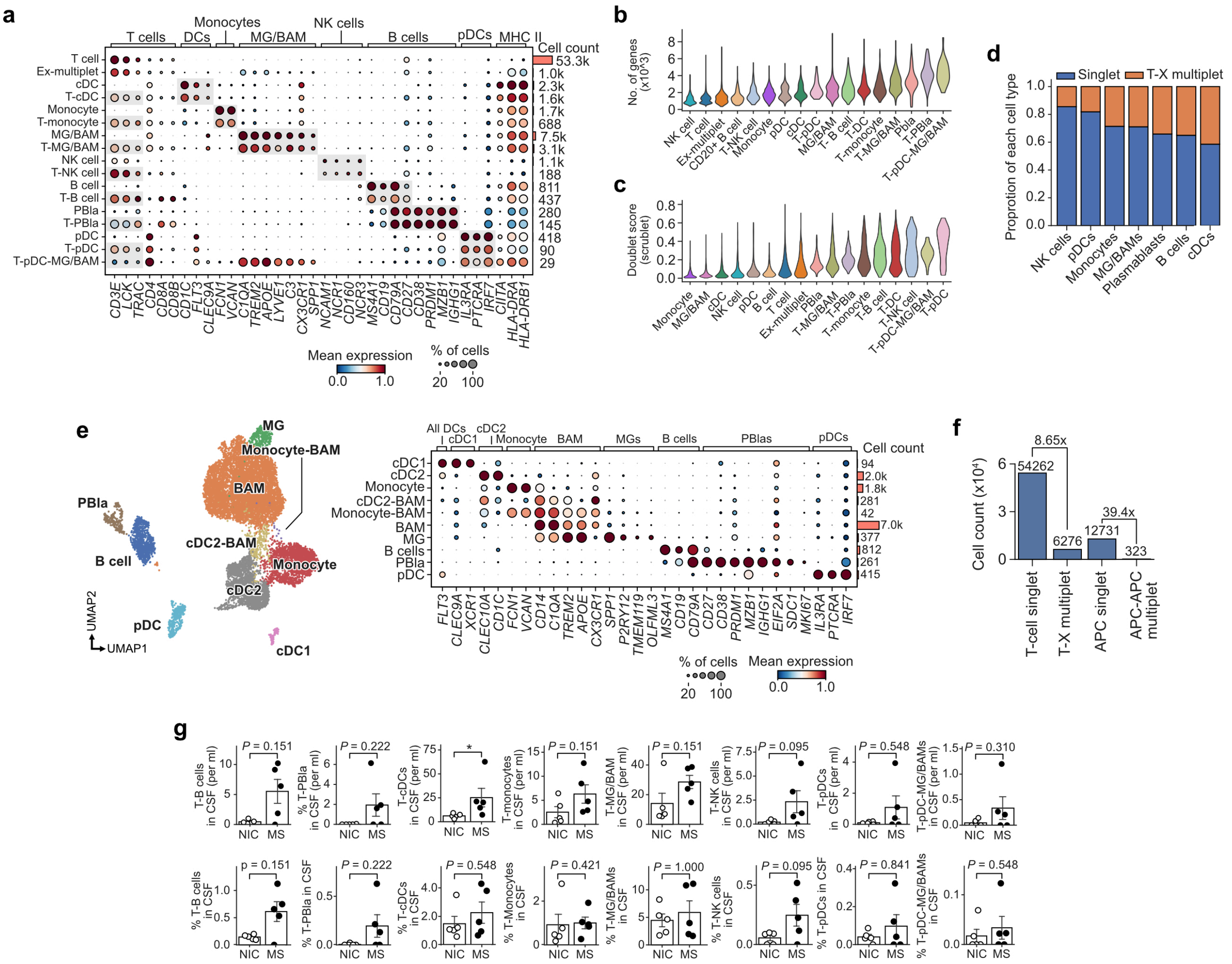

**Supplementary figure 9: Supporting information for the identification and characterization of T–X multiplets in CSF.** (a) Gene expression summary of reclustered immune cell subsets from CSF. Marker gene sets used to annotate corresponding singlet and T–X multiplet clusters are highlighted in gray. (b) Number of detected genes per cell in each identified cluster. (c) Doublet score calculated by Scrublet per identified cluster. (d) Proportion of identified cells (x-axis) found among singlets (blue) or T–X multiplets (orange). (e) Reclustered and reannotated APC subsets, including identification of APC–APC multiplets. (f) Total cell counts of singlet and multiplet subsets. Ratios between specific singlets and multiplets are highlighted. (g) Cell counts per mL CSF (top row) and frequency in CSF (bottom row) for all T–X multiplets. *P* values were calculated using a Wilcoxon test. *, *P* < 0.05.

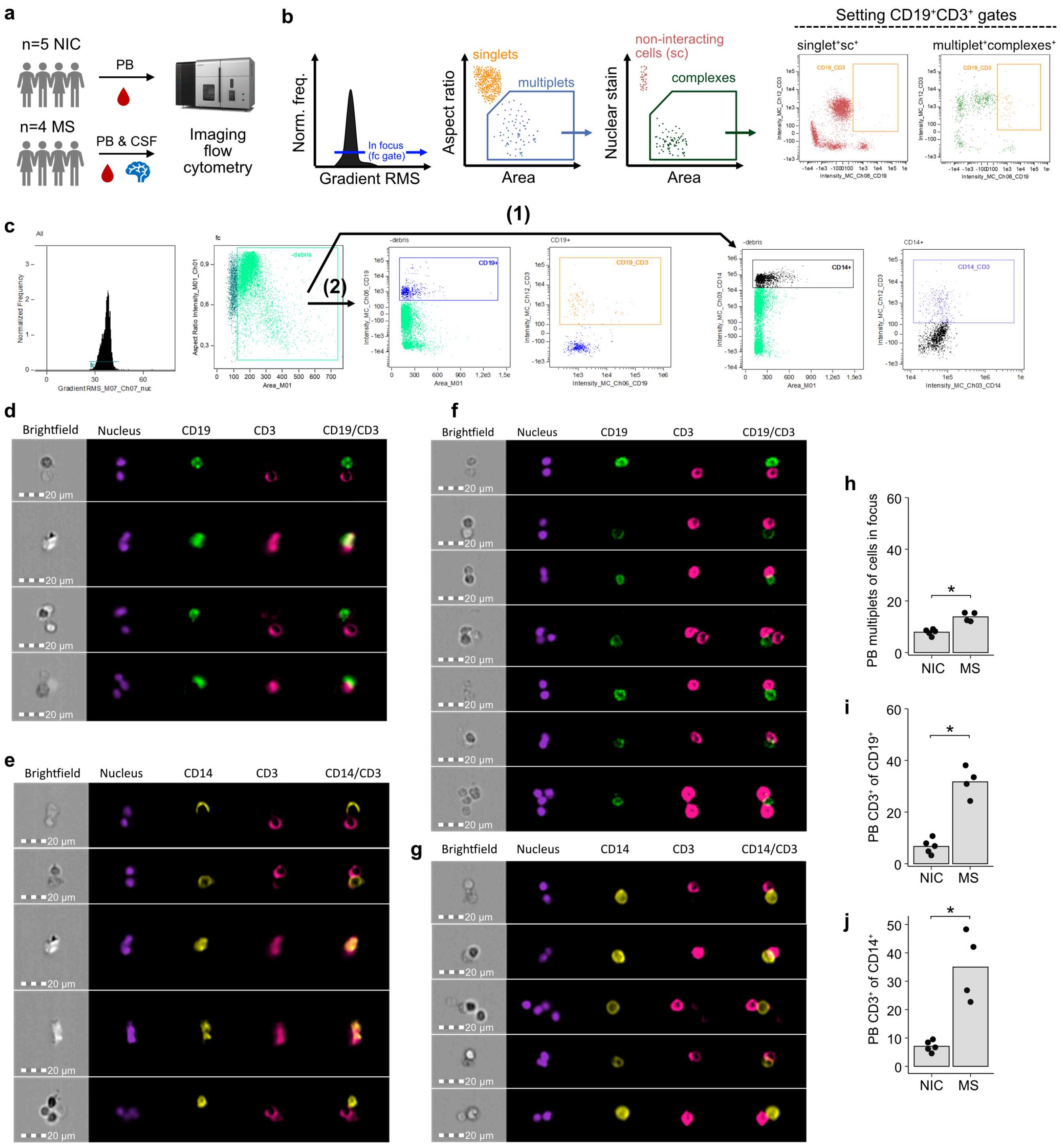

a
n=5 NIC
n=4 MS
PB
PB & CSF
Imaging flow cytometry
b
Norm. freq.
In focus (fc gate)
Gradient RMS
Aspect ratio
singlets
multiplets
Area
Nuclear stain
non-interacting cells (sc)
complexes
Area
Setting CD19+CD3+ gates
singlet+sc+
multiplet+complexes+
c
(1)
(2)
d
Brightfield
Nucleus
CD19
CD3
CD19/CD3
20 µm
e
Brightfield
Nucleus
CD14
CD3
CD14/CD3
f
Brightfield
Nucleus
CD19
CD3
CD19/CD3
g
Brightfield
Nucleus
CD14
CD3
CD14/CD3
h
PB multiplets of cells in focus
NIC
MS
i
PB CD3+ of CD19+
NIC
MS
j
PB CD3+ of CD14+
NIC
MS

**Supplementary figure 10: Identification of $CD3^+CD19^+$ and $CD3^+CD14^+$ multiplets using imaging flow cytometry.** (a) Cohort and sample type overview. CSF was available only from MS patients. (b) Gating strategy for defining multiplet gates for subsequent analyses relied on gating interacting cells in focus ($multiplet^+$, $complexes^+$), with $CD3^+CD19^+$ and $CD3^+CD14^+$ gates defined using negative intrasample controls gated on singlets in focus ($singlet^+$, $sc^+$). $CD3^+CD19^+$ gating strategy is here shown as an example. (c) Gating strategy for quantification of (1) $CD3^+CD14^+$ and (2) $CD3^+CD19^+$ multiplets. (d-g) Examples of T cells interacting with antigen-presenting cells in MS patients. (d) T cells interacting with $CD19^+$ B cells in CSF. (e) T cells interacting with $CD14^+$ monocytes/macrophages in CSF. (f) T cells interacting with $CD19^+$ B cells in PB. (g) T cells interacting with $CD14^+$ monocytes/macrophages in PB. (h) Frequency of multiplets in PB among cells in focus. (i) Frequency of $CD3^+$ multiplets out of $CD19^+$ cells. (j) Frequency of $CD3^+$ multiplets out of $CD14^+$ cells. *P* values were calculated using a Wilcoxon test. *, $P < 0.05$.

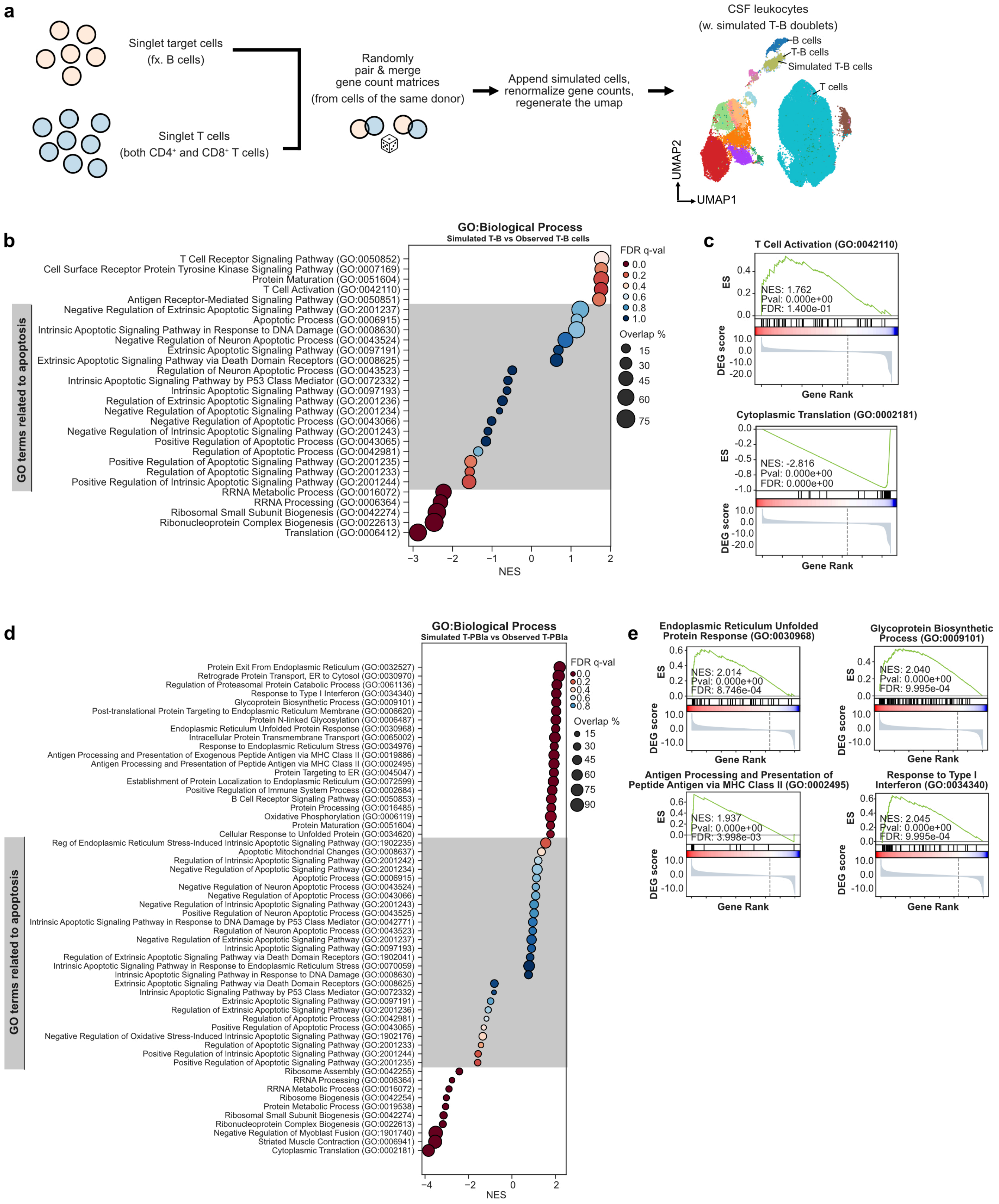

a
Singlet target cells
(fx. B cells)
Singlet T cells
(both CD4+ and CD8+ T cells)
Randomly
pair & merge
gene count matrices
(from cells of the same donor)
Append simulated cells,
renormalize gene counts,
regenerate the umap
CSF leukocytes
(w. simulated T-B doublets)
B cells
T-B cells
Simulated T-B cells
T cells
UMAP2
UMAP1
b
GO:Biological Process
Simulated T-B vs Observed T-B cells
GO terms related to apoptosis
T Cell Receptor Signaling Pathway (GO:0050852)
Cell Surface Receptor Protein Tyrosine Kinase Signaling Pathway (GO:0007169)
Protein Maturation (GO:0051604)
T Cell Activation (GO:0042110)
Antigen Receptor-Mediated Signaling Pathway (GO:0050851)
Negative Regulation of Extrinsic Apoptotic Signaling Pathway (GO:2001237)
Apoptotic Process (GO:0006915)
Intrinsic Apoptotic Signaling Pathway in Response to DNA Damage (GO:0008630)
Negative Regulation of Neuron Apoptotic Process (GO:0043524)
Extrinsic Apoptotic Signaling Pathway (GO:0097191)
Extrinsic Apoptotic Signaling Pathway via Death Domain Receptors (GO:0008625)
Regulation of Neuron Apoptotic Process (GO:0043523)
Intrinsic Apoptotic Signaling Pathway by P53 Class Mediator (GO:0072332)
Intrinsic Apoptotic Signaling Pathway (GO:0097193)
Regulation of Extrinsic Apoptotic Signaling Pathway (GO:2001236)
Negative Regulation of Apoptotic Signaling Pathway (GO:2001234)
Negative Regulation of Apoptotic Process (GO:0043066)
Negative Regulation of Intrinsic Apoptotic Signaling Pathway (GO:2001243)
Positive Regulation of Apoptotic Process (GO:0043065)
Regulation of Apoptotic Process (GO:0042981)
Positive Regulation of Apoptotic Signaling Pathway (GO:2001235)
Regulation of Apoptotic Signaling Pathway (GO:2001233)
Positive Regulation of Intrinsic Apoptotic Signaling Pathway (GO:2001244)
RRNA Metabolic Process (GO:0016072)
RRNA Processing (GO:0006364)
Ribosomal Small Subunit Biogenesis (GO:0042274)
Ribonucleoprotein Complex Biogenesis (GO:0022613)
Translation (GO:0006412)
−3
−2
−1
0
1
2
NES
FDR q-val
0.0
0.2
0.4
0.6
0.8
1.0
Overlap %
15
30
45
60
75
c
T Cell Activation (GO:0042110)
ES
0.4
0.2
0.0
NES: 1.762
Pval: 0.000e+00
FDR: 1.400e-01
DEG score
10.0
0.0
-10.0
-20.0
Gene Rank
Cytoplasmic Translation (GO:0002181)
0.0
-0.2
-0.5
-0.8
-1.0
NES: -2.816
Pval: 0.000e+00
FDR: 0.000e+00
Gene Rank
d
GO:Biological Process
Simulated T-PBla vs Observed T-PBla
GO terms related to apoptosis
Protein Exit From Endoplasmic Reticulum (GO:0032527)
Retrograde Protein Transport, ER to Cytosol (GO:0030970)
Regulation of Proteasomal Protein Catabolic Process (GO:0061136)
Response to Type I Interferon (GO:0034340)
Glycoprotein Biosynthetic Process (GO:0009101)
Post-translational Protein Targeting to Endoplasmic Reticulum Membrane (GO:0006620)
Protein N-linked Glycosylation (GO:0006487)
Endoplasmic Reticulum Unfolded Protein Response (GO:0030968)
Intracellular Protein Transmembrane Transport (GO:0065002)
Response to Endoplasmic Reticulum Stress (GO:0034976)
Antigen Processing and Presentation of Exogenous Peptide Antigen via MHC Class II (GO:0019886)
Antigen Processing and Presentation of Peptide Antigen via MHC Class II (GO:0002495)
Protein Targeting to ER (GO:0045047)
Establishment of Protein Localization to Endoplasmic Reticulum (GO:0072599)
Positive Regulation of Immune System Process (GO:0002684)
B Cell Receptor Signaling Pathway (GO:0050853)
Protein Processing (GO:0016485)
Oxidative Phosphorylation (GO:0006119)
Protein Maturation (GO:0051604)
Cellular Response to Unfolded Protein (GO:0034620)
Reg of Endoplasmic Reticulum Stress-Induced Intrinsic Apoptotic Signaling Pathway (GO:1902235)
Apoptotic Mitochondrial Changes (GO:0008637)
Regulation of Intrinsic Apoptotic Signaling Pathway (GO:2001242)
Negative Regulation of Apoptotic Signaling Pathway (GO:2001234)
Apoptotic Process (GO:0006915)
Negative Regulation of Neuron Apoptotic Process (GO:0043524)
Negative Regulation of Apoptotic Process (GO:0043066)
Negative Regulation of Intrinsic Apoptotic Signaling Pathway (GO:2001243)
Positive Regulation of Neuron Apoptotic Process (GO:0043525)
Intrinsic Apoptotic Signaling Pathway in Response to DNA Damage by P53 Class Mediator (GO:0042771)
Regulation of Neuron Apoptotic Process (GO:0043523)
Negative Regulation of Extrinsic Apoptotic Signaling Pathway (GO:2001237)
Intrinsic Apoptotic Signaling Pathway (GO:0097193)
Regulation of Extrinsic Apoptotic Signaling Pathway via Death Domain Receptors (GO:1902041)
Intrinsic Apoptotic Signaling Pathway in Response to Endoplasmic Reticulum Stress (GO:0070059)
Intrinsic Apoptotic Signaling Pathway in Response to DNA Damage (GO:0008630)
Extrinsic Apoptotic Signaling Pathway via Death Domain Receptors (GO:0008625)
Intrinsic Apoptotic Signaling Pathway by P53 Class Mediator (GO:0072332)
Extrinsic Apoptotic Signaling Pathway (GO:0097191)
Regulation of Extrinsic Apoptotic Signaling Pathway (GO:2001236)
Regulation of Apoptotic Process (GO:0042981)
Positive Regulation of Apoptotic Process (GO:0043065)
Negative Regulation of Oxidative Stress-Induced Intrinsic Apoptotic Signaling Pathway (GO:1902176)
Regulation of Apoptotic Signaling Pathway (GO:2001233)
Positive Regulation of Intrinsic Apoptotic Signaling Pathway (GO:2001244)
Positive Regulation of Apoptotic Signaling Pathway (GO:2001235)
Ribosome Assembly (GO:0042255)
RRNA Processing (GO:0006364)
RRNA Metabolic Process (GO:0016072)
Ribosome Biogenesis (GO:0042254)
Protein Metabolic Process (GO:0019538)
Ribosomal Small Subunit Biogenesis (GO:0042274)
Ribonucleoprotein Complex Biogenesis (GO:0022613)
Negative Regulation of Myoblast Fusion (GO:1901740)
Striated Muscle Contraction (GO:0006941)
Cytoplasmic Translation (GO:0002181)
−4
−2
0
2
NES
FDR q-val
0.0
0.2
0.4
0.6
0.8
Overlap %
15
30
45
60
75
90
e
Endoplasmic Reticulum Unfolded
Protein Response (GO:0030968)
ES
NES: 2.014
Pval: 0.000e+00
FDR: 8.746e-04
DEG score
Gene Rank
Glycoprotein Biosynthetic
Process (GO:0009101)
NES: 2.040
Pval: 0.000e+00
FDR: 9.995e-04
Gene Rank
Antigen Processing and Presentation of
Peptide Antigen via MHC Class II (GO:0002495)
NES: 1.937
Pval: 0.000e+00
FDR: 3.998e-03
Gene Rank
Response to Type I
Interferon (GO:0034340)
NES: 2.045
Pval: 0.000e+00
FDR: 9.995e-04
Gene Rank

**Supplementary figure 11: Differential gene expression and gene set enrichment analyses for characterization of pathways and phenotypes associated with T–B and T–PBIa multiplets.** (a) Detailed scheme for computational mixing of singlet T cells and APCs, with B cells used as an example. T and B cell singlets were randomly sampled from each donor according to the smaller of the two parent populations. (b) Gene set enrichment analyses using differentially expressed genes in observed T–B multiplets compared with simulated T–B multiplets as input. GO Biological Process gene sets were used as the reference database to identify enriched pathways. Pathways/gene sets related to apoptotic processes were manually curated. (c) Example gene set enrichment plots from the analysis done in (b). (d) Gene set enrichment analyses as in (b), but using genes differentially expressed in observed T–PBIa multiplets compared to simulated T–PBIa multiplets as input. (e) Example gene set enrichment plots from the analysis done in (d).

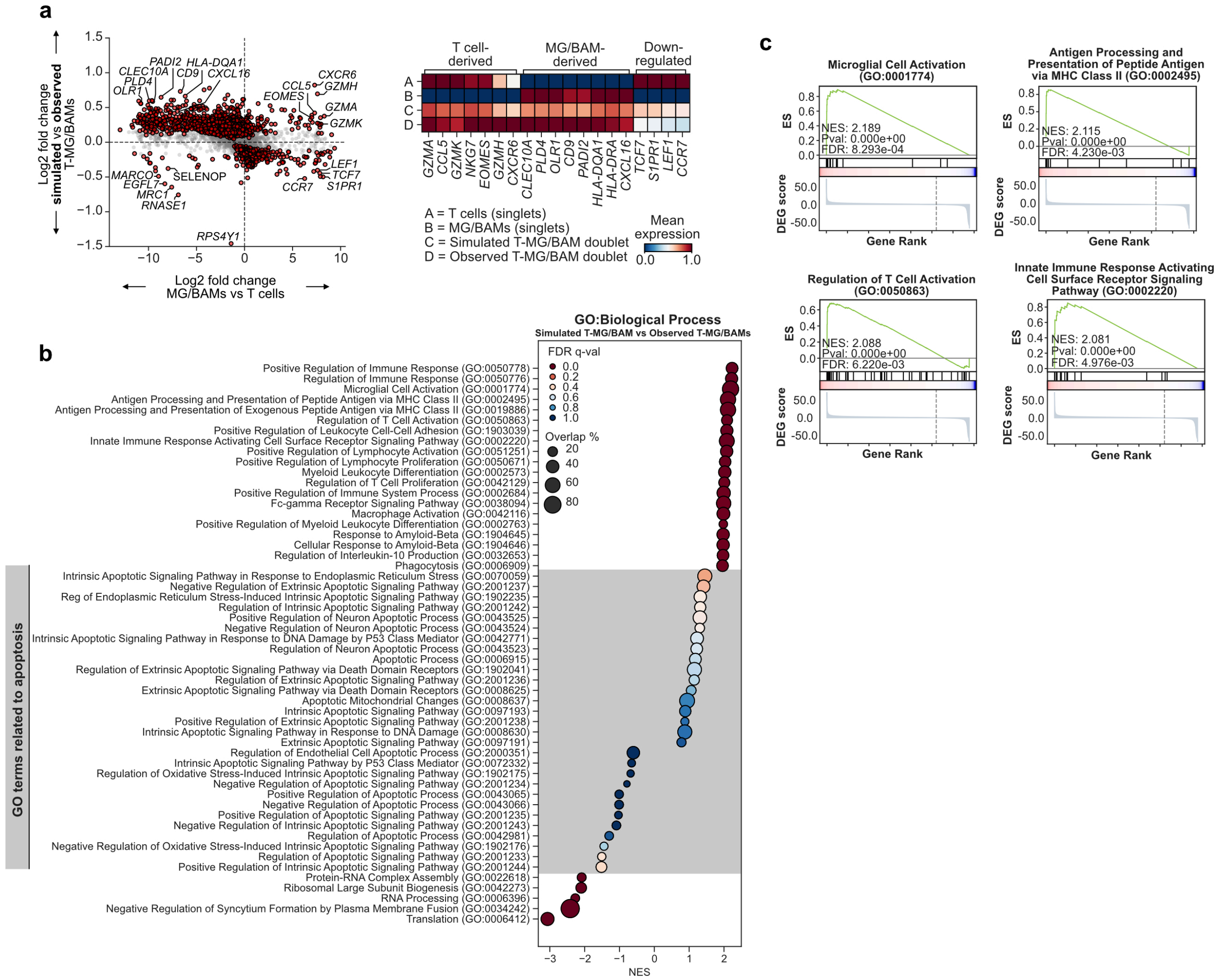


**Supplementary figure 12: Supplementary analysis of T–MG/BAM multiplets. (a) Comparison of differentially expressed genes.** T cell- and MG/BAM-specific expression (x-axis) versus differential gene expression in observed multiplets compared with simulated doublets generated from singlet parent populations (y-axis). A matrix plot accompanies the scatter plot as a summary of noteworthy differentially expressed genes. (b) Gene set enrichment analysis using differentially expressed genes in observed T–MG/BAM multiplets compared with simulated T–MG/BAM multiplets as input. GO Biological Process was used as the target database. (c) Example gene set enrichment plots from the analysis done in (b).

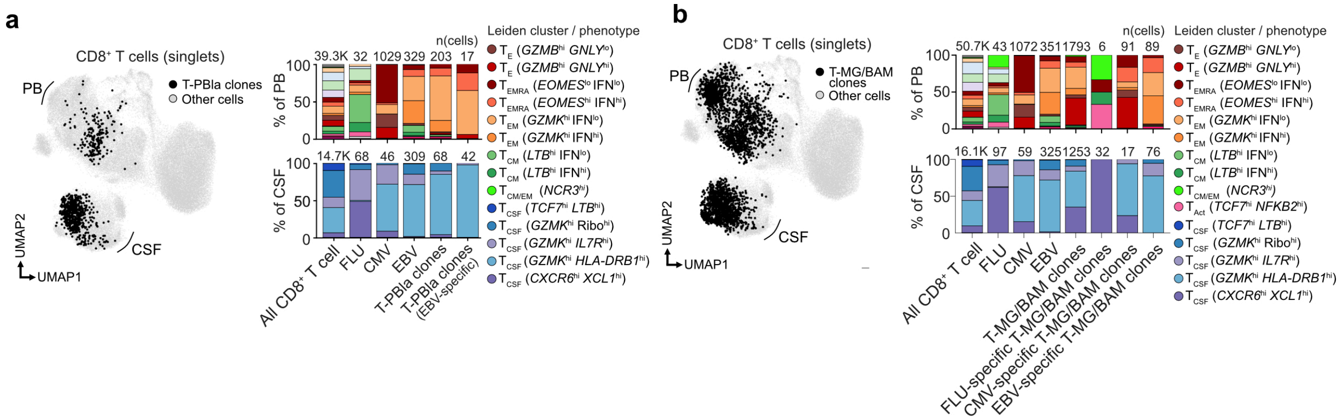


**Supplementary figure 13: Phenotypes associated with antigen-specific interactions in T–PBIa and T–MG/BAM multiplets.** (a) UMAP and phenotypic distribution highlighting singlet cells belonging to clones interacting with PBIa in CSF. Only donors with detectable PBIa or T–PBIa multiplets in CSF are represented (four MS patients and four NIC). (b) UMAP and phenotypic distribution highlighting singlet cells belonging to clones interacting with MG/BAMs in CSF. All MS and NIC donors are represented.

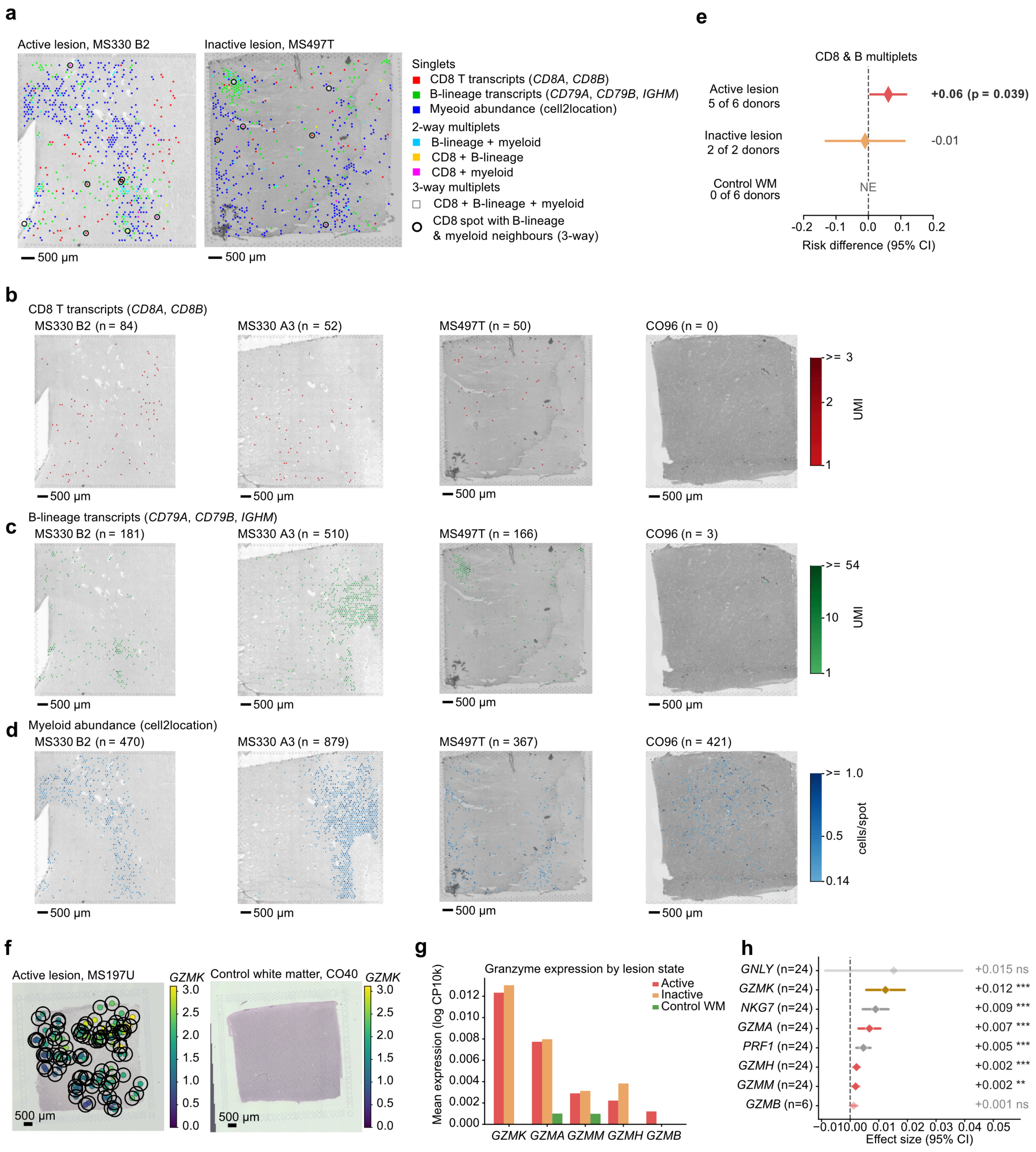
a
Active lesion, MS330 B2
Inactive lesion, MS497T
Singlets
CD8 T transcripts (CD8A, CD8B)
B-lineage transcripts (CD79A, CD79B, IGHM)
Myeloid abundance (cell2location)
2-way multiplets
B-lineage + myeloid
CD8 + B-lineage
CD8 + myeloid
3-way multiplets
CD8 + B-lineage + myeloid
CD8 spot with B-lineage & myeloid neighbours (3-way)
500 μm
e
CD8 & B multiplets
Active lesion 5 of 6 donors
+0.06 (p = 0.039)
Inactive lesion 2 of 2 donors
-0.01
Control WM 0 of 6 donors
NE
Risk difference (95% CI)
b
CD8 T transcripts (CD8A, CD8B)
MS330 B2 (n = 84)
MS330 A3 (n = 52)
MS497T (n = 50)
CO96 (n = 0)
UMI
c
B-lineage transcripts (CD79A, CD79B, IGHM)
MS330 B2 (n = 181)
MS330 A3 (n = 510)
MS497T (n = 166)
CO96 (n = 3)
d
Myeloid abundance (cell2location)
MS330 B2 (n = 470)
MS330 A3 (n = 879)
MS497T (n = 367)
CO96 (n = 421)
cells/spot
f
Active lesion, MS197U
Control white matter, CO40
GZMK
g
Granzyme expression by lesion state
Active
Inactive
Control WM
Mean expression (log CP10k)
GZMK GZMA GZMM GZMH GZMB
h
GNLY (n=24) +0.015 ns
GZMK (n=24) +0.012 ***
NKG7 (n=24) +0.009 ***
GZMA (n=24) +0.007 ***
PRF1 (n=24) +0.005 ***
GZMH (n=24) +0.002 ***
GZMM (n=24) +0.002 **
GZMB (n=6) +0.001 ns
Effect size (95% CI)

**Supplementary figure 14: Supporting data and enrichment of *GZMK*-expressing spots in active lesions compared to control sections.** (a) Additional representative Visium sections from Lerma-Martin et al. 2024 of active and inactive MS lesions from postmortem MS patients. Spots per section: CD8 84 and 50; B lineage 181 and 166; myeloid 470 and 367; ringed 8 and 6. (b-d) Spots are disaggregated by cell type annotation. (b) CD8 T cells, (c) B cells, (d) myeloid cells. (e) Risk difference, the fraction of spots with CD8 transcripts and B lineage transcripts in the same or an adjacent spot by lesion state minus the same fraction among other lymphoid spots. Whiskers indicate the 95% confidence interval. A section is not estimable (NE) with fewer than 20 CD8-B spots or background spots. One active-lesion donor and six control-WM donors were NE. The observed effect among CD8–B neighborhoods is carried by Lerma-Martin et al. 2024 sections (+0.075, $P = 0.003$) and absent in the single active donor from Schroeder et al. 2025 sections (+0.005, $P = 0.84$). (f) *GZMK* expression (CP10k, log1p) per Visium spot in an active lesion and in control white matter. Circles mark *GZMK*-positive spots. (g) Mean normalized gene expression of granzymes per lesion state across 24 sections. (h) Effect size per gene, i.e., differences in mean gene expression between active and control lesions modeled using ordinary least squares of spot-level gene expression across 24 Visium sections. n, sections in which the gene is measured; 95% CI, 95 percent confidence intervals; GZMK gold, other granzymes salmon, cytotoxic reference genes gray. *GZMB* is represented as a mean instead of a contrast as it was only measured in sections from a single donor. *$P < 0.05$, **$P < 0.01$, ***$P < 0.001$. *P* values were calculated using a donor-level DerSimonian–Laird random-effects analysis with both normal-approximation and Hartung–Knapp inference in (e), and a cluster-robust Wald test in (h). Visium sections originate from Lerma-Martin et al. 2024 and Schroeder et al. 2025.

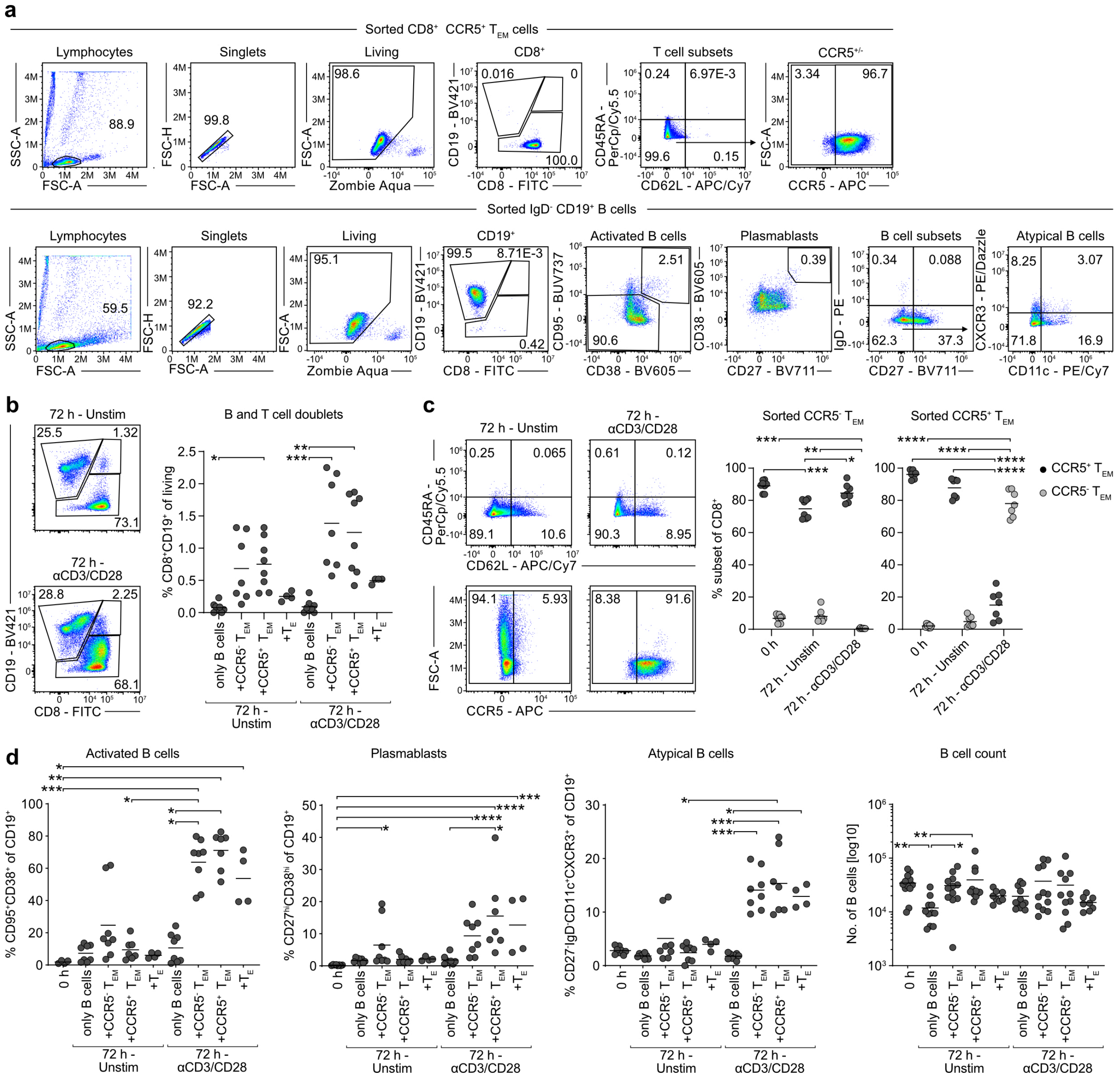

**Supplementary figure 15: GZMK$^+$ T$_{EM}$ cells influence the activation and differentiation of B cells.** (a) Representative flow cytometry data from one donor showing the gating strategy for sorted CD8$^+$ CCR5$^+$ T$_{EM}$ cells and CD19$^+$ IgD$^-$ B cells before co-culture. (b) Representative flow cytometry plots and quantification of B and T cell doublets (CD19$^+$CD8$^+$) after 72 h co-culture with unstimulated or activated (αCD3/CD28) sorted CD8$^+$ T cells. Dots represent 1-2 technical replicates from four donors across two independent experiments. (c) Representative flow cytometry plots showing the T cell phenotype of sorted CCR5$^+$ T$_{EM}$ cells and quantification of the T cell phenotype of sorted CCR5$^+$ and CCR5$^-$ T$_{EM}$ cells after 72 h co-culture with sorted CD19$^+$ B cells. Dots represent 1-2 technical replicates from four donors across two independent experiments. (d) B cell activation, plasmablast formation, atypical B cell formation and total B cell count after 72 h co-culture with unstimulated or activated (αCD3/CD28) sorted CD8$^+$ T cells. Dots represent 1-2 technical replicates from four donors across two independent experiments. *P* values were calculated using Dunn’s test and adjusted using Šídák's correction for multiple comparisons. *, **, ***, **** correspond to $P < 0.05$, $P < 0.01$, $P < 0.001$, and $P < 0.0001$, respectively.

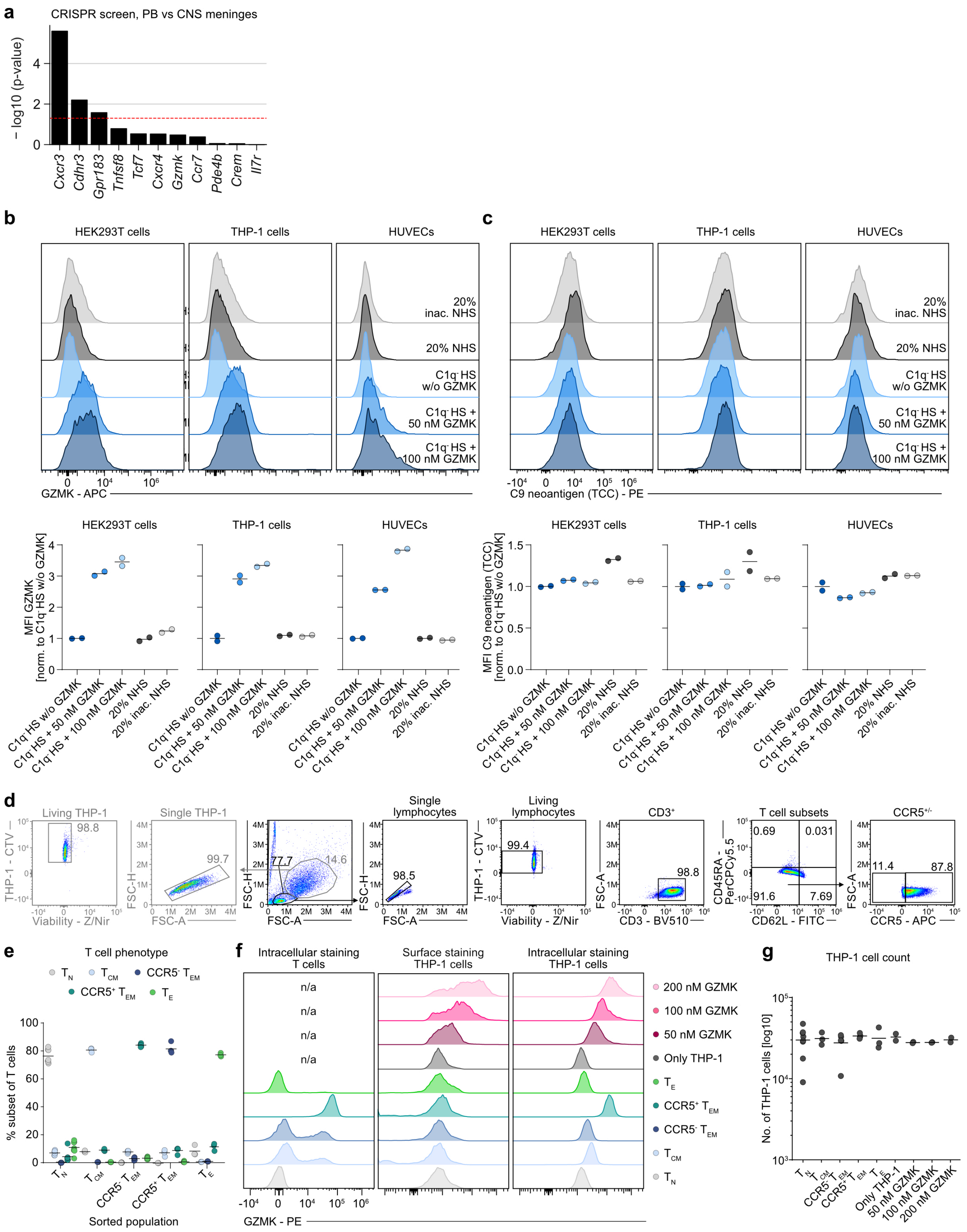
a
CRISPR screen, PB vs CNS meninges
− log10 (p-value)
Cxcr3
Cdhr3
Gpr183
Tnfsf8
Tcf7
Cxcr4
Gzmk
Ccr7
Pde4b
Crem
Il7r
b
HEK293T cells
THP-1 cells
HUVECs
20% inac. NHS
20% NHS
C1q⁻HS w/o GZMK
C1q⁻HS + 50 nM GZMK
C1q⁻HS + 100 nM GZMK
GZMK - APC
c
C9 neoantigen (TCC) - PE
MFI GZMK [norm. to C1q⁻HS w/o GZMK]
MFI C9 neoantigen (TCC) [norm. to C1q⁻HS w/o GZMK]
d
Living THP-1
Single THP-1
Single lymphocytes
Living lymphocytes
CD3⁺
T cell subsets
CCR5⁺/⁻
THP-1 - CTV
Viability - Z/Nir
FSC-H
FSC-A
CD3 - BV510
CD45RA - PerCPCy5.5
CD62L - FITC
CCR5 - APC
98.8
99.7
77.7
14.6
98.5
99.4
98.8
0.69
0.031
91.6
7.69
11.4
87.8
e
T cell phenotype
% subset of T cells
Sorted population
f
Intracellular staining T cells
Surface staining THP-1 cells
Intracellular staining THP-1 cells
n/a
200 nM GZMK
100 nM GZMK
50 nM GZMK
Only THP-1
GZMK - PE
g
THP-1 cell count
No. of THP-1 cells [log10]

**Supplementary figure 16: Supporting information on the extracellular and intracellular accumulation of GZMK in target cells.** (a) Observed effects of CRISPR-mediated knockouts *in vivo* reported by Kendirli et al. 2023 (PB vs. meninges) for genes overlapping with the CSF-biased score. (b) Representative flow cytometry plots and quantification of GZMK expression on HEK293T, THP-1 and HUVECs after 2 h at 37 °C. Dots represent technical replicates from one donor. (c) Representative flow cytometry plots and quantification of C9 neoantigen expression on HEK293T, THP-1 and HUVECs after 2 h at 37 °C. Dots represent a technical replicate from one donor. (d) Representative gates for the intracellular detection of GZMK in T cells, and the surface as well as intracellular detection of GZMK in THP-1 cells. (e) Percentage of color-indicated T cell subsets after 4 h of co-culture of sorted $CD8^+$ T cell subsets with THP-1 cells. Dots indicate 1-4 technical replicates per donor. (f) Fluorescence intensity histograms of anti-GZMK in surface and intracellular stains for the color-indicated sorted $CD8^+$ T cell subsets after 4 h of co-culture with THP-1 cells. Surface staining of $CD8^+$ T cells was not performed. (g) Log-normalized absolute THP-1 cell counts after 4 h of co-culture with sorted $CD8^+$ T cell subsets. Dots indicate 1-4 technical replicates per donor.
